# Identification of seven candidate structured noncoding RNA motifs in protists by comparative genomics

**DOI:** 10.64898/2026.11.14.765823

**Authors:** Yipeng Zhang, Qunying Liang, Tianshu Wang, Yixin Huang, Anqi Duan, Jiaojiao Wang, Wenlei Wang, Huan Yu, Binfen Chen, Zhilan Ying, Yin Jia, Chen Fu, Sanshu Li

**Affiliations:** Medical School, College of Materials Science and Engineering, Key Laboratory of Fujian Molecular Medicine, Key Laboratory of Xiamen Marine and Gene Drugs, Key Laboratory of Precision Medicine and Molecular Diagnosis of Fujian Universities, Institute of Genomics, Huaqiao University, Xiamen, 361021, China; Faculty of Life Science and Medicine, King’s College London, SE1 9RT, London, United Kingdom; Institutes of Biomedical Sciences, Fudan University, Shanghai 200032, China; Anyang No.1 Middle School, Anyang 456000, China; Fisheries College, Jimei University, Xiamen, 361021, China

**Author notes:** These authors contributed equally to this work (Yipeng Zhang, Qunying Liang, Tianshu Wang, Yixin Huang, Anqi Duan, Jiaojiao Wang).

**Keywords:** Comparative genomics, Structured noncoding RNAs, Protists, CMfinder, R-scape, Bioinformatics pipeline

## Abstract

Protists represent a highly diverse assemblage of predominantly unicellular eukaryotes, including protozoa, algae, and slime molds. Their essential roles in global ecosystems, along with their significance as human parasites, underscore the need for a deeper understanding of their conserved genetic elements. In this study, we employed an integrated computational approach combining CMfinder and R-scape to systematically identify and evaluate structured non-coding RNA (ncRNA) motifs across protist genomes. Through rigorous assessment and functional annotation, we report seven conserved structured ncRNAs that are likely functional RNA motifs involved in regulating the synthesis of the carbon fixation protein ribulose-1,5-bisphosphate carboxylase/oxygenase (RuBisCO), the light-harvesting and transfer protein phycocyanin, subunits of photosystems I and II, as well as mRNA splicing regulation. Notably, we also identified a structured ncRNA motif that may possess a dual function, capable of bidirectionally regulating the photosystem genes *psaI* and *psbJ* simultaneously. Our findings provide valuable insights into the diversity and functional significance of structured ncRNAs in microbial eukaryotes.

## Introduction

Protists are a diverse, mostly unicellular group of eukaryotic organisms that form a kind of “catch-all” kingdom (Protista) for eukaryotes that don’t fit neatly into the other three kingdoms (animals, plants, and fungi) (1–5). They share several key features: they are eukaryotic and mostly unicellular, though some form simple multicellular structures (e.g., seaweed, slime molds). Protists include organisms that resemble animals (protozoa – e.g., Amoeba, Paramecium), plants (algae – e.g., Euglena, Chlamydomonas, kelp), and fungi (slime molds, water molds). They also exhibit variable modes of nutrition, including photosynthetic (algae), heterotrophic (protozoa), or mixotrophic. Among these, protozoa constitute a motile and heterotrophic subgroup (3,5). Free-living protozoa function as phagotrophic organisms, playing a crucial role in regulating bacterial and microbial populations (1,2). In contrast, parasitic protozoans are often implicated in human diseases and present significant public health concerns, particularly in developing countries (3,4). These pathogens are responsible for millions of cases of morbidity and mortality each year (6). Unicellular algae, on the other hand, account for the bulk of carbon fixation in marine and freshwater ecosystems (7,8). Meanwhile, slime molds serve as key consumers of bacteria, fungi, and other decomposers, inhabiting organic-rich and forest soils (1,9).

Given the critical roles of protists in global ecosystems including carbon fixation, their basic foundation of the food chain, and their impact as human parasites—a deeper investigation into their conserved genetic elements is warranted. In particular, we are interested in structured noncoding RNAs (ncRNAs). Recent advances in noncoding RNA research in protists have primarily focused on long noncoding RNAs (lncRNAs), microRNAs (miRNAs), siRNAs, circRNAs, antisense RNAs, and other small regulatory RNAs. In 2007, comparative genomic analyses across *Plasmodium* species led to the identification of a putative snRNA promoter element (SNPE), along with several small nuclear and nucleolar RNAs (10). By 2010, transcriptomic studies in the human malaria parasite *Plasmodium falciparum* had experimentally identified 630 novel ncRNA candidates, comprising both snoRNAs and antisense RNAs (11). In 2014, a computational pipeline applied to published RNA-seq data from *P. falciparum* revealed 164 novel lncRNAs implicated in metabolic and biosynthetic processes, regulation of biological pathways, and host–pathogen interactions (12). In 2020, a study in *Cryptosporidium parvum* identified 396 putative novel lncRNAs, the majority of which were antisense transcripts located predominantly at the 3′ ends of sense mRNAs (13).

In contrast, microRNAs (miRNAs) are ∼22 nucleotide small noncoding RNAs that play key roles in regulating gene expression involved in development, differentiation, immune response, homeostasis, and cell-cell communication (14–16). Although miRNAs were initially characterized in multicellular eukaryotes, their presence in unicellular organisms was not established until 2007, with the discovery of miRNAs in the green alga *Chlamydomonas reinhardtii* (17,18). Subsequent studies identified miRNAs in two brown algal species (19,20), an additional green alga (21), two dinoflagellates (22,23), an ichthyosporean species (24), and a social amoeba (25). Most recently, in 2024, eight additional amoeba species were reported to encode miRNAs (26).

Certain non-coding RNAs (ncRNAs), including tRNAs, rRNAs, ribozymes (e.g., group I, group II, RNase P, hammerhead, hatchet, pistol, twister, and others) (27–34) and riboswitches (35), exhibit extensive base-pairing in their secondary structures and are therefore commonly referred to as structured ncRNAs (36–39). Computational methods for identifying structured ncRNAs often utilize tools such as QRNA (40), RNAz (41), CMfinder (42), and R-scape (43), which detect evolutionary signatures, particularly nucleotide covariation, rather than relying solely on sequence conservation. Among these, R-scape is notable for evaluating the statistical significance of detected covariation events.

Structured ncRNAs have been explored in *Plasmodium falciparum*. In 2008, the RNAz program was employed to predict conserved RNA structures within noncoding regions of the *P. falciparum* genome by comparing it with seven other *Plasmodium* species (44). This effort identified 668 conserved secondary structures; however, most alignments included only two sequences and lacked statistically significant covariation support. In 2021, some small RNAs resembling piRNAs were identified in *Trichomonas vaginalis*, where they play a role in regulating protein-coding genes and transposable elements (45). In 2022, a total of 79 novel intermediate-sized non-coding RNA candidates, including several snoRNAs, were identified in *Cryptosporidium parvum* using sRNA-seq and comparative genomics (46). Our group and others have utilized CMfinder and R-scape to discover novel numerous structured RNA candidates across diverse organisms, including bacteria (37,47–49), fungi (39,50), and metazoans—including humans (38,51,52). CMfinder (42) employs an expectation-maximization algorithm analogous to that of MEME (53) to identify potential RNA secondary structures from unaligned sequences using covariance models. To improve the accuracy of consensus secondary structure predictions, CMfinder also incorporates a thermodynamic model for base-pairing prediction (42). In contrast, R-scape uses a statistical approach that evaluates the significance of covariations in an alignment by applying a random model to detect phylogenetic signals (43). Together, the combined use of CMfinder and R-scape offers a robust strategy for predicting evolutionarily conserved RNA structures with high confidence.

In this study, we aim to expand the discovery of structured noncoding RNAs (ncRNAs) in protists—including protozoa, slime molds, and algae—by leveraging genomic data from 894 species available in public databases The broad phylogenetic diversity of protists offers a rich resource for detecting nucleotide covariations, thereby enhancing the identification of novel conserved RNA structures with functional significance.

The overall structural quality of each RNA motif is assessed based on three criteria: the identification of significant covariations (using R-scape), the detection of compatible mutations (via CMfinder and visualized with R2R), and structural complexity (determined by the number of hairpins and their organization). The best-structured ncRNA candidates were selected for further functional analyses.

These functional analyses include examining the genomic location of the motif along with its upstream and downstream genes, performing assays using RegRNA (54) and RBPmap (55). The goal is to determine whether these motifs function as riboswitches, ribozymes, or regulators of transcription or translation, among other possibilities. Notably, discoveries such as the long-distance regulation of alternative splicing by fungal TPP riboswitches (56), eukaryotic fluoride channel proteins (57), candidate ligands for the yjdF riboswitch (58), and three novel classes of natural ribozymes (47) were all based on functional analyses.

Through our pipeline, many structured ncRNA candidates in protists were discovered, and seven of the selected candidates are reported here. These ncRNAs exhibit a broad range of potential functions. Some may be involved in regulating the synthesis of carbon fixation proteins, light-harvesting and transfer proteins, photosystems I and II components, and various other proteins. Notably, we also identified a structured ncRNA motif that may function bidirectionally in regulating the photosystem genes *psaI* and *psaJ*. Together, our findings highlight the diversity and functional importance of structured ncRNAs in protists, a diverse group of predominantly unicellular eukaryotes.

## Materials and Methods

### Genome acquisition and species composition

The initial collection of protozoan genome sequences was obtained from NCBI RefSeq Release 201 (accessed via https://ftp.ncbi.nlm.nih.gov/refseq/release/protozoa) on July 15, 2020 (59). This dataset was subsequently updated to incorporate protozoan sequences available as of July 11, 2025 (RefSeq Release 230). The updated genome assembly served as the reference database for Infernal (Version 1.1.2) (60) to identify additional homologous sequences for all structured noncoding RNA candidates.

The protozoan genome collection comprises a total of 894 species (see Supplementary File 2 (Part I) for the complete list). The genomic coverage extends beyond typical protozoans—such as amebae, flagellates, ciliates, and parasitic sporozoans—to also encompass algae, reflecting the broad taxonomic scope of protists. Accordingly, the noncoding RNA motifs identified in this study originate from a diverse array of protist lineages.

For reference, the following is an alphabetically ordered subset of species included in the analysis: *Acanthamoeba castellanii* (a small free-living amoeba commonly found in soil and freshwater), *Babesia bigemina* (an apicomplexan parasite that infects cattle and causes bovine babesiosis), *Biddulphia tridens* (a diatom species),

*Cafeteria roenbergensis* (a heterotrophic marine nanoflagellate), *Choreocolax polysiphoniae* (a red algal parasite), *Cryptosporidium bovis* (one of the Cryptosporidium species infecting cattle), *Haemoproteus belopolskyi* (a cosmopolitan haemosporidian parasite), *Naegleria fowleri* (a pathogenic amoeba), *Ochromonas danica* (a mixotrophic flagellate), *Pachymeniopsis lanceolata* (a red alga), *Phytophthora agathidicida* (a pathogen of New Zealand kauri trees), *Rhizochromulina marina* (a marine alga), *Saccharina diabolica* (a brown alga), *Sargassum fulvellum* (a brown alga), *Taenioma perpusillum* (a species of macroalgae), *Ulnaria ulna* (a diatom), *Vacuolaria virescens* (a green alga), *Wildemania schizophylla* (an algal species), *Xiphosiphonia pinnulata* (an alga), and *Yamadaella caenomyce* (a red alga).

### Identification of candidate RNA motifs

The analytical pipeline used in this study (Fig. 1) was adapted from previously established methods for identifying structured ncRNAs in fungi (39) and humans (38). Briefly, intergenic regions (IGRs) and introns were extracted from all annotated genes across the genomes using a custom PERL script (Supplementary File 3). These sequences were compared to one another using NCBI BLAST (version 2.2.25) (61) with the following parameters: blastn-word_size 7-evalue 1e-6-perc_identity 75.0. Similar sequences were grouped automatically by BLAST and an additional PERL script to form preliminary clusters (Supplementary File 4). Each cluster was first filtered by performing a BLAST search against the Rfam database (version 14.1) (62) with parameters:-word_size 10-evalue 1e-10-perc_identity 90.0 in order to exclude known RNAs. The resulting candidate clusters were further filtered with RNAcode (version 0.3) (63) using a P-value cutoff of 0.001 to remove clusters displaying protein-coding potential. The remaining clusters were analyzed with CMfinder (version 0.4.1.9) (42) to predict RNA secondary structures based on covariation signals among sequences. Clusters exhibiting little covariation or containing fewer than three sequences were discarded. Finally, Infernal (60) was used to search for additional homologs using both sequence and structure similarity and to refine the RNA secondary structure when more examples were identified.

**Fig. 1.**
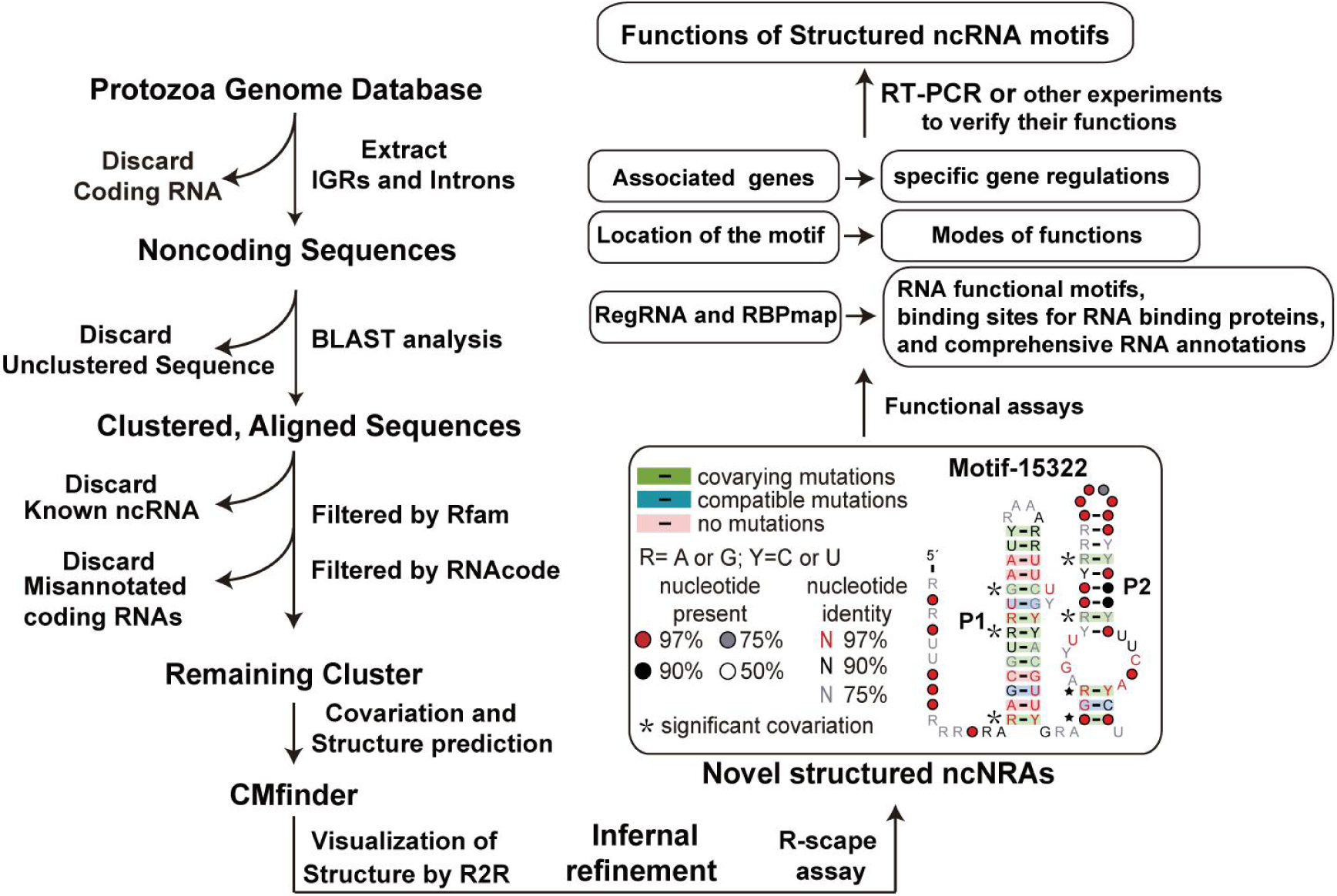
Workflow for the identification of novel structured noncoding RNAs in protists. Detailed procedures are provided in the Materials and Methods section. Covariations refer to paired nucleotide substitutions that maintain canonical base pairing (e.g., A–U → G–C). Compatible mutations are defined as single-nucleotide changes that retain the potential for base pairing (e.g., A–U → G–U). A star indicates a base pair showing statistically significant covariation as assessed by R-scape.

### Assessing the novelty of motifs

To assess the novelty of the motifs, we searched the Rfam database (https://rfam.org/) (62) using Infernal to identify and thereby remove any motifs with homologous non-coding RNAs. Subsequently, we used the sequences of the remaining motifs to search against RNAcentral (64), allowing us to exclude known RNA motifs.

### Algae cultures

The thallus (gametophytes) of *P. haitanensis* were cultured in natural seawater under 22℃ or 28℃ at light for 12 days before harvesting the thallus(65). The samples were stored at-80℃ before RNA extraction.

### RNA extraction

RNA extraction was performed by using the TIANGEN RNAprep Kit (TIANGEN Biotech Co., LTD) with 80 mg of thallus sample and following the manufacturer’s protocol.

### Reverse Transcription PCR (RT-PCR)

RT-PCR was performed as previously described(38). Table S1 (in the Supplementary file 1) lists the primer sequences and the expected sizes of the RT-PCR products. During RT-PCR, a negative control, in which the RT reaction lacked reverse transcriptase, was included to ensure there is no DNA contamination. The gene ubiquitin-conjugating enzyme (*UBC*) from *P. haitanensis* was used for internal control (66) (Fig. S1 in the Supplementary file 1).

### Quantitative Real-Time PCR (qPCR)

qPCR was performed on the Agilent Mx3000P real-time PCR system using the TransStart® Top Green qPCR SuperMix kit according to the manufacturer’s instructions (Transgen Biotech, Beijing, China). Table S1 lists the primer sequences and the expected sizes of the qPCR products. qPCR cycles include Segment 1: 50℃ 120 s; Segment 2: 95℃ 120 s; Segment 3: 95℃ 15 s, 54℃ 15s, 72℃ 15s(end point)for 42 cycles; Segment 4: 95℃ 60s, 55℃ 30s (all point), 95℃ 30s.

## Results

### Structured ncRNA candidate identification

We obtained genomic data for protist species from NCBI, encompassing a broad taxonomic spectrum including amoebae, diatoms, nanoflagellates, red algae, brown algae, and other groups. Our computational pipeline for structured ncRNA identification (Fig. 1) was adapted from established methods previously applied to fungi (39) and humans (38). Detailed procedures for identifying structured ncRNA candidates are provided in the Materials and Methods section. In brief, noncoding regions were extracted from protist genomes using a custom PERL script. Sequence-similar regions were then clustered using BLAST. These clusters were subsequently filtered to remove those matching known ncRNAs in Rfam (version 14.1) (62), followed by the elimination of clusters with significant protein-coding potential as predicted by RNAcode (63). The remaining candidate clusters were analyzed with Cmfinder to detect nucleotide covariation patterns (42). Conserved sequences and substructures for each ncRNA candidate were iteratively refined using Infernal (60). Finally, statistical support for the conserved secondary structures was assessed using R-scape (43).

Based on the predicted structure—taking into account covariations (assessed by R-scape), compatible mutations, the number of sequence representatives, and structural complexity—we selected several candidate RNA motifs for further analysis. We then used RBPmap(55) and RegRNA (Version 3.0) (54) to identify conserved binding sites for RNA-binding proteins and to detect regulatory elements within the RNA, respectively. This allowed us to infer the type of genetic element these structured non-coding RNA motifs might represent, such as ribozymes, riboswitches, alternative splicing regulators, or translational regulators. Below, we describe the most promising candidate motifs that are likely to represent functional ncRNAs. The corresponding alignments for these high-ranking motifs in Stockholm format are provided in Supplemental Files 3-11. The motif is typically assigned a simple identifier, such as Motif_15322, where the numeric component (e.g., 15322) is automatically generated by the analysis pipeline during motif discovery. If, however, the genes associated with the motif are known, its name is instead constructed by combining a relevant gene symbol with a numeric suffix, for example, RbcL_8943.

### The RbcL_8943 RNA motif

This motif comprises six distinct representative sequences, each derived from the chloroplast genome of one of six Porphyra species, respectively: *P. pulchra*, *P. umbilicalis*, *P. haitanensis*, *P. yezoensis*, *P. purpurea*, and *P. perforata*. The consensus sequence and secondary structure model (Fig. 2A) reveal that this RNA motif contains three stem-loop structures (designated P1 to P3). The P1 stem contains one compensatory mutation, and the P2 stem contains six covarying mutations, one of which is statistically significant evaluated by R-scape (43). These features suggest that this RNA motif likely forms stem-loop structures to fulfill its functional role.

**Fig. 2.**
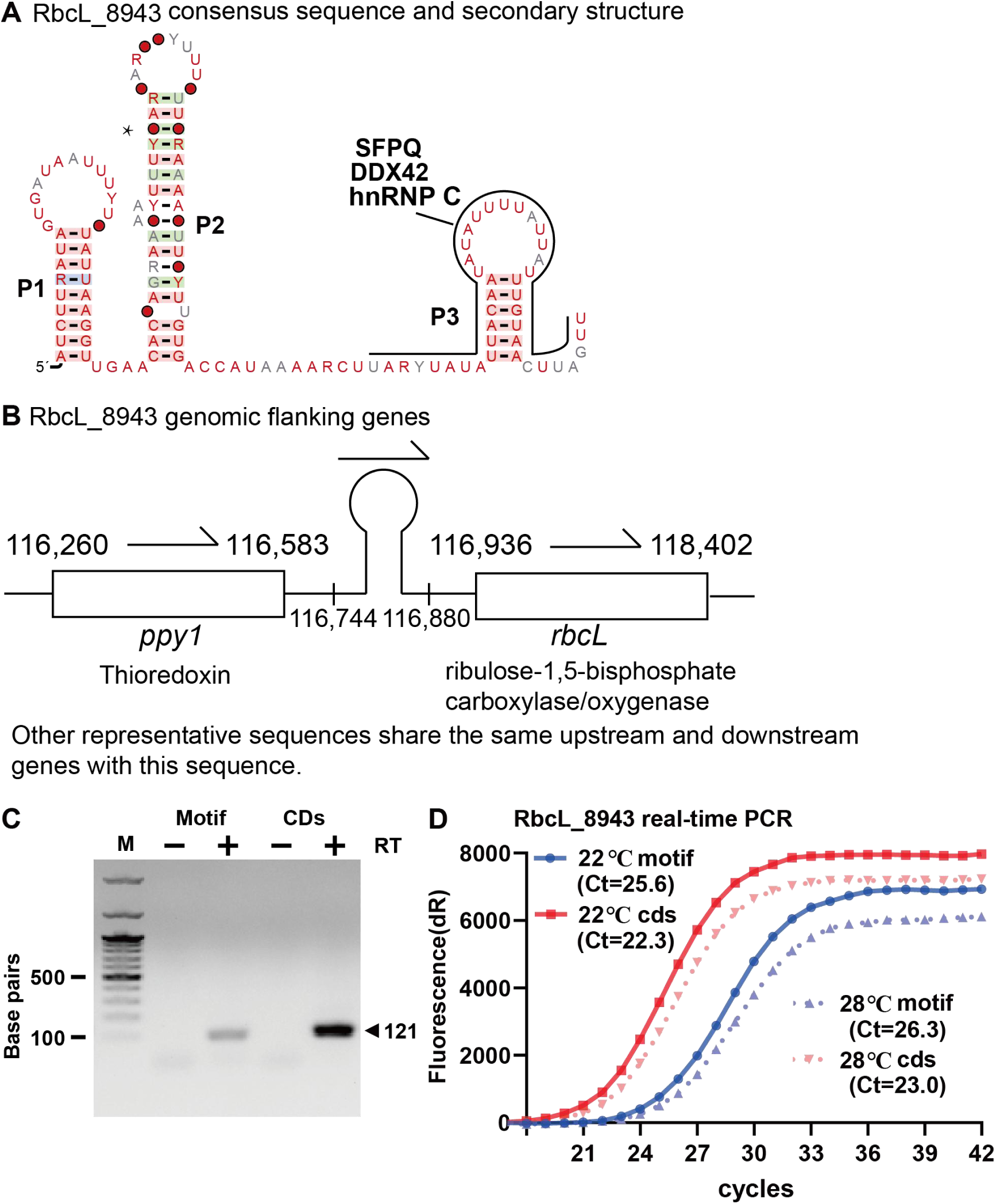
The RbcL_8943 RNA motif and its expression. **A.** Consensus sequence and secondary structure of the RbcL_8943 motif. The RNA binding sites for the RNA-binding proteins are shown around the P3 stem. Other annotations are as listed in Fig.1. **B.** Genomic context depicting the RNA motif (shown as a hairpin) and its flanking genes (boxed), along with their respective genomic positions (not drawn to scale). Gene orientations are indicated by arrows. Agarose gel shows the separation of RT-PCR products generated by using primers specific for the transcripts as designated. M designates double-stranded DNA size markers, and lanes containing PCR products generated with (+) or without (–) the use of reverse transcriptase (RT) are identified. **C.** RT-PCR analysis of the motif and the coding sequence (CDS) of its adjacent gene. **D.** qPCR for the motif RNA for algae cultured under temperature stress. Ct represents threshold cycles to observe the exponential increase of the amplified PCR product. The experiment was conducted twice, and consistent results were obtained both times.

Further analysis of each sequence within the motif revealed its close proximity (<50 nucleotides) to the ribosomal binding site (RBS) (Fig. 2B), potentially regulating downstream genes by modulating ribosome binding. The downstream genes associated with this motif are all photosynthesis-related *rbcL* genes, which encode ribulose-1,5-bisphosphate carboxylase/oxygenase (Rubisco) – a key enzyme governing carbon assimilation rates in photosynthesis and photorespiration in plants. Magnesium ions are crucial participants in Rubisco-catalyzed reactions, with their concentration directly affecting carbon fixation efficiency. The CO2 concentration at the catalytic site is another critical factor influencing carbon fixation. Additionally, chloroplasts contain various other metabolites (primarily phosphorylated compounds) that may interfere with or interact during the activation process (67). Therefore, Motif_8943 may utilize its formed RNA structure to participate in the regulatory processes of these small molecules.

In addition, RBPmap (55) predicts that the RNA motif contains a small hairpin at its 3’ end that serves as the binding site for several RNA-binding proteins, including hnRNP C (heterogeneous nuclear ribonucleoprotein C), SFPQ (splicing factor, proline-and glutamine-rich), and DDX42 (an ATP-dependent RNA helicase) (Fig. 2A). These proteins are involved in RNA structure recognition, spliceosome assembly, and RNA remodeling(68–70).

To determine whether the RbcL_8943 RNA motif is expressed in cells, whether its expression level is comparable to that of housekeeping genes, and whether its expression is influenced by temperature, we performed Reverse transcription-PCR (RT-PCR) and quantitative real-time PCR (qPCR) using specific primers (Table S1 in Supplementary File 1)

RT-PCR analysis demonstrated that primers specific to the RbcL_8943 motif and its flanking CDS successfully amplified the expected products (Fig. 2C; see Supplementary File 2 (Part III) for the uncropped gel image), confirming the expression of both the motif and the corresponding CDS RNAs in the cells. The *ubc* gene (encoding ubiquitin-conjugating enzyme) (66) was used as an internal control (Fig. S1 in Supplementary File 1).

The expression level of the RbcL_8943 RNA motif was further quantified using qPCR. Results indicated that the abundance of the RbcL_8943 motif was lower than that of its associated coding sequence RNA. Additionally, both the motif and the RbcL CDS RNA showed moderately higher expression at 22°C compared to 28°C (Fig. 2D). Furthermore, expression levels of the RbcL gene transcript exceeded those of the *ubc* housekeeping gene (Fig. 2D and Fig. S1).

### The SRP_like_8976 RNA motif

This motif comprises 124 representative sequences, primarily derived from the chloroplast genomes of diverse algal species. These include brown algae of the genus *Porphyra* (e.g., *P. perforata*, *P. haitanensis*, *P. yezoensis*, *P. pulchra*, *P. purpurea*) and red algal species such as *Sheathia arcuata*, *Kumanoa americana*, *Thorea hispida*, *Schimmelmannia schousboei*, and *Plocamium cartilagineum*, among others. The consensus sequence and secondary structure model (Fig. 3A) reveal a long hairpin structure containing internal loops, one of which forms a large 38-nucleotide loop. The stem region displays 13 covarying mutations, seven of which are statistically significant according to R-scape analysis. These features suggest that the motif likely functions through hairpin formation.

**Fig. 3.**
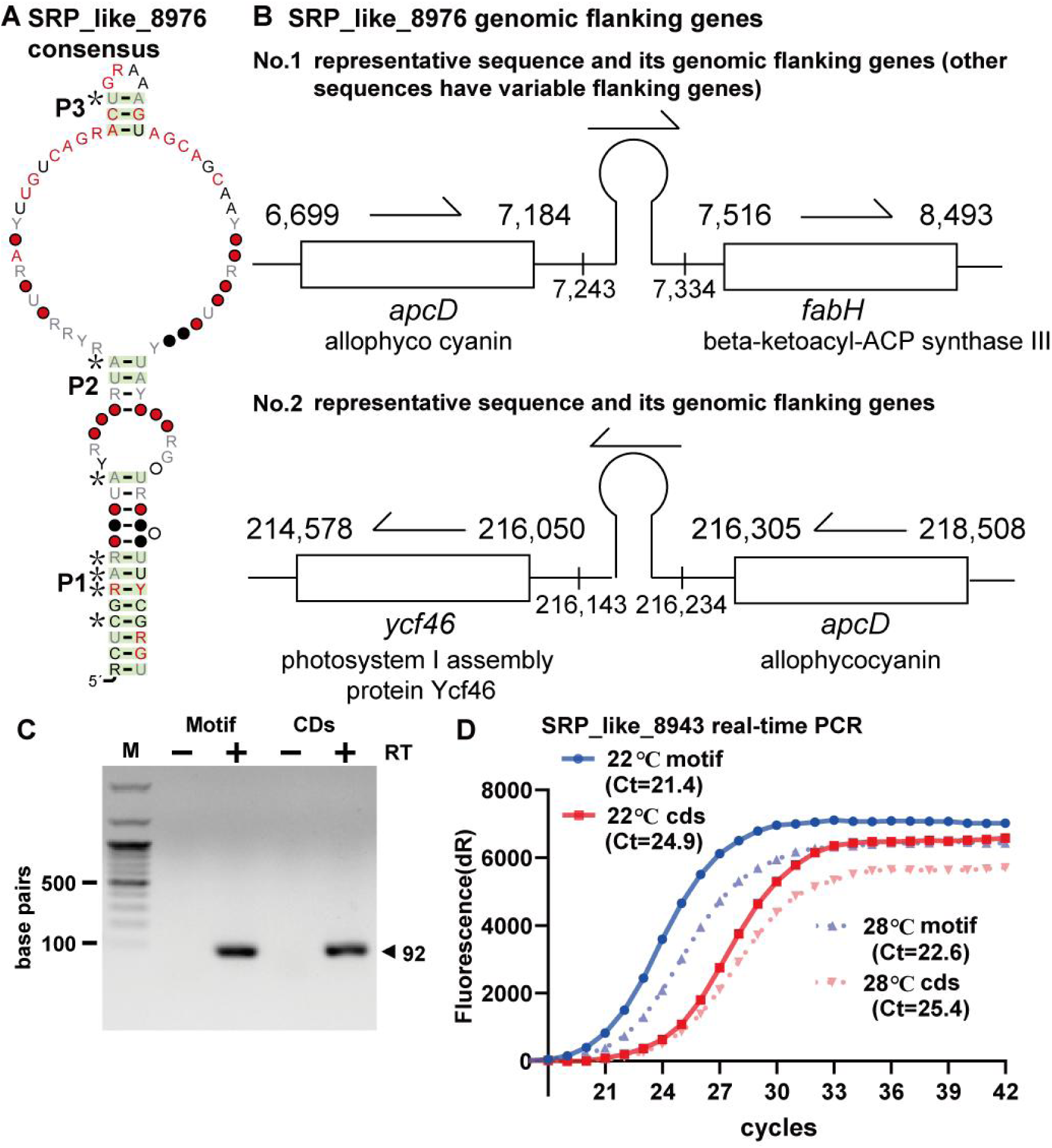
The SRP_like_8976 RNA motif and its expression. **A.** Consensus sequence and secondary structure of the SRP_like_8976 motif. **B.** Representative genomic context of the motif, showing the most common flanking genes. **C.** RT-PCR analysis of the motif RNA and the CDS region of an adjacent gene. **D.** qPCR quantification of motif expression in algae subjected to temperature stress. Other annotations are consistent with those in Fig. 2.

In most representative sequences, the motif is located between the upstream gene *apcD* (encoding allophycocyanin) and the downstream gene *fabH* (encoding beta-ketoacyl-ACP synthase III) (Fig. 3B). In some sequences, the downstream gene is *ycf46*, while *apcD*remains upstream. This genomic context implies a potential role in regulating *apcD*function, provided the RNA motif possesses regulatory activity.

However, this motif most likely functions as an independent structural RNA. One representative sequence shows similarity to the signal recognition particle (SRP) RNA found in algal chloroplasts. SRP RNAs are highly variable in sequence and structure across species, complicating their prediction(71). Confirmation of this similarity would demonstrate the efficacy of our approach in identifying additional SRP like RNAs.

RT-PCR analysis using primers targeting Motif_8976 and its flanking coding regions yielded products of the expected sizes (Fig. 3C; see Supplementary File 2 (Part III) for uncropped gel images), confirming transcription of both the motif and adjacent CDS regions. Expression analysis revealed that SRP_like_8976 RNA levels exceeded those of the neighboring CDS at both 22 °C and 28 °C (Fig. 3D). Additionally, both the RNA motif and the adjacent CDS exhibited modestly higher expression at 22 °C than at 28 °C (Fig. 3D).

### The PsaI_8813 RNA motif

This motif consists of 54 distinct representative sequences, 12 of which originate from the chloroplast genomes of diverse brown algae (*Phaeophyta*) species. These include representatives from *Sargassum thunbergii*, *Sargassum horneri*, *Saccharina japonica*, *Coccophora langsdorfii*, *Costaria costata*, and *Undaria pinnatifida*. Consensus sequence and secondary structure analysis (Fig. 4A) indicate that this RNA motif adopts a hairpin structure characterized by more than 20 consecutive canonical Watson–Crick base pairs. Among these, 18 are covarying mutations, 16 of which are statistically significant, suggesting that the structural integrity is critical for RNA function. Among these sequences, 12 originate from chloroplast genomes and occupy the same genomic locus (Fig. 4B), with their 5′ upstream gene encoding the psaI subunit of photosystem I or psbJ subunit of photosystem II on the neighbor gene with opposite direction. The organization on the genome indicates that both *psbJ* and *psaI* gene can use it to stabilize the mRNA, especially the RNA motif is at the end of the stop codon of both *psaI* and *psbJ* genes (Fig. 4C). The most notable feature of this RNA motif is its ability to form over 20 consecutive canonical Watson–Crick base pairs, allowing it to function in both the sense and antisense directions.

**Fig. 4.**
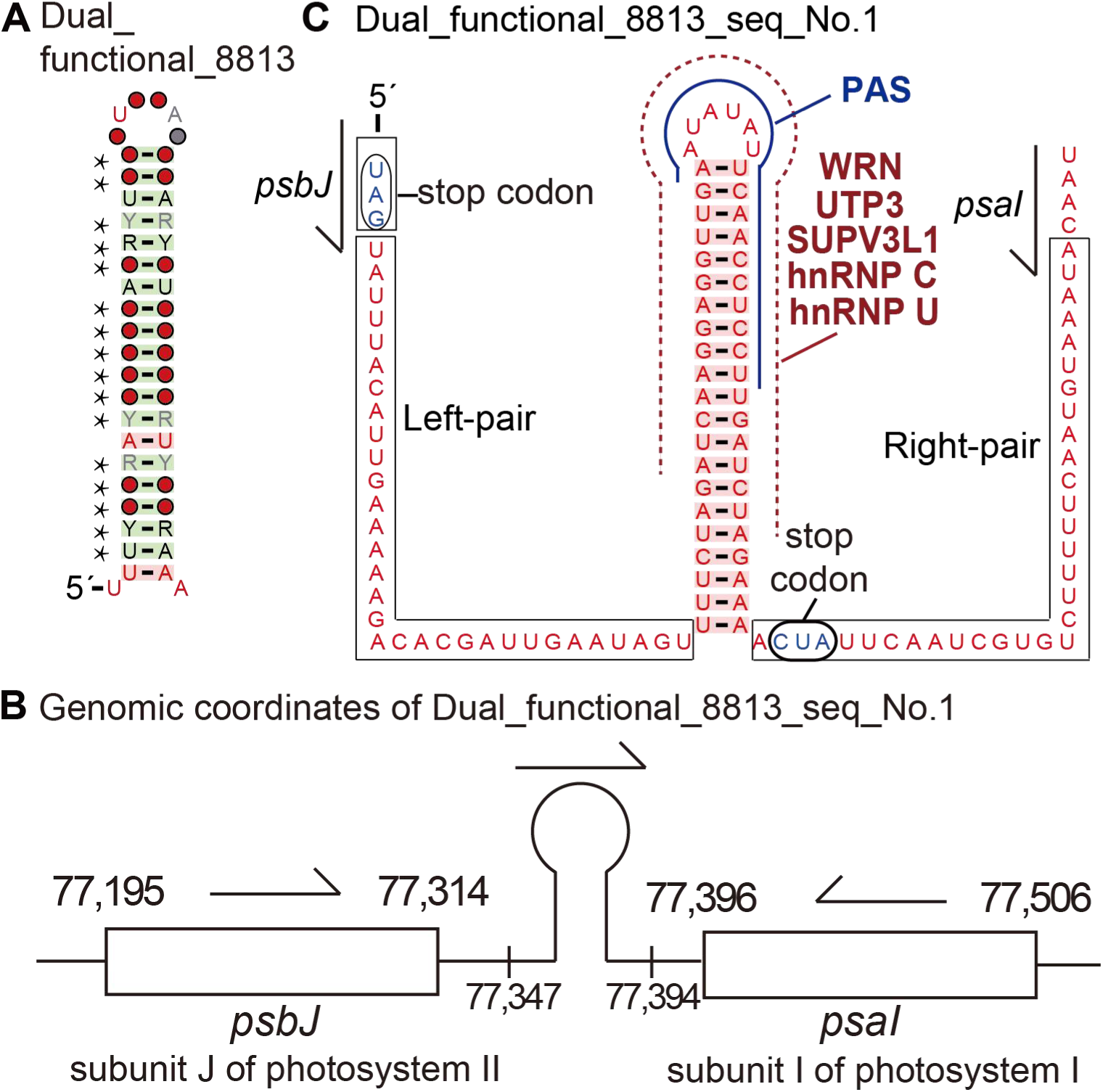
Structural and genomic features of the PsaI_8813 RNA motif. **A.** Consensus sequence and predicted secondary structure of the PsaI_8813 motif. The RBP binding sites and PAS sequences are shown around the long stem. **B.** Genomic organization of the conserved RNA motif. The representative sequences are shown with its flanking genes. **C.**Extended base-pairing interactions (boxed regions) within the RNA motif. Thirty-two additional base pairs were identified in the flanking regions of the motif. The RNA motif appears to form a short hairpin immediately after the stop codon of the *psaI* gene on the complementary strand (indicated by *psaI* and an arrow), while forming a longer hairpin for *psbJ* after its stop codon (on the sense strand, indicated by *psbJ* and an arrow). Arrows indicate the transcriptional directions of the two oppositely oriented genes. Other annotations are as described for Fig. 2.

In addition, RegRNA(54) predicts the formation of a long stem structure, consistent with our CMfinder results (Fig. 4A). Intriguingly, this long-stem hairpin contains a polyadenylation signal (PAS) of 30 nucleotides (nt) located on the right side of the stem. On the opposite strand, the stem also harbors an identical PAS. RBPmap predicts that the hairpin regions serve as binding sites for several RNA-binding proteins, including heterogeneous nuclear ribonucleoprotein C (hnRNP C), hnRNP U, SUPV3L1, WRN, and UTP3. These proteins are involved in diverse functions such as RNA splicing, normal linear mRNA production, mRNA stabilization and half-life extension, degradation of unwanted or aberrant RNA transcripts, RNA 3’-end processing, and ribosome assembly(72).

### The CpcB_4508 RNA motif

This motif consists of seven representative sequences derived from the chloroplast genomes of seven *Porphyra* species: *P. pulchra*, *P. umbilicalis*, *P. haitanensis*, *P. yezoensis*, *P. purpurea*, *P. schizophylla*, and *P. perforata*. The consensus sequence and secondary structure model (Fig. 5A) reveal that the RNA motif folds into two stem-loop structures (P1 and P2). The stems of P1 and P2 display two covarying mutations, one of which represents a significant covariation, along with four compensatory mutations. These findings support the structural integrity of the motif, which is likely essential for its function.

**Fig. 5.**
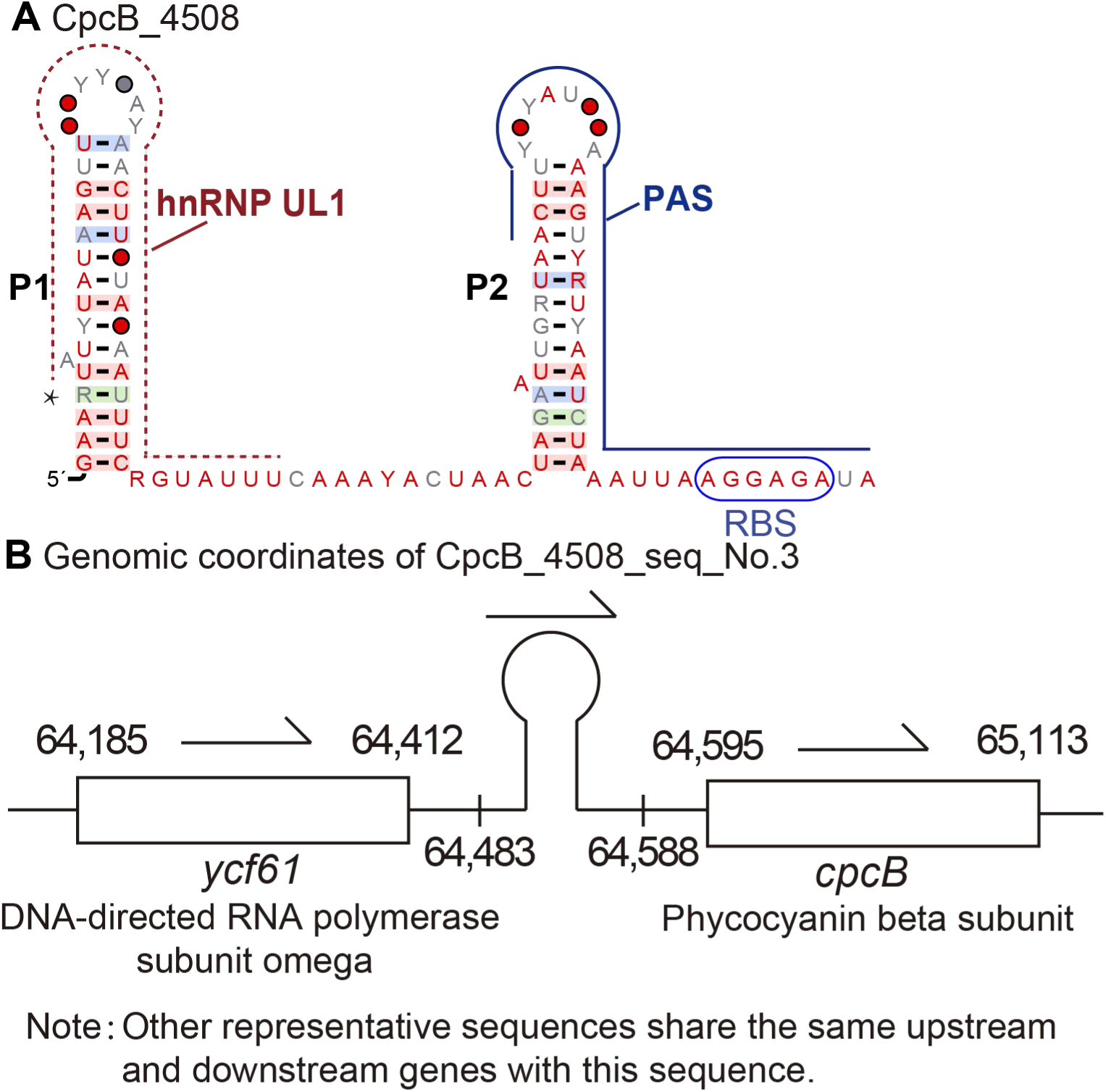
Structural and genomic features of the CpcB_4508 RNA motif. **A.** Consensus sequence and predicted secondary structure of the CpcB_4508 motif. The RBP (hnRNP UL1) binding sites and the PAS sequence are annotated. **B.** Genomic context of the conserved RNA motif, showing its relative position to neighboring genes and the ribosomal binding site (RBS). Other annotations follow the conventions established in Fig. 2.

In addition, RegRNA (54) reveals that the RNA motif contains a polyadenylation signal (PAS) at its 3’ end (Fig. 5A). This PAS forms a small hairpin as part of the P2 stem and extends toward the ribosome binding site (RBS). RBPmap shows that the P1 stem often serves as a binding site for RNA-binding proteins such as hnRNP UL1 (Heterogeneous Nuclear Ribonucleoprotein U-like 1) and STAU2 (Staufen Double-Stranded RNA Binding Protein 2). hnRNP UL1 primarily acts as a transcriptional regulator and DNA damage response factor in the nucleus(73), while STAU2 mainly functions in the transport and localization of mRNA within neuronal cells(74).

Notably, these sequences are conserved in their genomic positioning: the *ycf61* gene (encoding DNA-directed RNA polymerase subunit omega) is located ∼70 bp upstream of the 5′ end, while the *cpcB* gene (encoding the phycocyanin beta subunit) resides 7–8 bp downstream of the 3′ end. Further examination revealed that the motif lies adjacent to the ribosomal binding site (RBS) of *cpcB*, implying a potential regulatory role in ribosome recruitment and translational control (Fig. 5B).

The *cpcB* promoter, native to cyanobacteria, drives the expression of phycocyanin-encoding mRNA and is known to be repressed under high light while modulating activity in response to CO₂ levels (75,76). Given its positioning and structural features (Fig. 5B), the RNA motif may contribute to translational regulation of *cpcB* in response to environmental cues such as light fluctuations or varying CO₂ concentrations or secondary messengers, behaving like a riboswitch (35).

### The PsbH_5095 RNA motif

This motif comprises 11 distinct representative sequences derived from the genomes of four different species within the genus *Nannochloropsis* (a genus of *Eustigmatophyceae* microalgae in the phylum of *Heterokontophyta*)(77). These species include *N. oculata*, *N. granulata*, *N. limnetica*, *N. salina*, *N. gaditana* and *N. oceanica*. The consensus sequence and secondary structure model (Fig. 6A) indicate that this RNA motif comprises two stem-loop structures. The first stem-loop displays four covariations, three of which are statistically significant, along with one compensatory mutation. The second stem-loop contains two covariations and two compensatory mutations. These structural features suggest that the motif functions as a structured RNA element.

**Fig. 6.**
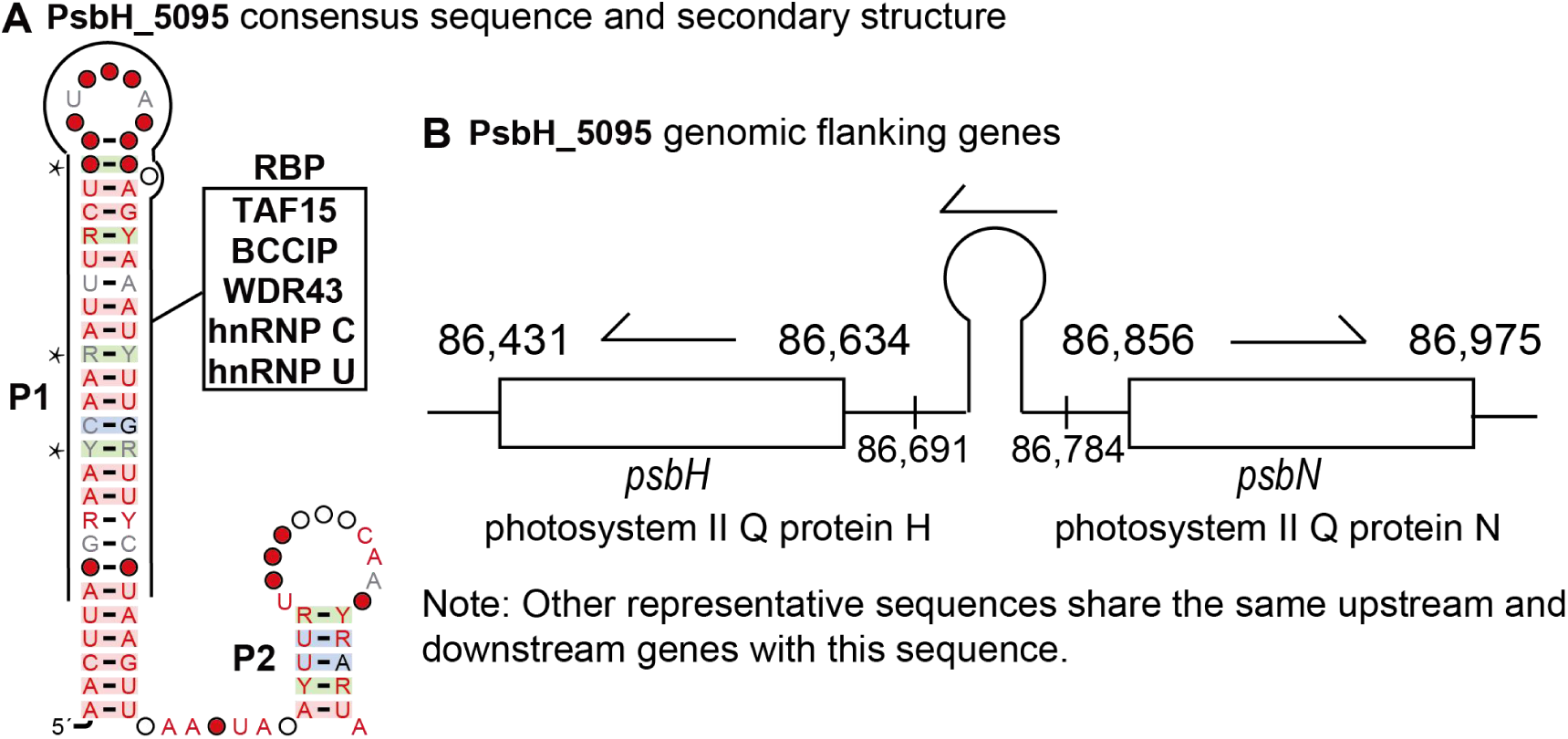
Structural and genomic features of the PsbH_5095 RNA motif. A. Consensus sequence and predicted secondary structure of the PsbH_5095 motif. B. Genomic organization of the conserved RNA motif. Other annotations are as described for Fig. 2.

In addition, RegRNA predicts the formation of a long stem structure (P1), which is consistent with our CMfinder predictions. RBPmap indicates that these long-stem hairpin regions serve as binding sites for several RNA-binding proteins, including TAF15 (TATA-box binding protein associated factor 15), BCCIP (BRCA2 and CDKN1A-interacting protein), WDR43 (WD Repeat Domain 43), hnRNP C and hnRNP U. These proteins are involved in diverse functions, such as RNA splicing, production of normal linear mRNA, mRNA stabilization, poly(A) RNA binding, maturation of the 18S rRNA, and releasing paused polymerase(72).

The gene most closely associated with this motif is the downstream *psbH* gene, which encodes a phosphorylated protein (PsbH) of photosystem II (PSII) (Fig. 6B). The immediately adjacent upstream gene is *psbN*, encoding the photosystem II protein N. The PsbH protein is a core component of the PSII complex, responsible for electron transfer between QA and QB. Mutants of PsbH exhibit heightened sensitivity to photoinhibition under high-light conditions(78). In Arabidopsis thaliana, PsbH has been demonstrated to be essential for the accumulation of CP47, a component of the internal antenna complex that channels energy from the external antenna to the reaction center (78).

Additionally, studies suggest that marine algae evolved from terrestrial ancestors. The *psbH* gene contains positive selection sites, which may have played a critical role in the adaptation of seagrasses to marine environments (78). Therefore, this motif may function by regulating *psbH* gene expression, thereby modulating the impact of high light intensity and elevated temperature on photosynthesis in cyanobacteria.

### The Motif_11540 RNA

This motif comprises 50 distinct representative sequences derived from the chloroplast genomes of various algae including *Porphyridium sordidum*, *Porphyridium purpureum*, *Bulboplastis apyrenoidosa*, *Dixoniella grisea*, and *Glaucosphaera vacuolata*. The consensus sequence and secondary structure model (Fig. 7A) reveal that this RNA motif contains multiple stem-loop structures. Stems P1 to P5 exhibit 16 covarying mutations, 13 of which show significant covariation, suggesting that this RNA motif likely functions through the formation of a complex structure.

**Fig. 7.**
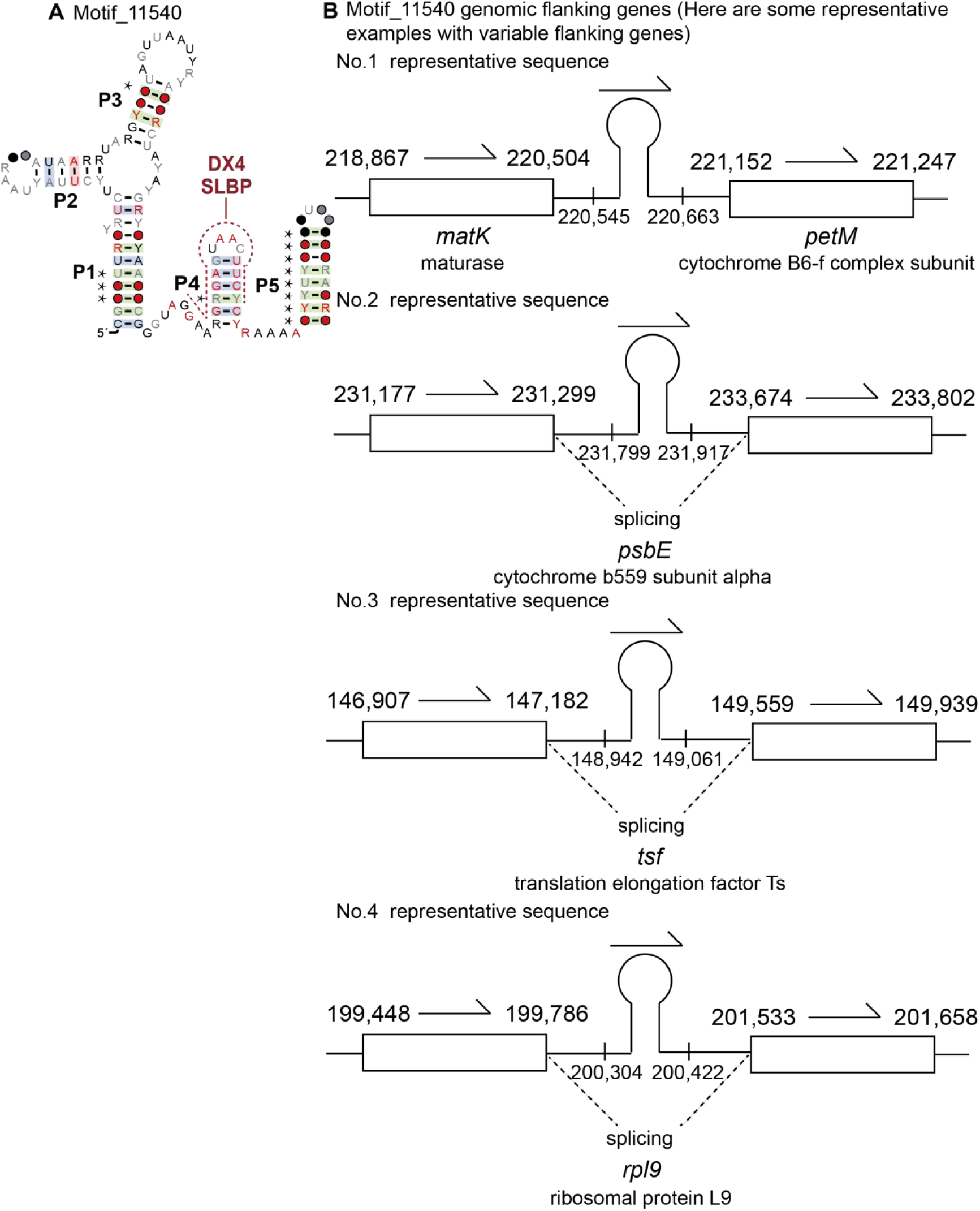
Structural and genomic features of the Motif_11540 RNA. **A.** Consensus sequence and predicted secondary structure of the Motif_11540 RNA. The RBP (SLBP and DX4) binding sites are annotated around the P4 stem. **B.** Genomic organization of the conserved RNA motif. Other annotations are as described for Fig. 2.

RBPmap predicts that the right side of the P4 stem contains binding sites for the RNA-binding proteins SLBP (Stem-Loop Binding Protein) and DX4 (DEAD-box Helicase 4 / VASA). SLBP plays a key role in mRNA transport and translation(79), whereas DDX4 utilizes the energy from ATP to bind and unwind RNA molecules, thereby remodeling RNA secondary structures or modulating RNA-protein interactions(80).

This motif is associated with variable genes (Fig. 7B). Examples of upstream and downstream genes include maturase-encoding *matK*, *petM* (encoding cytochrome b6 subunit), *psbB* (encoding photosystem II CP37 protein), *psaM* (encoding photosystem I subunit XII), and *gltB* (encoding glutamate synthase). Additionally, some of the representatives are located within introns of genes such as *psbE* (encoding cytochrome b559 alpha subunit), *tsf* (encoding translation elongation factor Ts) and *rpl9* (encoding ribosomal protein).

Here list functions of some associated genes. The *gltB* gene encodes glutamate synthase. Dr. Breaker’s lab and other research groups have discovered riboswitches capable of sensing glutamine and regulating gene expression (81,82). Given that this motif resides within the *gltB* intron, it may also respond to metabolite concentration changes. PsaK and PsaM are small, integral membrane proteins associated with the photosystem I complex in cyanobacteria (83). While both *psaK1* and *psaK2* genes are expressed in cyanobacteria, their absence only slightly reduces photosystem I electron transport. In contrast, the lack of PsaM decreases trimer recovery by 75%, indicating its essential role in stabilizing photosystem I trimers (83). In addition, the *psbB* gene, encoding a photosystem II core protein, is highly upregulated in heat-stressed plants treated with nitric oxide and proline (84). Given that Motif_11540 RNA is associated with these genes, it may undergo structural changes in response to variations in light, temperature, or metabolite levels.

### The Motif_15322 RNA

This RNA motif comprises 24 distinct representative sequences derived from the chloroplast genomes of various algae, encompassing 12 species: Colacium mucronatum, *Cryptoglena skujai*, Discoplastis spathirhyncha, *Euglena archaeoplastidiata*, *Euglena anabaena*, *Euglena clara*, Euglena deses, *Euglena hiemalis*, Flexiglena variabilis, *Lepocinclis steinii*, *Lepocinclis tripteris*, and Strombomonas costata. The consensus sequence and secondary structure model (Fig. S2A) reveal that this RNA motif contains two hairpin structures, P1 and P2, with 12 covarying and three compensatory mutations in their stems. In addition, the P1 stem contains a conserved two-nucleotide (UY) bulge which may be important for a functional structured noncoding RNA.

In addition, RBPmap reveals that the conserved sequences flanking both sides of the P1 stem serve as binding sites for splicing factors or RNA-binding proteins (Fig. S2A). Specifically, the left-side sequences are recognized by splicing factors such as CFTR, AGER, and TPM2, as well as RNA-binding proteins including FUS and SFPQ. The right-side sequences are bound by splicing factors like SRC and FAS, and RNA-binding proteins such as PABPN1. Moreover, the P1 stem itself may contain polyadenylation signals or binding sites for RNA-binding proteins, including WRN (Werner syndrome protein) and CPEB4 (cytoplasmic polyadenylation element-binding protein 4). WRN is involved in regulating small RNA biogenesis(85), whereas CPEB4 binds to the cytoplasmic polyadenylation element (CPE)—a conserved UUUUUAU sequence in mRNA 3′-UTRs—to control protein synthesis(86).

Motif_15322 may be closely linked to the *rbcL* gene (encoding ribulose-1,5-bisphosphate carboxylase/oxygenase) due to its proximity to the gene’s start codon (Fig. S2B). Ribulose-1,5-bisphosphate carboxylase/oxygenase (Rubisco) catalyzes the incorporation of inorganic CO₂ into organic molecules of life, such as nucleotides (67). Many aspects of Rubisco, including its complex biosynthesis and multi-step catalytic reactions, are tightly regulated by factors such as light, cellular metabolites, and molecular chaperones(67). *Motif_15322* may participate in these regulatory mechanisms through structural changes, behaving like a riboswitch(35,58). Additionally, it is located within the introns of several photosynthetic system genes, such as *psbC*, *psbD*, and *psaB*.

Thus, this RNA motif may also be involved in the regulation of photosynthesis in response to light, temperature, nutrients, and other environmental factors. These hypotheses, however, remain to be validated through experimental investigation.

## Discussion

### Discovery of noncoding structured RNA in protists

Protists comprise a diverse group of single-celled eukaryotes, including protozoa, algae, and slime molds (1–5). Although some structured small noncoding RNAs have been previously studied in certain protists (44), the full diversity of structured RNAs across protists remains largely unexplored.

To address this gap, we systematically surveyed conserved structured RNA motifs in protists using CMfinder (42) and R-scape (43). CMfinder detects nucleotide covariation patterns and evaluates the thermodynamic stability of hairpin structures, enabling the prediction of conserved RNA motifs from sequence alignments even under conditions of low primary sequence similarity (42). In a complementary approach, R-scape analyzes multiple RNA sequence alignments and computes a pairwise covariation statistic for each pair of aligned columns, thereby providing rigorous statistical support for conserved base-pairing interactions (43). Only RNA motifs with at least one statistically significant covariation signal according to R-scape were retained for downstream functional annotation.

Furthermore, we investigated the genomic contexts of these predicted motifs—such as their positional relationships to adjacent genes—to infer possible regulatory functions. These may include roles in transcription, translation, or alternative splicing. Collectively, this work aims to identify structured noncoding RNA motifs that could act as functional regulatory elements in protists.

### Analysis of the functions of these structured RNA candidates

Among the identified structured RNA motifs, RbcL_8943 and SRP_like_8976 were found to originate from the edible brown alga Porphyra haitanensis. Gene expression analysis revealed that the expression level of RbcL_8943 was lower than that of its adjacent coding sequences (CDS). This finding is consistent with our previous observations that many structured non-coding RNA motifs exhibit lower expression levels than neighboring coding regions (38). In contrast, SRP_like_8976 showed higher expression relative to its flanking CDS. As a putative signal recognition particle RNA (71), SRP_like_8976 is likely to operate independently of the adjacent coding sequence and may not co-regulate with it.

The PsaI_8813 RNA motif is at the end of the stop codon of both *psaI* and *psbJ* genes (Fig. 4C). Hairpin structures at mRNA 3′ termini sometimes protect transcripts from RNase degradation and enhance mRNA stability (87), while sometimes enhance RNA decay (For example, structure-mediated RNA decay by UPF1 and G3BP1(88)). In contrast, 5′ terminal hairpins may possess regulatory functions such as structured RNA elements called riboswitches (35,58,89,90) or ribosomal proteins binding sequences (91). Thus, the PsaI_8813 RNA motif might sense temperature variations through conformational changes, enabling psaI to mediate temperature-responsive functionality in addition to the stabilization of the mRNA (Fig. 4C). The most notable feature of this RNA motif is its capacity to form over 20 consecutive canonical Watson–Crick base pairs, which may enable it to function in the regulation of both the upstream genes, despite their opposing transcriptional orientations.

Certain RNA motifs, including Motif_11540, reside within gene introns and are typically positioned far (>100 bp) from splice sites. Despite this distance, long-range RNA–RNA interactions can modulate splicing outcomes, as evidenced by studies on fungal TPP riboswitches (56). For instance, in *Neurospora*, an intronic TPP riboswitch located approximately 500 nt from the 5’ splice site forms base pairs with sequences near the splice junction, thereby promoting splicing and facilitating translation under low thiamine pyrophosphate conditions (56). Thus, analogous mechanisms may be employed by the RNA motifs described here to regulate alternative splicing and, consequently, gene expression.

## Conclusions

By employing an integrated computational pipeline combining CMfinder and R-scape, we identified and assessed structured RNA motifs from a protist genomic database that includes numerous single-celled eukaryotic species. This pipeline has revealed the existence of a variety of novel structured ncRNA candidates. Genome contexts and similarities to known ncRNA motifs provide strong evidence for the biological functions of some newly found motifs. These motifs are likely involved in diverse regulatory processes, including the modulation of RuBisCO, phycocyanin synthesis, and photosynthetic regulation. Notably, we also identify a structured RNA candidate that may be capable of bidirectionally regulating two genes simultaneously. Our RT-PCR results show the expression of some of the motifs under different temperatures which may also be related to their regulatory functions for *Pyropia haitanensis* in responding to high temperature stress. However, more in-depth analyses are required to reveal their functions.

## Availability of data and materials

The datasets (such as genomes) were downloaded from NCBI (https://ftp.ncbi.nlm.nih.gov/refseq/release/protozoa). Most data generated or analyzed during this study are included in this published article. Stockholm files for the reported seven motifs are in the Supplementary File 2 (Part II). Custom Perl scripts are in the Supplementary Files 3 and 4. The remaining Stockholm files are available from the corresponding author on request.

## CRediT author statement

Yipeng Zhang: Writing–review & editing, Writing–original draft, Investigation. Qunying Liang: Investigation, Validation. Tianshu Wang: Investigation, Methodology. Yixin Huang: Software, Validation. Anqi Duan: Software, Methodology, Investigation. Jiaojiao Wang: Software, Validation, Investigation. Wenlei Wang: Methodology. Huan Yu: Investigation. Binfen Chen: Software, Validation. Zhilan Ying: Software. Yin Jia: Validation. Chen Fu: Data curation. Sanshu Li: Writing - original draft, Conceptualization, Resources, Supervision.

