## Supplementary File1-4 for "Identification of seven candidate structured noncoding RNA motifs in protists by comparative genomics": Supplementary File 1 Figure S1 and Table S1.docx

^3^ Current address: Institutes of Biomedical Sciences, Fudan University, Shanghai 200032, China

^4^ Current address: Anyang No.1 Middle School, Anyang 456000, China

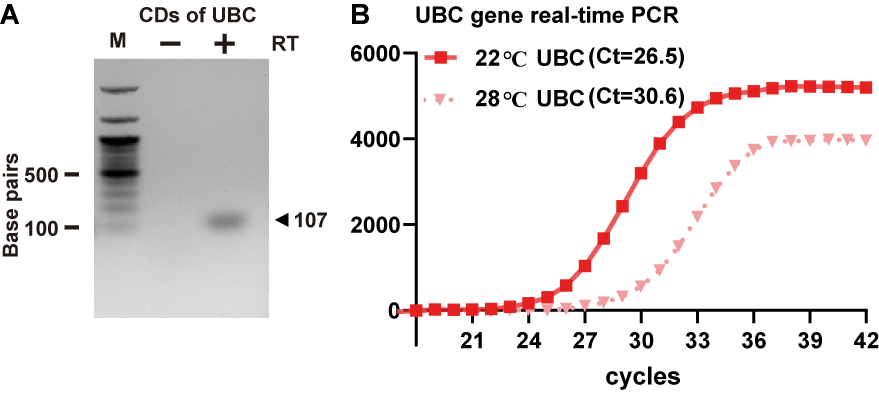

**Fig. S1** The ubiquitin-conjugating enzyme (UBC) encoding gene and its expression. A. the reverse-transcription PCR (RT-PCR) of the coding regions (CDs) of the UBC gene. B. Quantitative RT-PCR (RT-qPCR) for the UBC gene for algae cultured under temperature stress. Agarose gel shows the separation of RT-PCR products generated by using primers specific for the coding region as designated. M designates double-stranded DNA size markers, and lanes containing PCR products generated with (+) or without (–) the use of reverse transcriptase (RT) are identified. The experiment was repeated twice with similar results.

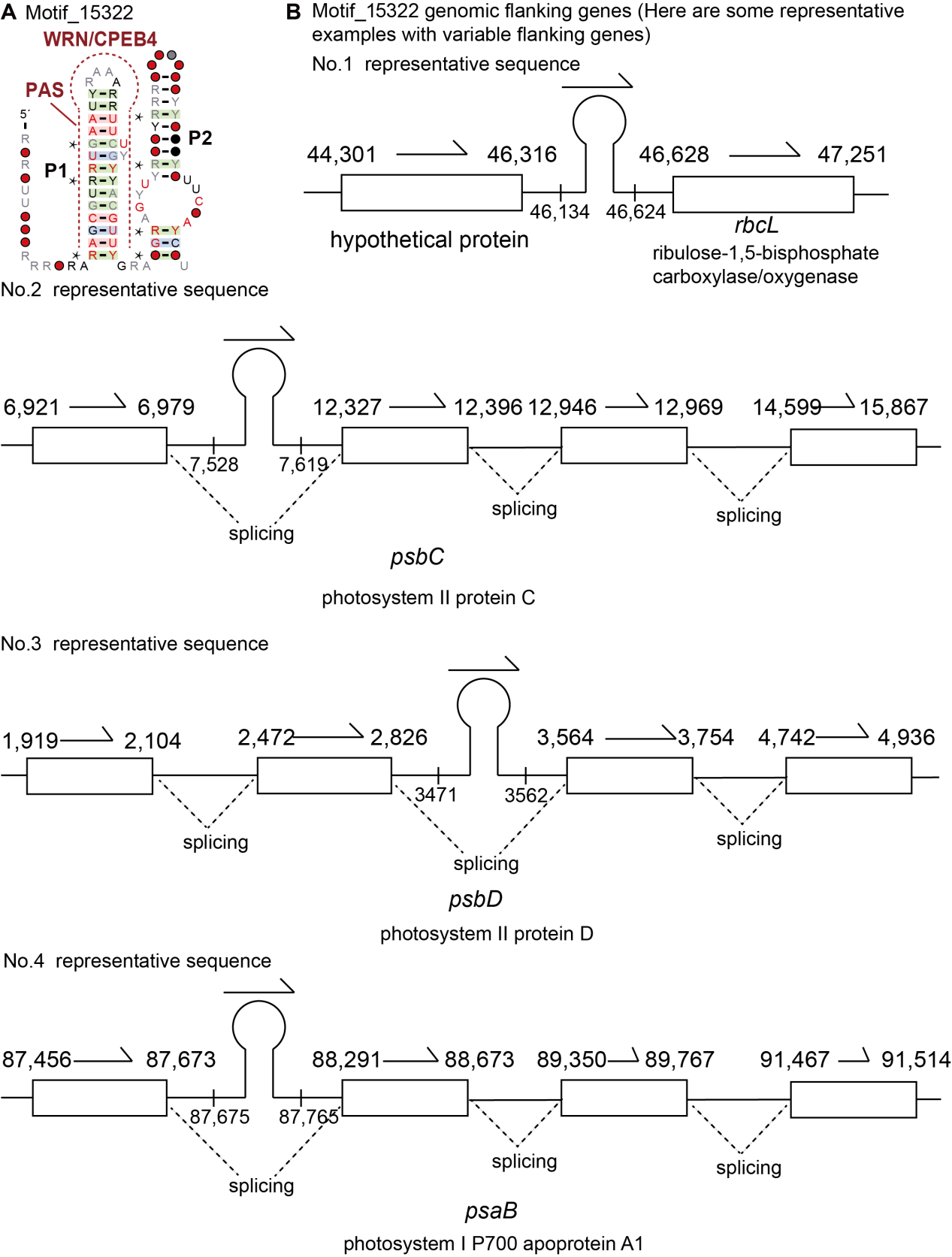

**Fig. S2.** **Structural and genomic features of t**he Motif_15322 RNA. (A) Consensus sequence and predicted secondary structure of the Motif_15322 RNA. The RBP (WRN and CPEB4) binding sites and the PAS sequence are annotated. (B) Genomic organization of the conserved RNA motif. Other annotations are as described for Figure 2.

**Table S1** Primers and RT-PCR product sizes

| Motif name | primer-F | primer-R | Product length (bp) |
| --- | --- | --- | --- |
| RbcL_8943 motif | GTGATAATTTTTCTATTAAGGTTGAACAC | GTTACAAATAATAAAATATATTGTAATATACTTAGCTTTC | 121 |
| RbcL_8943 cds | CTCAATCCGTAGAATCACGGAC | GCTAAAACATCTGTTTCTTTAATTACATAGTC | 121 |
| SRP_8976 motif | GCCCGAAAGCATATAATTGTAAAG | ACCCGAAAATCATAACTGTTG | 92 |
| SRP_8976 cds | CAATGTCTAAAAGAAGCTTCTCTAG | GACATGCCTCGAATTATGTAGTC | 92 |
| UBC gene | TCACAACGAGGATTTACCACC | GAGGAGCACCTTGGAAACG | 107 |

**Additional information: primers design and their positions on the genome**

I. RbcL_8943 RNA motif and its adjacent coding regions (Red uppercase letters indicate RNA motifs; lowercase letters denote coding sequences. ATG in red is the start codon. Underlined letters mark the primers used for PCR amplification of both the RNA motif and the coding region.)

ATCTTGATAGTGATAATTTTTCTATTAAGGTTGAACACTAGAAATCATTTTTCAATCATATATTTTTTTGAAAAACTTCTTTGTGACCATGAAAGCTAAGTATATTACAATATATTTTATTATTTGTAACTTATTTCTTATAATGGTATAAACAACGCAAAAGATACTGCCTTATAATCAAGGAGGAATACatgtctcaatccgtagaatcacggactaggattaaaagcgaacgttacgaatctggagtaatcccttacgctaaaatgggttactgggatgctgactatgtaattaaagaaacagatgttttagctctgttcagaatcactcctcagccgggtgttgaccctattgaagcatctgctgca

8943_motif_F (Tm:53℃)

5' GTGATAATTTTTCTATTAAGGTTGAACAC

8943_motif_R (53℃)

5' GTTACAAATAATAAAATATATTGTAATATACTTAGCTTTC

8943_cds_F (56℃)

5' ctcaatccgtagaatcacggac

8943_cds_R (55℃)

5' gctaaaacatctgtttctttaattacatagtc

II. SRP_8976 RNA motif (annotations are the same as the one above. Three letters “taa” in red is the stop codon)

tcaatgtctaaaagaagcttctctagaaatgcttggtcaagatgatatcaaaattattgcgccttattttgactacataattcgaggcatgtcataaATAGTAAATTCAATTAATTAGTTGAATAAAAGCTTGTATAATGTTATGATTAGTTCAAGCCCGAAAGCATATAATTGTAAAGACTAAATCTTGTCAGAACCGGAAGGTAGCAGCAATAATGTTTAATTACAACAGTTATGATTTTCGGGTGTTTACTATTTGTA

8976_motif_F (Tm: 54℃)

5' GCCCGAAAGCATATAATTGTAAAG

8976_motif_R (Tm: 52℃)

5' ACCCGAAAATCATAACTGTTG

8976_cd_F (53℃)

5' CAATGTCTAAAAGAAGCTTCTCTAG

8976_cd_R (54℃)

5' GACATGCCTCGAATTATGTAGTC
