## Supplementary File1-4 for "Identification of seven candidate structured noncoding RNA motifs in protists by comparative genomics": Supplementary File 2 species, sto-files, and original gels.docx

^3^ Current address: Institutes of Biomedical Sciences, Fudan University, Shanghai 200032, China

^4^ Current address: Anyang No.1 Middle School, Anyang 456000, China

Part I Species in the database----------------------Page 2-8

Part II Stockholm files for nine RNA motifs---------Page 8-59

Stockholm files for motif­_8943----------------------page 9

Stockholm files for motif­_8976----------------------page 10

Stockholm files for motif­_8813----------------------page 29

Stockholm files for motif­_4508----------------------page 37

Stockholm files for motif­_5095----------------------page 38

Stockholm files for motif­_11540---------------------page 40

Stockholm files for motif­_15322---------------------page 50

Part III Original gel pictures-----------------------Page 54-58

Part I Species in the database

```
Totally, 894 species.
Acanthamoeba castellanii; Acanthoceras zachariasii; Achlya hypogyna; Acrasis kona; Acrochaetium secundatum; Acrosorium ciliolatum; Actinocyclus subtilis; Acytostelium subglobosum; Agarum clathratum; Ahnfeltia fastigiata; Ahnfeltia plicata; Alaria crassifolia; Alaria crispa; Alaria esculenta; Alaria marginata; Alaria praelonga; Amoeboaphelidium protococcarum; Amplisiphonia pacifica; Analipus japonicus; Ancoracysta twista; Ancyromonas sigmoides; Andalucia godoyi; Antithamnionella miharai; Anunuuluaehu liula; Aphanocladia delicatula; Aphanocladia stichidiosa; Aphanomyces astaci; Aphanomyces invadans; Apophlaea sinclairii; Ascoseira mirabilis; Asparagopsis taxiformis; Asterionella formosa; Asterionellopsis glacialis; Astrosyne radiata; Attheya longicornis; Aureococcus anophagefferens; Aureophycus aleuticus; Aureoumbra lagunensis; Babesia bigemina; Babesia bovis; Babesia caballi; Babesia duncani; Babesia microti; Babesia orientalis; Babesia ovata; Bacillaria paxillifer; Bakuella subtropica; Balamuthia mandrillaris; Balbiania investiens; Bangia atropurpurea; Bangia fuscopurpurea; Bangiopsis subsimplex; Batrachospermum macrosporum; Berkeleya fennica; Besnoitia besnoiti; Betaphycus gelatinus; Biddulphia biddulphiana; Biddulphia tridens; Bigelowiella natans; Blastocystis hominis; Boldia erythrosiphon; Bostrychia moritziana; Bostrychia simpliciuscula; Bostrychia tenella; Bostrychia tenuissima; Bremia lactucae; Bryothamnion seaforthii; Bulboplastis apyrenoidosa; Cafeteria roenbergensis; Cafileria marina; Calcarina hispida; Calliarthron tuberculosum; Caloglossa beccarii; Caloglossa intermedia; Caloglossa monosticha; Campylaephora boydenii; Campylaephora kondoi; Campylaephora sungminbooi; Capsaspora owczarzaki; Catenella fusiformis; Caulacanthus okamurae; Caulacanthus ustulatus; Cavenderia fasciculata; Centroceras gasparrinii; Ceramium japonicum; Ceramium sungminbooi; Cerataulina daemon; Chaetoceros gelidus; Chaetoceros muellerii; Chaetoceros simplex; Characiopsis acuta; Chattonella marina; Chondria tumulosa; Chondrus crispus; Chorda asiatica; Chorda filum; Choreocolax polysiphoniae; Choristocarpus tenellus; Chromera velia; Chroodactylon ornatum; Chroomonas placoidea; Chroothece richteriana; Chrysochromulina parva; Chrysodidymus synuroideus; Chrysotila carterae; Cladosiphon okamuranus; Cliftonaea pectinata; Cocconeiopsis kantsiensis; Coccophora langsdorfii; Coeloseira compressa; Colacium mucronatum; Colpomenia peregrina; Colpomenia sinuosa; Compsopogon caeruleus; Corallina chilensis; Corallina ferreyrae; Corallina officinalis; Coscinodiscus granii; Coscinodiscus radiatus; Coscinodiscus wailesii; Costaria costata; Crassa caudata; Crassiphycus birdiae; Crassiphycus corneus; Crassiphycus crassissimus; Cryptocaryon irritans; Cryptoglena skujai; Cryptomonas curvata; Cryptomonas gyropyrenoidosa; Cryptomonas paramecium; Cryptomonas pyrenoidifera; Cryptosporidium andersoni; Cryptosporidium bovis; Cryptosporidium hominis; Cryptosporidium muris; Cryptosporidium parvum; Cryptosporidium ryanae; Cryptosporidium ubiquitum; Cumathamnion serrulatum; Cutleria multifida; Cyanidiococcus yangmingshanensis; Cyanidioschyzon merolae; Cyanidium caldarium; Cyanidium chilense; Cyanophora biloba; Cyanophora paradoxa; Cyanophora sudae; Cyanoptyche gloeocystis; Cyclospora cayetanensis; Cyclotella cryptica; Cyclotella pseudostelligera; Cylindrotheca closterium; Cymathaere triplicata; Cystoclonium purpureum; Cystoisospora suis; Dasya binghamiae; Dasya naccarioides; Dasyclonium flaccidum; Dasysiphonia japonica; Deltalsia parasitica; Dermonema virens; Desmarestia aculeata; Desmarestia viridis; Diacronema viridis; Dichotomaria marginata; Dictyocha speculum; Dictyomenia sonderi; Dictyoneurum californicum; Dictyopteris divaricata; Dictyostelium citrinum; Dictyostelium discoideum; Dictyostelium fasciculatum; Dictyostelium intermedium; Dictyostelium purpureum; Dictyota dichotoma; Dictyotopsis propagulifera; Didymosphenia geminata; Digenea simplex; Diphylleia rotans; Dipterocladia arabiensis; Dipterosiphonia australica; Discoplastis spathirhyncha; Ditylum brightwellii; Dixoniella grisea; Durinskia baltica; Echinothamnion hookeri; Echinothamnion hystrix; Ecklonia cava; Ecklonia maxima; Ecklonia radiata; Ectocarpus siliculosus; Egregia menziesii; Eimeria acervulina; Eimeria chapmani; Eimeria furonis; Eimeria kofoidi; Eimeria kongi; Eimeria legionensis; Eimeria maxima; Eimeria mitis; Eimeria necatrix; Eimeria stiedai; Eimeria tenella; Eisenia bicyclis; Emiliania huxleyi; Endarachne binghamiae; Entamoeba dispar; Entamoeba histolytica; Entamoeba invadens; Entamoeba nuttalli; Erythrolobus coxiae; Erythrotrichia carnea; Eucampia zodiacus; Eucheuma denticulatum; Euglena agilis; Euglena archaeoplastidiata; Euglena clara; Euglena deses; Euglena gracilis; Euglena hiemalis; Euglena longa; Euglena undulata; Euglena viridis; Euglenaformis proxima; Euglenaria anabaena; Eunotia naegelii; Euplotes cristatus; Euplotes vanleeuwenhoeki; Euplotes vannus; Eutreptiella eupharyngea; Eutreptiella gymnastica; Fibrocapsa japonica; Fistulifera saprophila; Fistulifera solaris; Flexiglena variabilis; Florenciella parvula; Fonticula alba; Fragilariopsis cylindrus; Fucus distichus; Fucus vesiculosus; Fushitsunagia catenata; Galaxaura rugosa; Galdieria phlegrea; Galdieria sulphuraria; Gelidiella acerosa; Gelidiella fanii; Gelidiella flabella; Gelidium arborescens; Gelidium coulteri; Gelidium elegans; Gelidium gabrielsonii; Gelidium galapagense; Gelidium isabelae; Gelidium kathyanniae; Gelidium sclerophyllum; Gelidium sinicola; Gelidium vagum; Gephyrocapsa ericsonii; Gephyrocapsa muellerae; Gephyrocapsa oceanica; Gephyrocapsa parvula; Giardia duodenalis; Giardia intestinalis; Glaucocystis incrassata; Glaucocystis nostochinearum; Glaucosphaera vacuolata; Gloeochaete wittrockiana; Gloiopeltis furcata; Gomphonema parvulum; Goniotrichopsis reniformis; Gonyostomum semen; Gracilaria baiana; Gracilaria bursa-pastoris; Gracilaria caudata; Gracilaria cearensis; Gracilaria cervicornis; Gracilaria changii; Gracilaria chilensis; Gracilaria chorda; Gracilaria chouae; Gracilaria cliftonii; Gracilaria domingensis; Gracilaria edulis; Gracilaria eucheumatoides; Gracilaria ferox; Gracilaria firma; Gracilaria flabelliformis; Gracilaria gracilis; Gracilaria hainanensis; Gracilaria isabellana; Gracilaria multipartita; Gracilaria ornata; Gracilaria pachydermatica; Gracilaria pacifica; Gracilaria salicornia; Gracilaria spinulosa; Gracilaria tenuistipitata; Gracilaria textorii; Gracilaria tikvahiae; Gracilaria usneoides; Gracilaria vermiculophylla; Gracilariophila oryzoides; Gracilariopsis andersonii; Gracilariopsis chorda; Gracilariopsis heteroclada; Gracilariopsis lemaneiformis; Gracilariopsis longissima; Gracilariopsis mclachlanii; Gracilariopsis tenuifrons; Grateloupia angusta; Grateloupia asiatica; Grateloupia cornea; Grateloupia elliptica; Grateloupia filicina; Grateloupia livida; Grateloupia taiwanensis; Grateloupia turuturu; Gredgaria maugeana; Gregarina niphandrodes; Griffithsia okiensis; Guillardia theta; Guinardia delicatula; Guinardia striata; Gymnochlora stellata; Haemoproteus belopolskyi; Haemoproteus columbae; Haemoproteus micronuclearis; Haemoproteus nisi; Haemoproteus payevskyi; Halamphora americana; Halamphora calidilacuna; Halamphora coffeaeformis; Halosiphon tomentosus; Halteria grandinella; Halymenia maculata; Hapterophycus canaliculatus; Haramonas pauciplastida; Hartmannella vermiformis; Haslea avium; Haslea karadagensis; Haslea nusantara; Haslea ostrearia; Haslea provincialis; Haslea pseudostrearia; Haslea tsukamotoi; Hedophyllum nigripes; Helicotheca tamesis; Helminthocladia australis; Helminthora furcellata; Hemiaulus sinensis; Hemiselmis andersenii; Hepatozoon catesbianae; Herposiphonia versicolor; Heterosigma akashiwo; Heterosiphonia pulchra; Hildenbrandia rivularis; Hildenbrandia rubra; Histiona aroides; Histomonas meleagridis; Hommersandiophycus borowitzkae; Hyaloperonospora arabidopsidis; Hydropuntia rangiferina; Hydropuntia urvillei; Hypnea brasiliensis; Hypnea cervicornis; Hypnea cornuta; Hypnea cryptica; Hypnea edeniana; Hypnea flava; Hypnea marchantiae; Hypnea musciformis; Hypnea nidifica; Hypnea nidulans; Hypnea pseudomusciformis; Hypnea spinella; Hypnea wynnei; Ichthyophthirius multifiliis; Imasa heleensis; Ishige okamurae; Isochrysis galbana; Isospora picoflavae; Isospora serinuse; Izziella formosana; Jakoba bahamiensis; Jakoba libera; Kappaphycus alvarezii; Kappaphycus malesianus; Kappaphycus striatus; Klossia helicina; Klossia razorbacki; Kryptoperidinium foliaceum; Kuetzingia canaliculata; Kumanoa ambigua; Kumanoa americana; Kumanoa mahlacensis; Laminaria digitata; Laminaria hyperborea; Laminaria rodriguezii; Laminaria solidungula; Laurencia catarinensis; Laurencia elata; Laurencia obtusa; Laurenciella marilzae; Legerella nova; Leishmania braziliensis; Leishmania donovani; Leishmania enriettii; Leishmania infantum; Leishmania major; Leishmania martiniquensis; Leishmania mexicana; Leishmania orientalis; Leishmania panamensis; Leishmania tarentolae; Lepidodinium chlorophorum; Lepocinclis ovum; Lepocinclis playfairiana; Lepocinclis steinii; Lepocinclis tripteris; Leptocylindrus danicus; Leptomonas pyrrhocoris; Lessonia flavicans; Lessonia nigrescens; Lessonia spicata; Lessonia variegata; Leucocryptos marina; Leucocytozoon caulleryi; Leucocytozoon fringillinarum; Leucocytozoon majoris; Leucocytozoon sabrazesi; Liagora brachyclada; Liagora harveyana; Liagoropsis maxima; Lietzensia polymorpha; Lithodesmioides polymorpha; Lithodesmium undulatum; Lithothamnion corallioides; Lophocladia kuetzingii; Lophurella caespitosa; Lophurella hookeriana; Lophurella mutabilis; Lotharella oceanica; Lotharella vacuolata; MAG Porospora; Macrocystis integrifolia; Macrocystis pyrifera; Madagascaria erythrocladioides; Malawimonas californiana; Malawimonas jakobiformis; Mallomonas splendens; Mastocarpus papillatus; Melanothamnus gigas; Melanothamnus harveyi; Melanthalia intermedia; Melosira undulata; Membranoptera platyphylla; Membranoptera tenuis; Membranoptera weeksiae; Meringosphaera mediterranea; Merotricha bacillata; Meteora sporadica; Mimica arnoldii; Minidiscus spinulatus; Ministeria vibrans; Minutocellus polymorphus; Monocercomonoides exilis; Monomorphina aenigmatica; Monomorphina parapyrum; Monosiga brevicollis; Myagropsis myagroides; Naegleria fowleri; Naegleria gruberi; Naegleria lovaniensis; Nannochloropsis gaditana; Nannochloropsis granulata; Nannochloropsis limnetica; Nannochloropsis oceanica; Nannochloropsis oculata; Nannochloropsis salina; Navicula arenaria; Navicula ramosissima; Navicula veneta; Nemalion vermiculare; Neogoniolithon spectabile; Neoizziella asiatica; Neoporphyra dentata; Neoporphyra seriata; Neorhodella cyanea; Neorotalia gaimardi; Neospora caninum; Nereocystis luetkeana; Neustupella aerophytica; Nitzschia alba; Nitzschia dissipata; Nitzschia dubiiformis; Nitzschia ovalis; Nitzschia supralitorea; Nitzschia traheaformis; Nuclearia simplex; Nycteria heischi; Ochromonas danica; Ochrosphaera neapolitana; Odontella aurita; Odontella regia; Odontella sinensis; Olisthodiscus luteus; Ophidocladus simpliciusculus; Osmundaria fimbriata; Osmundea sinicola; Pachymeniopsis lanceolata; Palisada intermedia; Palmaria decipiens; Palmaria palmata; Palpitomonas bilix; Papenfussiella kuromo; Paracercomonas marina; Parahaemoproteus vireonis; Paralagenidium karlingii; Paralia sulcata; Paramecium aurelia; Paramecium caudatum; Paramecium gigas; Paramecium tetraurelia; Paramoeba pemaquidensis; Paravannella minima; Partenskyella glossopodia; Paulinella chromatophora; Paulinella micropora; Pavlova lutheri; Pelagomonas calceolata; Periphykon beckeri; Perkinsus marinus; Perkinsus olseni; Peronosclerospora sorghi; Peronospora belbahrii; Peronospora tabacina; Petalonia fascia; Phacus arnoldii; Phacus inflexus; Phacus pleuronectes; Phaeocystis antarctica; Phaeocystis globosa; Phaeocystis rex; Phaeodactylum tricornutum; Phaeostrophion irregulare; Pharyngomonas kirbyi; Phymatolithon calcareum; Physarum polycephalum; Phytophthora agathidicida; Phytophthora aleatoria; Phytophthora amnicola; Phytophthora andina; Phytophthora aquimorbida; Phytophthora asparagi; Phytophthora boehmeriae; Phytophthora botryosa; Phytophthora brassicae; Phytophthora cactorum; Phytophthora cambivora; Phytophthora capensis; Phytophthora capsici; Phytophthora captiosa; Phytophthora castaneae; Phytophthora cinnamomi; Phytophthora citricola; Phytophthora citrophthora; Phytophthora clandestina; Phytophthora cocois; Phytophthora colocasiae; Phytophthora crassamura; Phytophthora cryptogea; Phytophthora elongata; Phytophthora europaea; Phytophthora fallax; Phytophthora fluvialis; Phytophthora fragariae; Phytophthora gemini; Phytophthora gibbosa; Phytophthora gonapodyides; Phytophthora gregata; Phytophthora hedraiandra; Phytophthora heveae; Phytophthora humicola; Phytophthora ilicis; Phytophthora infestans; Phytophthora insolita; Phytophthora ipomoeae; Phytophthora kernoviae; Phytophthora lactucae; Phytophthora lateralis; Phytophthora litchii; Phytophthora litoralis; Phytophthora meadii; Phytophthora megakarya; Phytophthora mirabilis; Phytophthora multivesiculata; Phytophthora multivora; Phytophthora nemorosa; Phytophthora nicotianae; Phytophthora obscura; Phytophthora ornamentata; Phytophthora palmivora; Phytophthora parasitica; Phytophthora phaseoli; Phytophthora pinifolia; Phytophthora plurivora; Phytophthora pluvialis; Phytophthora polonica; Phytophthora pseudotsugae; Phytophthora quercetorum; Phytophthora quercina; Phytophthora ramorum; Phytophthora rubi; Phytophthora sansomeana; Phytophthora sojae; Phytophthora taxon; Phytophthora thermophila; Phytophthora tropicalis; Phytophthora tubulina; Phytophthora versiformis; Phytophthora x; Phytopythium vexans; Plagiogramma staurophorum; Plagiogrammopsis vanheurckii; Plasmodium berghei; Plasmodium brasilianum; Plasmodium chabaudi; Plasmodium coatneyi; Plasmodium cynomolgi; Plasmodium falciparum; Plasmodium floridense; Plasmodium fragile; Plasmodium gaboni; Plasmodium gallinaceum; Plasmodium gonderi; Plasmodium homocircumflexum; Plasmodium inui; Plasmodium juxtanucleare; Plasmodium knowlesi; Plasmodium malariae; Plasmodium mexicanum; Plasmodium reichenowi; Plasmodium relictum; Plasmodium simium; Plasmodium vinckei; Plasmodium vivax; Plasmodium yoelii; Plasmopara halstedii; Plasmopara viticola; Platysiphonia delicata; Pleurocladia lacustris; Pleurophycus gardneri; Pleurosigma inscriptura; Pleurosigma intermedium; Pleurostichidium falkenbergii; Pleurostomum flabellatum; Plocamiocolax pulvinata; Plocamium cartilagineum; Polychromophilus murinus; Polymyxa betae; Polyopes affinis; Polyopes lancifolius; Polysiphonia brodiei; Polysiphonia elongata; Polysiphonia infestans; Polysiphonia morrowii; Polysiphonia schneideri; Polysiphonia scopulorum; Polysiphonia sertularioides; Polysiphonia stricta; Polysphondylium pallidum; Porcisia hertigi; Porolithon onkodes; Porphyra pulchra ( red alga; Porphyra purpurea; Porphyra umbilicalis; Porphyra yezoensis; Porphyridium aerugineum; Porphyridium purpureum; Porphyridium sordidum; Poterioochromonas malhamensis; Proteomonas sulcata; Proteromonas lacertae; Psammodictyon constrictum; Psammoneis japonica; Psammoneis obaidii; Pseudellipsoidion edaphicum; Pseudo-nitzschia americana; Pseudo-nitzschia cuspidata; Pseudo-nitzschia delicatissima; Pseudo-nitzschia micropora; Pseudo-nitzschia multiseries; Pseudo-nitzschia multistriata; Pseudo-nitzschia pungens; Pseudo-nitzschia simulans; Pseudoceramium inkyui; Pseudoceramium tenerrimum; Pseudoerythrocladia kornmannii; Pseudopedinella elastica; Pseudoperonospora cubensis; Pseudoperonospora humuli; Pteridomonas danica; Pterocladia media; Pterocladia mexicana; Pterocladia musciformis; Pterocladia robusta; Pterocladiella capillacea; Pterosiphonia complanata; Pterothamnion plumula; Pulvinaster venetus; Pylaiella littoralis; Pyropia fucicola; Pyropia haitanensis; Pyropia kanakaensis; Pyropia nitida; Pyropia perforata; Pyropia tenera; Pyropia yezoensis; Pythium insidiosum; Pythium ultimum; Reclinomonas americana; Rhizochromulina marina; Rhizosolenia fallax; Rhizosolenia imbricata; Rhizosolenia setigera; Rhodaphanes brevistipitata; Rhodella violacea; Rhodochorton tenue; Rhodomela confervoides; Rhodomelopsis africana; Rhodomonas salina; Rhodospora sordida; Rhodymenia pseudopalmata; Roundia cardiophora; Saccharina angustata; Saccharina coriacea; Saccharina diabolica; Saccharina japonica; Saccharina latissima; Saccharina longipedalis; Saccharina longissima; Saccharina ochotensis; Saccharina religiosa; Saccharina sculpera; Sahlingia subintegra; Salpingoeca rosetta; Saprolegnia diclina; Saprolegnia ferax; Saprolegnia parasitica; Sarcopeltis skottsbergii; Sargassum aquifolium; Sargassum confusum; Sargassum feldmannii; Sargassum fluitans; Sargassum fulvellum; Sargassum fusiforme; Sargassum graminifolium; Sargassum hemiphyllum; Sargassum henslowianum; Sargassum horneri; Sargassum ilicifolium; Sargassum kjellmanianum; Sargassum macrocarpum; Sargassum mcclurei; Sargassum muticum; Sargassum natans; Sargassum nigrifolium; Sargassum patens; Sargassum phyllocystum; Sargassum plagiophyllum; Sargassum polycystum; Sargassum serratifolium; Sargassum siliquastrum; Sargassum spinuligerum; Sargassum thunbergii; Sargassum vachellianum; Sargassum yezoense; Schimmelmannia schousboei; Schizocladia ischiensis; Schizymenia dubyi; Scinaia undulata; Scytosiphon canaliculatus; Scytosiphon lomentaria; Scytosiphon promiscuus; Scytothamnus australis; Sebdenia flabellata; Seculamonas ecuadoriensis; Sheathia arcuata; Silvetia siliquosa; Sirodotia delicatula; Skeletonema costatum; Skeletonema grevillei; Skeletonema marinoi; Skeletonema potamos; Skeletonema pseudocostatum; Skeletonema subsalsum; Skeletonema tropicum; Sonderella linearis; Sphaeroforma arctica; Sphaerothecum destruens; Spironucleus salmonicida; Spirostomum yagiui; Spongospora subterranea; Sporochnus bolleanus; Sporolithon durum; Spyridia filamentosa; Stachyamoeba lipophora; Stephanodiscus niagarae; Stephanopyxis turris; Strombomonas costata; Stylonema alsidii; Symphyocladia dendroidea; Symphyocladia marchantioides; Symphyocladiella dendroidea; Synarthrophyton chejuense; Synarthrophyton patena; Synedra acus; Synura uvella; Syringoderma abyssicola; Taenioma perpusillum; Tayloriella tenebrosa; Teleaulax amphioxeia; Tenuicylindrus belgicus; Tetrahymena malaccensis; Tetrahymena paravorax; Tetrahymena pigmentosa; Tetrahymena pyriformis; Tetrahymena rostrata; Tetrahymena thermophila; Thalassionema bacillare; Thalassionema frauenfeldii; Thalassionema nitzschioides; Thalassiosira nordenskioeldii; Thalassiosira oceanica; Thalassiosira profunda; Thalassiosira pseudonana; Thalassiosira rotula; Thalassiosira tenera; Thalassiosira weissflogii; Thaumatella adunca; Thecamoeba quadrilineata; Thecamonas trahens; Theileria annulata; Theileria equi; Theileria lestoquardi; Theileria luwenshuni; Theileria orientalis; Theileria parva; Theileria taurotragi; Theileria velifera; Thorea hispida; Thraustotheca clavata; Thuretia quercifolia; Thuricola similis; Timspurckia oligopyrenoides; Tisochrysis lutea; Titanophycus setchellii; Tolypiocladia glomerulata; Toxarium undulatum; Toxoplasma gondii; Trachelomonas volvocina; Trachydiscus minutus; Triceratium dubium; Trichogloeopsis pedicelleta; Trichomonas vaginalis; Triparma laevis; Tritrichomonas foetus; Tryblionella apiculata; Trypanosoma brucei; Trypanosoma conorhini; Trypanosoma cruzi; Trypanosoma equiperdum; Trypanosoma grayi; Trypanosoma melophagium; Trypanosoma rangeli; Trypanosoma theileri; Tsukubamonas globosa; Tsunamia transpacifica; Turbinaria ornata; Ulnaria ulna; Undaria peterseniana; Undaria pinnatifida; Uronema marinum; Vacuolaria virescens; Vaucheria litorea; Vertebrata australis; Vertebrata isogona; Vertebrata lanosa; Vertebrata thuyoides; Vischeria punctata; Vischeria stellata; Wildemania schizophylla; Xiphosiphonia pinnulata; Yamadaella caenomyce.


Part II Stockholm files for nine RNA motifs

#for motif_8943
# STOCKHOLM 1.0

NC_029861.1/116744-116880         AUCUUGAUAGUGAUAAUUUUUCUAUUAAGGUUGAACACUAGAAGUAA.UUCU.CuAAG..CAAAUAUUUA..........CUU.G.AGAA...ACUUCUUuGUGACCAUAAAAGCUUAAUUAUAUUACAAUAUAUUUUGUUAUUUGUAACUUAGUU
#=GR NC_029861.1/116744-116880 PP ***********************************************.****.99***..**********..........***.*.****...******99*******************************************************
NC_035573.1/29201-29339           AUCUUAAUAGUGAUAAUUUCUAUAUUAAGGUUGAACACAAGCAAGAA.CUUUaC.GAA..UAGUUUUUUA..........UUC.G.AAAAauaCUUGCUU.GUGACCAUAAAAGCUUAGUUAUAUUACAAUAUAUUUUAUUGUUUGUAAAUCAGUU
#=GR NC_035573.1/29201-29339   PP ***********************************************.99999*.***..**********..........***.*.88888*********.*******************************************************
NC_007932.1/29375-29511           AUCUUGAUAGUGAUAAUUUUUAUAUUAAGGUUGAACACUAGAAAUAAcUUUU.C.AAG..CAAACUCUUA..........UUU.G.AAAA...AUUUCUCuGUGACCAUAAAAACUUAAUUAUAUUACAAUAUAUUUUAUUAUUUGUAACUUAGUU
#=GR NC_007932.1/29375-29511   PP ***********************************************99999.*.***..**********..........***.*.****...******99*******************************************************
NC_000925.1/29257-29395           AUCUUAAUAGUGAUAGUUUCUAUAUUAAGGUUGAACACAAAGAAGAG.UUUU.CuAAA..UAGUUCUUUA..........UUUgG.AAAAu.aCUUCUUU.GUGACCAUAGAAGCUUAGUUAUAUUACAAUAUAUUUUAUUAUUUGUAACUUCGUU
#=GR NC_000925.1/29257-29395   PP ***********************************************.9999.97888..**********..........88868.8888*.********.*******************************************************
NC_024050.1/29346-29482           AUCUUGAUAGUGGUAAUUUCUUUAUUAAGGUUGAACACUAGAAAUAA.CUUU.U.CAAguAAAUAUUUUU..........UUG.A.AAAA...CUUAUUU.GUGACCAUAAAAACUUAACUAUAUUACAAUAUAUUUUAUUAUUUGUAACUUAGUU
#=GR NC_024050.1/29346-29482   PP ***********************************************.9999.8.888866667777777..........888.8.8888...*******.*******************************************************
NC_021189.1/29307-29442           AUCUUGAUAGUGAUAAUUUUUCUAUUAAGGUUGAACACUAGAAAUCA.UUUU.U.CAAucAUAUAUUUUU..........UUG.A.AAAA...CUUCUUU.GUGACCAUGAAAGCU-AAGUAUAUUACAAUAUAUUUUAUUAUUUGUAACUUAUUU
#=GR NC_021189.1/29307-29442   PP ***********************************************.****.*.***877788888888..........***.*.****...*******.**************9.999************************************
#=GC SS_cons                      <<<<<<<<<_____________>>>>>>>>>,,,,<<<-<<<<<<--.<<<<.<.<<<..__________~~~~~~~~~.>>>.>.>>>>...>>>>>>-.>>>--------------------<<<<<<_____________>>>>>>:::::::
#=GC RF                           auCuuaauaGUGAUAAUUUuUauauuaaGguUGAACaCuaggaagAA.uuuu.c.gaa..uAauuuUUUa~~~~~~~~~.uuc.g.aaaa...cuuccuU.GuGACCAUAAAAgCUUAauUAUAuuaCaaUAUAUUUUAUUAUuuGuaacUUAGUU
//

#for Motif_8976
# STOCKHOLM 1.0

NC_024050.1/7243-7334             GCCCGAAAG..CGUA....UA...AU.UGUAA......AAAC.UAAAUCUUGUCAGAACCGGAAGGUAGCAGCAAUAAU.GUUUAAU.....UACA.A.C.AGUUAUGA.UUUUCGGGU
#=GR NC_024050.1/7243-7334     PP *********..****....**...**.*****......****.************************************.*******.....****.*.*.********.*********
NC_021189.1/7222-7313             GCCCGAAAG..CAUA....UA...AU.UGUAA......AGAC.UAAAUCUUGUCAGAACCGGAAGGUAGCAGCAAUAAU.GUUUAAU.....UACA.A.C.AGUUAUGA.UUUUCGGGU
#=GR NC_021189.1/7222-7313     PP *********..****....**...**.*****......****.************************************.*******.....****.*.*.********.*********
NC_007932.1/7229-7320             GCCCGAGAG..CAUA....UA...AU.UGUAA......AAAC.UAAAUCUUGUCAGAACCGGAAGGUAGCAGCAAUAAU.GUUUCAA.....UACA.A.A.AGUUAUGA.UUUUCGGGU
#=GR NC_007932.1/7229-7320     PP *********..****....**...**.*****......****.************************************.*******.....****.*.*.********.*********
NC_029861.1/92666-92758           GCCCGAAAG..CAUA....UA...AU.UGUAA......AAAC.UAAAUCUUGUCAGAACCGGAAGGUAGCAGCAAUAAC.GUUUAAC.....UACA.A.A.AGUUAUGAcUUUUCGGGU
#=GR NC_029861.1/92666-92758   PP *********..****....**...**.*****......****.************************************.*******.....****.*.*.*******967899*****
NC_000925.1/7203-7294             GCCCGAAAG..UAUA....UA...AU.UGUAG......AAAC.UAAAUCUUGUCAGAACCGGAAGGUAGCAGCAAUAAU.GUUUACA.....UAUA.G.A.AGAUAUGA.UUUUCGGGU
#=GR NC_000925.1/7203-7294     PP *********..****....**...**.*****......****.************************************.*******.....****.*.*.********.*********
NC_029576.1/7217-7308             GCCCGAAAA..CAUA....UA...AU.UGUAG......AAGC.AAAAUCUUGUCAGAACCGGAAGGUAGCAGCAAUAAU.GUUUAUC.....UAUA.A.C.AGUUAUGA.CUUUCGGGU
#=GR NC_029576.1/7217-7308     PP *********..****....**...**.*****......****.************************************.*******.....****.*.*.********.*********
NC_035573.1/7200-7291             UCCCGAAAG..CGUA....UA...AU.UGUAU......AAAC.AUAAUCUUGUCAGAACCGGAAGGUAGCAGCAAUAAU.GUUUACC.....UACA.A.A.AGAUAUGA.UUUUCGGGU
#=GR NC_035573.1/7200-7291     PP *********..****....**...**.*****......****.************************************.*******.....****.*.*.********.*********
NC_035231.1/10119-10210           GCUCGGAAA..UUUA....UU...AA.UGUAA......UAGC.AAAAACUUGUCAGAACUAGACAGUAGCAGCAAUAAA.GUUUUAU.....UACA.G.A.UAGUAAAU.UUUCCGAGU
#=GR NC_035231.1/10119-10210   PP *********..****....**...**.*****......****.************************************.*******.....****.*.*.********.*********
NC_031178.1/108741-108833         GCUCGGGAA..UUUA....UA...AA.UGUAA......UAGC.AUAAACUUGUCAGGACUGGAAAGUAGCAGCAAUAAU.GCUUAAU.....UACA.A.AcAGAUAAAU.UUUCCGAGC
#=GR NC_031178.1/108741-108833 PP *********..****....**...**.*****......****.************************************.*******.....***9.6.53899*****.*********
NC_031171.1/115621-115529         ACUCGGAGA..UUUA....UA...AA.UGUAA......UAGCcAAAAACUUGUCAGAACUGGAAAGUAGCAGCAAUAUA.GCUUAAU.....UACA.A.U.AAGUAAAU.UCUUCGAGU
#=GR NC_031171.1/115621-115529 PP *********..****....**...**.*****......77751667899******************************.*******.....****.*.*.********.*********
NC_031147.1/127154-127063         GCUCGGAAG..UUUA....UA...GA.UAUAG......UAGC.UCAAACUUGUCAGAACUGGAAAGUAGCAGCAAAAAA.GCUAAAU.....UAUA.G.U.AAGUAAAU.CUUUCGAGU
#=GR NC_031147.1/127154-127063 PP *********..****....**...**.*****......****.************************************.*******.....****.*.*.********.*********
NC_031664.1/23490-23399           GCUCGGAGA..UUCA....CA...AA.UGUAA......UAGC.UCAAACUUGUCAGAACUGAAAGGUAGCAGCAAUAAA.GCUCAAU.....UAUA.A.G.AGGUGAAU.UCUUCGAGU
#=GR NC_031664.1/23490-23399   PP *********..****....**...**.*****......****.************************************.*******.....****.*.*.********.*********
NC_031168.1/105054-105144         ACUCGGAAG..AUUA....CA...AA.UGUAA......UAGC.UGAAACUUGUCAGAACUGGAAAGUAGCAGCAAUUAG.GCUUCAU.....UACA.A.A.GG-UAUAU.UUUCCGGGU
#=GR NC_031168.1/105054-105144 PP *********..****....**...**.*****......****.************************************.*******.....****.*.9.99.99999.*********
NC_031179.1/113044-112955         ACUCGGAAA..AUUA....UA...UA.UGUAA......UAGC.UAAAAAUUGUCAGGACUGGAAAGUAGCAGCAAUACA.GUUCAAU.....UACA.C.A.GA-UAUU-.UUUCCGGGU
#=GR NC_031179.1/113044-112955 PP *********..****....**...**.*****......****.************************************.*******.....****.9.8.77.**99..*********
NC_031148.1/116830-116740         ACUCGGAAA..UUUU....UA...UA.AGUAA......UAGC.UGAAAGUUGUCAGGACUGGAAAGUAGCAGCAAUAAA.GCUUAAU.....UACA.U.A.GUU-AAAU.UUUCCGGGU
#=GR NC_031148.1/116830-116740 PP *********..****....**...**.*****......****.************************************.*******.....***9.8.8.875.5556.*********
NC_031654.1/23521-23430           GCUCGAAAA..CUUA....CG...AU.CGUAG......UAUC.UUAAAAUUGUCAGGACCGGAAGGUAGCAGCAAAUAA.GAAUUUC.....UACG.A.A.AGGUCAGC.UUUUCGAGC
#=GR NC_031654.1/23521-23430   PP *********..****....**...**.*****......****.************************************.*******.....****.*.*.********.*********
NC_035286.1/9504-9597             GCUCGAAAAauGAUA....CA...AG.AUUAA......UAGC.UAAAACUUGUCAGAACUGAAAAGUAGCAGCAAUUAG.GUUUAAU.....UAAA.A.U.GG-UAUCUuUUUUCGGGU
#=GR NC_035286.1/9504-9597     PP *********9*****....**...**.*****......****.************************************.*******.....***9.8.8.88.****999********
NC_031180.2/200093-200185         GCUCGGGGU..UACA....CA...AU.UAUAG......UAAA.UUAAACUUGUCAGAACUGGAAAGAAGCAGCAAUAAG.AUUUAAU.....UAUA.A.A.UGGUGUAUuAUCCCGAGU
#=GR NC_031180.2/200093-200185 PP *********..****....**...**.*****......****.************************************.*******.....****.*.*.*******88*********
NC_032041.1/19496-19405           GCUCGAAAA.uGAUA....CA...AC.AUUAA......UUAC.UGAAAUUUGUCAGAACUGGAAAGUAGCAGCAAUUAA.GUUAAAU.....UA-U.A.A.UGGUAUCU.UUUUCGGGU
#=GR NC_032041.1/19496-19405   PP *********.*****....**...**.*****......****.************************************.**99996.....66.6.7.7.777*****.*********
NC_031656.1/23385-23293           GCUCGGAGA..UUCA....CA...UC.UGUAA.....gAAAU.UCAAUUUUGUCAGGACUGAAAAGUAGCAGCAAUAAA.ACUUCAU.....UACA.A.U.UGGUGAAU.CCUCCGAGU
#=GR NC_031656.1/23385-23293   PP *********..****....**...**.*****.....99999.9***********************************.*******.....****.*.*.********.*********
NC_031658.1/23552-23461           GCUCGGAGA..UUUA....CA...CU.UAUAG......UAGC.CUAAAGUUGUCAGGACUGGAAAGUAGCAGCAAUACG.GAAUUAU.....UAUA.A.A.AGGUAAGU.UCUUCGAGC
#=GR NC_031658.1/23552-23461   PP *********..****....**...**.*****......****.************************************.*******.....****.*.*.********.*********
NC_021075.1/17898-17807           ACUCGGAAA..GAUA....UA...CA.UAUAA......UAAC.UAAAAUUUGUCAGGACUGGAAAGUAGCAGCAAUUUA.GUUUACU.....UAUA.AuU.AGGUAGC-.UUUCCGGGU
#=GR NC_021075.1/17898-17807   PP *********..****....**...**.*****......****.************************************.*******.....***9.625.789****..*********
NC_031655.1/23413-23322           ACUCGAAGA..UUUA....CA...UU.UAUAG......UAUC.CUAAACUUGUCAGGACUGGAAAGUAACAGCAAUACG.GAAUUAC.....UAUG.A.A.AGGUAAAU.UCUUCGAGU
#=GR NC_031655.1/23413-23322   PP *********..****....**...**.*****......****.************************************.*******.....****.*.*.********.*********
NC_031169.1/101681-101770         GCUCGGAAA..AGCA....CA...UA.UGUAA......CAGC.CAAAUUUUGUCAGAACUGGAAAGUAGCAGCAAUUAA.GCUAAAU.....UACA.U.G.GG-UAAU-.UUUCCGGGU
#=GR NC_031169.1/101681-101770 PP *********..****....**...**.*****......****.************************************.*******.....****.9.9.99.****..*********
NC_040158.1/102304-102394         ACUCGGAAA..GUUA....CA...UA.UGUAA......CAGC.UGAAAUUUGUCAGGACUGGAAAGUAGCAGCAAUCAA.ACUGAAU.....UACA.U.A.GG-UAUAU.UUUCCGGGU
#=GR NC_040158.1/102304-102394 PP *********..****....**...**.*****......****.************************************.*******.....****.*.9.99.*****.*********
NC_036637.1/4532-4623             GCUCGGAAA..AUUA....UG...UA.UGUAA......UAGC.UAAAACUUGUCAGAACUGGAAAGUAGCAGCAACAAA.GCUAAAU.....UGCA.U.A.GUUGUAUU.UUUCCGGGU
#=GR NC_036637.1/4532-4623     PP *********..****....**...**.*****......****.************************************.*******.....****.9.9.88777777.*********
NC_029859.1/100480-100570         ACUCGGAAA..GUUA....CA...UA.UGUAA......CAGC.UAAAAUUUGUCAGGACUGGAAAGUAGCAGCAAUCAA.ACUGAAU.....UACA.U.A.GG-UAUAU.UUUUCGGGU
#=GR NC_029859.1/100480-100570 PP *********..****....**...**.*****......****.************************************.*******.....****.*.9.99.*****.*********
NC_023785.1/4911-4999             ACUCGGAAA..-AUA....CA...UU.AUUAA......CAGC.UUAAAUUUGUCAGAACUGGAAAGUAGCAGCAAUUAA.GCUCAAU.....UAGA.U.A.AGGUAU--.UUUCCGGGU
#=GR NC_023785.1/4911-4999     PP ********6...9**....**...**.*****......****.************************************.*******.....****.*.*.*****9...6********
NC_029858.1/100463-100553         ACUCGGAAA..GUUA....CA...UA.UGUGA......CAGC.UAAAAUUUGUCAGGACUGGAAAGUAGCAGCAAUUAA.ACUGAAU.....UACA.U.A.GG-UAUAU.UUUCCGGGU
#=GR NC_029858.1/100463-100553 PP *********..****....**...**.*****......****.************************************.*******.....****.*.9.99.*****.*********
NC_031145.1/110260-110350         ACUCGGGAA..GUUA....CA...UA.UGUGA......UGGC.UAAAAUUUGUCAGGACUGGAAAGUAGCAGCAAUAAA.GCUAAAU.....UACA.A.A.AGGUAUA-.UUUCCGGGU
#=GR NC_031145.1/110260-110350 PP *********..****....**...**.*****......****.************************************.*******.....****.*.*.****997..*********
NC_039977.1/84091-84000           ACUCGAAAG..GUUA....UA...AA.UAUAU......UAGC.UAAAACUUGUCAGGACUGGAAAGUAGCAGCAAUUUA.GCUAAAU.....UAUAuU.U.AGAUGGC-.UUUUCGGGU
#=GR NC_039977.1/84091-84000   PP *********..****....**...**.*****......****.************************************.*******.....****44.6.789****..*********
NC_020795.1/4845-4935             ACUCAGAAA..AUUA....CA...UG.UGUAA......UAGC.UGAAAUUUGUCAGGACUGGAAAGUAGCAGCAAUUAA.GCUAAAU.....UACA.A.A.AG-UGUAU.UUUCUGGGU
#=GR NC_020795.1/4845-4935     PP *********..999*....**...**.*****......****.************************************.*******.....****.*.*.**.66666.*********
NC_039143.1/24539-24451           ACUCGGAAA..-CUA....CA...UU.AGUAA......UAGC.UUAAACUUGUCAGGACUGGAAAGUAGCAGCAAUUCA.GCUUAAU.....UACA.U.A.AGGUAU--.UUUCCGGGU
#=GR NC_039143.1/24539-24451   PP ********7...79*....**...**.*****......****.************************************.*******.....****.*.*.****97...7********
NC_035298.1/9404-9493             GCUCGAAAU..AAUA....CA...UA.AAUAA......UUGU.UAAAUCUUGUCAGGACUGAAAAGUAGCAGCAAUUAA.CUUAUGA.....UAUU.U.A.GG-UAUU-.GUUUCGGGU
#=GR NC_035298.1/9404-9493     PP *********..****....**...**.*****......****.************************************.*******.....****.*.9.99.****..*********
NC_035283.1/9337-9426             GCUCGAAAU..UAUA....UA...UA.AAUAG......UAAG.UAAAUUUUGUCAGGACUGAAAAGUAGCAGCAAUUAG.CUUAUCU.....UAUU.U.A.GG-UAUU-.AUUUCGGGU
#=GR NC_035283.1/9337-9426     PP *********..****....**...**.*****......****.************************************.*******.....****.9.9.99.***9..*********
NC_035260.1/9497-9589             GCUCGAAAAauAAUA....CA...AA.AUGAA......UAAC.UCAAAUUUGUCAGGACUGAAAAGUAGCAGCAAUAAA.GUUAAAC.....UUUU.U.A.GG-UAUUU.UUUUCGGGU
#=GR NC_035260.1/9497-9589     PP *********99****....**...**.*****......****.************************************.*******.....**99.9.9.88.*****.*********
NC_039140.1/24322-24234           ACUCGGAAA..-AUA....CA...UU.AUUAA......UAGC.UUAAACUUGUCAGAACUGGAAAGUAGCAGCAAUUCA.GCUUAAU.....UAAG.U.A.AAGUAU--.UUUUCGGGU
#=GR NC_039140.1/24322-24234   PP ********6...9**....**...**.*****......****.************************************.*******.....****.*.*.**9**9...6********
NC_031170.1/116721-116810         ACUCGGAAA..AUCA....CA...UC.AGUAA......UAGC.UAAAACUUGUCAGGACUGGAAAGUAGCAGCAACUAA.GCUAAAU.....UACA.A.A.GG-UAUU-.UUUCCGGGU
#=GR NC_031170.1/116721-116810 PP *********..****....**...**.*****......****.************************************.*******.....***9.9.9.88.8777..*********
NC_039144.1/24161-24073           ACUCGGAAA..-CUA....CA...UU.AAUAA......UAGC.UUAAACUUGUCAGGACUGGAAAGUAGCAGCAAUUCA.GCUUAAU.....UAUA.U.A.AGGUAU--.UUUCCGGGU
#=GR NC_039144.1/24161-24073   PP ********7...79*....**...**.*****......****.************************************.*******.....****.*.*.****97...7********
NC_031668.1/23533-23443           GCUCGGGAA..CUAG....CA...AU.UAUAA......AGUU.UUAAACUUGUCAGAACUGGAAAGUAGCAGCAACAAA.AACUUA-.....UAUA.A.A.AGGCUAGU.CUCUCGAGU
#=GR NC_031668.1/23533-23443   PP *********..****....**...**.*****......****.************************************.988888......****.*.*.********.*********
NC_031167.1/4886-4976             ACUCGGAAA..AUUA....CA...CA.UGUAA......UGGC.UGAAAUUUGUCAGGACUGGAAAGUAGCAGCAACUAA.GCUAAAU.....UACA.A.A.GG-UAUAU.UUUUCGGGU
#=GR NC_031167.1/4886-4976     PP *********..****....**...**.*****......****.************************************.*******.....****.*.9.99.99999.*********
NC_032396.1/20396-20305           GCUCGAAAA.uGGUA....CA...GC.AUUAA......UUAC.UGAAAUUUGUCAGGACUGGAAAGUAGCAGCAAUUAA.GUUAAAU.....UA-U.A.A.UGGUAUCU.UUUUCGGGU
#=GR NC_032396.1/20396-20305   PP *********.*****....**...**.*****......****.************************************.**99996.....66.6.7.7.777*****.*********
NC_032399.1/20405-20314           GCUCGAAAA.uGGUA....CA...GC.AUUAA......UUAC.UGAAAUUUGUCAGGACUGGAAAGUAGCAGCAAUUAA.GUUAAAU.....UA-U.A.A.UGGUAUCU.UUUUCGGGU
#=GR NC_032399.1/20405-20314   PP *********.*****....**...**.*****......****.************************************.**99996.....66.6.7.7.777*****.*********
NC_035287.1/9486-9575             GCUCGAAAU..AUUA....CA...UU.AGUAU......UCAA.UAAAUUUUGUCAGGACUGGAAAGUAGCAGCAAUCAU.UUUGUAU.....UACU.U.A.GG-UGUU-.AUUUCGGGU
#=GR NC_035287.1/9486-9575     PP *********..****....**...**.*****......****.************************************.*******.....****.*.9.99.*999..*********
NC_031211.1/18729-18640           ACUCGGAAA..AUUU....AC...AU.AGUAA......UAGC.UUAAAUUUGUCAGAACUGAAAAGUAGCAGCAAUAUA.GCUUAA-.....UAAU.A.C.GGUAUAU-.UUUUCGGGU
#=GR NC_031211.1/18729-18640   PP *********..****....**...**.*****......****.************************************.****99......****.*.*.**99999..*********
NC_031657.1/23391-23296           GCUCGGAUA..UUCAcauaUA...UA.UGUAA......AAAA.AUAAACUUGUCAGGACUGAAAAGUAGCAGCAAUACA.AAUUAAU.....UACA.A.A.UGGUGAAU.UUUCCGAGU
#=GR NC_031657.1/23391-23296   PP *********..****333335...68.99999......****.************************************.*******.....****.*.*.********.*********
NC_031176.2/109485-109390         GCUCGGAAA..UAUG....CA...AU.UAUAAa...uaCAAA.UCUAACUUGUCAGAACUGGAAAGAAGCAGCAAUAAGaCUUUAUU.....UAUA.U.A.UGGCAUUU.AUUCCGAGC
#=GR NC_031176.2/109485-109390 PP *********..****....**...**.****98...778889.999********************************978899999.....****.*.*.********.*********
NC_040156.1/100655-100745         ACUCGGAAA..GUUA....CA...UA.UGUGA......CAGC.UGAAAUUUGUCAGGACUGGAAAGUAGCAGCAAUCAA.ACUGAAU.....UACA.U.A.GG-UAUAU.UUUUCGGGU
#=GR NC_040156.1/100655-100745 PP *********..****....**...**.*****......****.************************************.*******.....****.*.9.99.*****.*********
NC_035273.1/9346-9435             GCUCGAAAU..AAUA....UA...UA.AAUAG......CAAG.UGAAUUUUGUCAGGACUGAAAAGUAGCAGCAAUUAG.CUUAUUC.....UAUU.U.A.GA-UAUU-.GUUUCGGGU
#=GR NC_035273.1/9346-9435     PP *********..****....**...**.*****......****.************************************.*******.....****.9.8.77.****..*********
NC_039092.1/24448-24360           ACUCGGAAA..-AUA....CA...UU.AUUAA......UAGC.UUAAACUUGUCAGGACUGGAAAGUAGCAGCAAUUCA.GCGUAAU.....UAUA.U.A.AGGUAU--.UUUCCGGGU
#=GR NC_039092.1/24448-24360   PP ********6...999....**...**.*****......****.************************************.*******.....****.*.*.**9999...6********
NC_029860.1/102624-102712         ACUCGGAAA..-AUA....CA...UU.AUUAA......UAGC.UCAAACUUGUCAGGACUGGAAAGUAGCAGCAAUUCA.GCUUAAU.....UAUA.U.A.AGGUGU--.UUUCCGGGU
#=GR NC_029860.1/102624-102712 PP ********6...899....99...99.99***......****.************************************.*******.....****.*.*.**9999...6********
NC_035278.1/9339-9428             GCUCGAAAU..AAUA....UA...UA.AAUAA......UAAG.UCAACUUUGUCAGAACUGAGAAGUAGCAGCAACUAA.CUUAUUU.....UAUU.U.A.GA-UAUU-.AUUUCGGGU
#=GR NC_035278.1/9339-9428     PP *********..****....**...**.*****......****.************************************.*******.....****.9.8.77.****..*********
NC_037841.1/58010-57922           GCUCGGAAA..ACUA....CA...C-.AGUAA......UAGC.UCAAACUUGUCAGAACUGGAAAGUAGCAGCAAUAUA.GCUAAAU.....UACA.U.A.GG-UAUG-.UUUCCGGGU
#=GR NC_037841.1/58010-57922   PP *********..8888....76...5..79***......****.************************************.*******.....**98.8.8.77.7765..*********
NC_021618.1/58016-57928           GCUCGGAAA..ACUA....CA...C-.AGUAA......UAGC.UCAAACUUGUCAGAACUGGAAAGUAGCAGCAAUAUA.GCUAAAU.....UACA.U.A.GG-UAUG-.UUUCCGGGU
#=GR NC_021618.1/58016-57928   PP *********..8888....76...5..79***......****.************************************.*******.....**98.8.8.77.7765..*********
NC_035261.1/9459-9547             GCUCGAAAU..UGUA....CA...AA.AAUAU......UAAA.UAAAGUUUGUCAGGACUGAAAAGUAGCAGCAAUUAU.UUUAUAU.....UAUU.U.A.GG-UAC--.UUUUCGGGU
#=GR NC_035261.1/9459-9547     PP *********..****....**...**.*****......****.************************************.*******.....****.9.9.99.**9...7********
NC_035276.1/9417-9507             GCUCGAAAC..AAUA....UA...UA.AAUAC......UCAA.UAAAUCUUGCCAGGACUGAAAAGUAGCAGCAAUAAAaCUUGUAG.....UAUU.U.G.GG-UAUU-.GUUUCGGGU
#=GR NC_035276.1/9417-9507     PP *********..****....**...**.*****......****.***********************************978888999.....****.9.9.99.****..*********
NC_031665.1/22736-22645           ACUCGGGAA..CUAG....CA...UU.UACAA......UAAC.UUGACAUUGUCAGGACUGGAAAGUAGCAGCAAACAA.GAUAUAU.....UGUA.A.A.AGGCAAGC.UUUUCGAGU
#=GR NC_031665.1/22736-22645   PP *********..****....**...**.*****......****.************************************.*******.....****.*.*.********.*********
NC_038100.1/103753-103841         ACUCGGGAA..-CUA....CA...UU.AAUAA......UAGC.UUAAACUUGUCAGGACUGGAAAGUAGCAGCAAUUCA.GCUUAAU.....UAUA.U.A.AGGUAU--.UUUCCGGGU
#=GR NC_038100.1/103753-103841 PP ********7...79*....**...**.*****......****.************************************.*******.....****.*.*.****97...7********
NC_039142.1/23799-23710           ACUCGGAAA..C-AA....CA...UU.AUUAA......UAGC.UUAAACUUGUCAGGACUGGAAAGUAGCAGCAAUUCA.GCUUAAU.....UAUA.U.A.AGGUUUU-.UUUCCGGGU
#=GR NC_039142.1/23799-23710   PP ********9..4.44....55...66.6699*......****.************************************.*******.....**99.9.9.9977665..9********
NC_026523.1/140137-140048         GCUCGGAAU..UAUA....UG...UA.AAUAG......UAAG.UGAAUUUUGUCAGGACUGAAAAGUAGCAGCAAUUAA.CUUAUUU.....UAUU.U.G.GA-UAUU-.AUUCCGGGU
#=GR NC_026523.1/140137-140048 PP *********..****....**...**.*****......****.************************************.*******.....***9.8.7.77.***9..*********
NC_031174.1/113216-113128         ACUCGAAAA..AUUU....AC...AU.AGUAA......UAGC.UUAAUUUUGUCAGAACUGAAAAGUAGCAGCAAUAGA.GCUUAA-.....UAAU.A.G.GU-AUAU-.UUUUCGGGU
#=GR NC_031174.1/113216-113128 PP *********..9999....99...99.9****......****.************************************.****99......9988.7.6.65.7777..*********
NC_029644.1/177638-177550         ACUCGGAAA..-CUA....CA...UU.AAUAA......UAGC.UUAAAUUUGUCAGGACUGGAAAGUAGCAGCAAUUCA.GCUUAAU.....UACA.U.A.AGGUAU--.UUUCCGGGU
#=GR NC_029644.1/177638-177550 PP ********7...799....**...**.*****......****.************************************.*******.....****.*.*.***997...7********
NC_031149.1/104004-104092         ACUCGGAAA..-CUA....CA...UU.AAUAA......UAGC.UUAAAUUUGUCAGGACUGGAAAGUAGCAGCAAUUCA.GCUUAAU.....UACA.U.A.AGGUAU--.UUUCCGGGU
#=GR NC_031149.1/104004-104092 PP ********7...799....**...**.*****......****.************************************.*******.....****.*.*.***997...7********
NC_038144.1/17087-16995           GCUCGGAAA.aAUUA....UA...AG.UAUAA......ACAU.UCUAUAUUGUCAGGACUGGAAAGUAGCAGCAAUUUU.AGUGAAU.....UAUA.U.U.UGUUUAGA.UUUCCGGGC
#=GR NC_038144.1/17087-16995   PP ********9.9****....**...**.*****......****.************************************.*******.....****.*.*.********.*********
NC_035268.1/8921-9009             GCUCGAAAU..UAUU....CA...UA.AAUAA......UCAC.UAAAUUUUGUCAGGACUGAAAAGUAGCAGCAAUUCA.GUUGUAU.....UAUU.U.A.---GGUAU.UUUUCGAGU
#=GR NC_035268.1/8921-9009     PP *********..****....**...**.*****......****.************************************.****999.....9999.8.8....66778.9********
NC_031659.1/23654-23565           ACUCGAGAG..CUAA....CG...AU.UAUAA......CGUU.UUCAACUUGUCAGGACUGAAAAGUAGCAGCAAAAAA.A--UAGC.....UAUA.A.A.AGGUUAGU.CUUUCGAGU
#=GR NC_031659.1/23654-23565   PP *********..****....**...**.*****......****.*******************************99887.6..5778.....****.*.*.********.*********
NC_039139.1/24707-24619           ACUCGGAAA..-AUA....CA...UU.AUUAA......UAGC.UUAAACUUGUCAGGACUGGAAAGUAGCAGCAAUUCA.GCUCAAU.....UACA.U.A.AGGUGU--.UUUCCGGGU
#=GR NC_039139.1/24707-24619   PP ********6...899....99...99.99***......****.************************************.*******.....****.*.*.**9999...6********
NC_031666.1/23373-23282           ACUCGGAGG..UCUA....CA...CU.UAUAG......UAGC.CGAACAUUGUCAGAACCGGAGGGUAGCAGCAAUAGG.AAACACU.....UAUA.A.U.AGGUAUAC.UCUCCGAGU
#=GR NC_031666.1/23373-23282   PP *********..****....**...**.*****......****.************************************.*******.....****.*.*.********.*********
NC_039141.1/24764-24676           ACUCGGAAA..-AUA....CA...UU.AUUAA......UAGC.UUAAACUUGUCAGGACUGGAAAGUAGCAGCAAUUCA.GCUUAAU.....UACA.U.A.AGGUGU--.UUUUCGGGU
#=GR NC_039141.1/24764-24676   PP ********6...899....99...99.99***......****.************************************.*******.....****.*.*.**9999...6********
NC_035291.1/9448-9537             GCUCGAAAU..UAUA....CA...UA.AGUAA......UCAG.UAAAGUUUGUCAGGACUGAAAAGUAGCAGCAAUUAA.CUUGUAU.....UAUU.U.A.GG-UAUU-.CUUUCGGGU
#=GR NC_035291.1/9448-9537     PP *********..****....**...**.*****......****.************************************.*******.....****.9.9.99.***9..9********
NC_035292.1/9331-9420             GCUCGAAUU..UAUA....UA...UA.GAUAA......UAAG.UUAAUCUUGCCAGAACUGAAGAGUAGCAGCAAUAAU.CUUAUAU.....UAUU.U.A.GG-UAUA-.UUUUCGGGU
#=GR NC_035292.1/9331-9420     PP *********..****....**...**.*****......****.************************************.*******.....****.9.9.99.****..99*******
NC_031144.1/131340-131249         GCUCGGAAA..ACUA....CA...UU.AAUAA......UAGC.UGAAACAUGUCAGAACUGGAAAGUAGCAGCAAUAAG.CUAAAAU.....UAUA.U.A.UAAGGUAU.UUUCCGGGU
#=GR NC_031144.1/131340-131249 PP *********..9999....99...**.*****......****.************************************.9999999.....8888.7.7.76433335.9********
NC_039978.1/169160-169069         CCUCGGAAA..GUUA....UA...CA.UAUAA......UUAC.UAUAACUUGUCAGAACUAAAAAGUAGCAGCAAAAUA.GGUUAAU.....UAUAuU.U.AGGUGAC-.UUUCCGGGU
#=GR NC_039978.1/169160-169069 PP *********..****....**...**.*****......****.************************************.*******.....****44.5.789****..*********
NC_035285.1/9403-9494             GCUCGAAAU..UAUA....CA...AUuUAUUA.....aUAGC.UUAAAUUUGUCAGGACUGGAAAGUAGCAGCAAUAUA.GCUAAAC.....AGUA.U.A.-GGUAUA-.UUUUCGGGU
#=GR NC_035285.1/9403-9494     PP *********..9999....98...74244444.....5****.************************************.***9997.....6665.5.5..55****..9********
NC_031660.1/23732-23641           ACUCGAAAG..CUAG....CG...UU.UAUAA......UAUC.UUGAAAUUGUCAGGACUGGAAAGUAGCAGCAAACAA.GAAAUAU.....UGUA.A.A.AGGCCAGU.AUUUCGAGU
#=GR NC_031660.1/23732-23641   PP *********..****....**...**.*****......****.************************************.*******.....****.*.*.********.*********
NC_039145.1/26285-26194           ACUCGGGAA.cUUGC....GU...UA.AUUAA......CAGC.UAAAACUUGUCAGAACUGGGAAGUAGCAGCAAUAAA.GCUUAAU.....UAUA.C.A.AG-GCAUU.UUCCCGGGU
#=GR NC_039145.1/26285-26194   PP *********.65555....56...66.66788......****.************************************.******9.....9988.8.8.88.55566.*********
NC_035263.1/7451-7542             GCUCGAAAA.cGAUA....UA...AU.AUUUA.....uAACU.UAUAAUUUGUCAGAACUGAAAAGUAGCAUCAAUAUA.AGUUAA-.....AUGU.A.A.UGAUAUC-.UUUUCGGGU
#=GR NC_035263.1/7451-7542     PP *********.*****....**...**.**999.....45566.788*********************************.999999......899*.*.*.*******..*********
NC_035284.1/9513-9602             GCUCGAAAU..AAUA....CG...UA.AAUAA......UUAG.UGAAUUUUGUCAGAACUGAGAAGAAGCAGCAAUUAA.CUUUUAU.....UAUU.U.A.GG-UAUU-.UUUUCGGGU
#=GR NC_035284.1/9513-9602     PP *********..****....**...**.*****......****.************************************.*******.....****.*.9.99.****..9********
NC_035266.1/9368-9456             GCUCGAAAU..AGUA....CA...GA.AAUAG......UCAA.UAAAUUUUGUCAGAACUGAAAAGUCGCAGCAAUAAU.AUUGUAU.....UAUU.U.A.GG-UAUU-.-UUUCGAGU
#=GR NC_035266.1/9368-9456     PP *********..****....**...**.*****......****.************************************.*******.....****.*.9.99.**85...9*******
NC_035257.1/9433-9525             GCUCGAAAAauGAUA....UA...G-.CAUAA.....aAAAU.UAAAAUUUGUCAGGACUGAAAAGUAGCAGCAGCUAA.UUCUAAU.....UAUG.U.A.GG-UAUCU.UUUUCGGGU
#=GR NC_035257.1/9433-9525     PP *********966666....66...5..66666.....49999.99*********************************9.9999996.....6666.6.6.66.66666.*********
NC_035293.1/10805-10894           GCUCGGAAU..UAUA....CA...GA.AGUAA......UCAA.UGAAUUUUGUCAGGACUGAAAAGUAGCAGCAAUUAA.CUUGUAU.....UAUU.U.G.GG-UAUU-.UUUUCGGGU
#=GR NC_035293.1/10805-10894   PP *********..****....**...**.*****......****.************************************.*******.....****.9.9.99.***9..9********
NC_031662.1/23796-23705           ACUCGGAGA..UUUA....CA...UUaUGUAA......UUAA.UGAAAAUUGUCAGGACUGGAAAGUAGCAGCAAUAAA.AACUAU-.....UACA.G.A.AGGUUAAU.ACUCCGAGU
#=GR NC_031662.1/23796-23705   PP *********..9999....88...64399999......****.************************************.999887......9999.9.9.99999999.*********
NC_006137.1/4964-5052             ACUCGGGAA..-AUA....CG...UU.AUUAA......UAGC.UUAAACUUGUCAGGACUGGAAAGUAGCAGCAAUUCA.GCUUAAU.....UACA.U.A.AGGUGU--.UUUCCGGGU
#=GR NC_006137.1/4964-5052     PP ********6...888....89...99.99***......****.************************************.*******.....****.*.*.*99988...6********
NC_031161.1/19409-19317           GCUCGAAAAauGAUU....CA...AA.AUUAA......UGAC.UUAAACUUGUUAGAACUGAAAAGUAGCAACAAUAUA.GUCAGA-.....UAAU.U.A.AGGUAUCU.UUUUCGGGU
#=GR NC_031161.1/19409-19317   PP **********99999....99...99.99999......****.**********************************99.988777......8999.9.9.99999999.*********
NC_035296.1/9231-9320             GCUCGGAAU..UUUA....UA...UA.AAUAG......AGUA.UUAAUUUUGUCAGGACUGAAAAGUAGCAGCAAUAAA.CUAAUUC.....UAUU.U.A.GG-UACU-.AUUUCGGGU
#=GR NC_035296.1/9231-9320     PP *********..****....**...**.*****......****.************************************.*******.....****.*.9.99.**99..*********
NC_035288.1/9447-9536             GCUCGAAAU..UGUA....CG...UA.GAUAG......UAAG.UGAAGUUUGUCAGGACUGAAAAGUAGCAGCAACAAU.CUUAUAU.....UAUU.G.A.GG-UAUU-.AUUUCGGGU
#=GR NC_035288.1/9447-9536     PP *********..****....**...**.*****......****.************************************.*******.....****.9.9.99.***9..*********
NC_035272.1/9304-9393             GCUCGAAAU..UUUA....CG...UA.AAUAG......UUUG.UGAAUCUUGCCAGAACUGAAAAGUAGCAGCAAUUAA.CUUAUUC.....UAUU.U.A.GG-UAUU-.AUUUCGGGU
#=GR NC_035272.1/9304-9393     PP *********..****....**...**.*****......****.************************************.*******.....****.*.9.99.**99..*********
NC_035267.1/9337-9426             GCUCGAAAU..UUUA....UA...UA.GAUGA......UAAG.UGAAGUUUGUCAGAACUGAAAAGUAGCAGCAAUUAA.CUUACAU.....AAUC.U.A.GG-UAUU-.UUUUCGGGU
#=GR NC_035267.1/9337-9426     PP *********..****....**...**.*****......****.************************************.*******.....****.9.9.99.**99..9********
NC_035274.1/9462-9551             GCUCGAAAU..CUUA....UG...UA.AAUAG......UAUG.UGAAUUUUGUCAGGACUGAAAAGUAGCAGCAAUUAG.CUUAUUC.....UAUU.U.A.GA-UAUU-.AUUUCGGGU
#=GR NC_035274.1/9462-9551     PP *********..****....**...**.*****......****.************************************.*******.....****.9.8.77.**98..*********
NC_033877.1/164870-164958         ACUCGGAAA..-AUA....CG...UU.AUUAA......UAGC.UUAAACUUGUCAGGACUGGAAAGUAGCAGCAAUAUA.GCUCAAU.....UCUA.U.A.AGGUGU--.UUUCCGGGU
#=GR NC_033877.1/164870-164958 PP ********8...555....56...66.66699......****.************************************.******9.....9888.8.8.887777...8********
NC_038051.1/181541-181453         ACUCGGAAA..-AUA....CG...UU.AUUAA......UAGC.UUAAACUUGUCAGGACUGGAAAGUAGCAGCAAUAUA.GCUCAAU.....UCUA.U.A.AGGUGU--.UUUCCGGGU
#=GR NC_038051.1/181541-181453 PP ********8...555....56...66.66699......****.************************************.******9.....9888.8.8.887777...8********
NC_035290.1/9400-9488             GCUCGAAAU..UUUA....CA...UA.AGUAA......UAAG.UGAAGUGUGUCAGGACUGAAAAGUAGCAGCAAUUAA.CUUUUAU.....UAUU.U.A.GG-UAU--.UUUUCGGGU
#=GR NC_035290.1/9400-9488     PP *********..****....**...**.*****......****.************************************.*******.....****.9.9.99.885...8********
NC_035279.1/9368-9458             GCUCGAAAU..UGUA....CA...UA.GGUAA.....cAUGU.UGAAGUUUGUCAGAACUGAAAAGUAGCAGCAAUAAA.CGAAUGC.....UACU.U.A.GG-UAUA-.UUUUCGGGU
#=GR NC_035279.1/9368-9458     PP *********..****....**...**.*****.....77788.9999********************************.999999*.....****.9.9.99.****..9********
NC_035281.1/9304-9396             GCCCGAAAU..UAUA....UAuaaUA.AAUAG......UAUA.UGAAUUUUGUCAGGACUGAAAAGUAGCAGCAAUCAA.UAUAUUC.....UAUU.U.A.GA-UAUU-.AUUUCGGGU
#=GR NC_035281.1/9304-9396     PP *********..6666....5411135.66777......****.************************************.9999999.....9988.7.7.65.8988..*********
NC_035289.1/9323-9412             GCUCGAAAU..AACA....CU...AA.AAUAA......UUAA.UUAAGAUUGUCAGGACUGAAAGGUAGCAGCAAGUAA.GUUAUGU.....UAUC.U.A.GG-UGUU-.UUUUCGGGU
#=GR NC_035289.1/9323-9412     PP *********..****....**...**.*****......****.************************************.*******.....***9.9.8.88.****..9********
NC_035265.1/7412-7504             GCUCGAAAAauGAUA....UG...AU.ACCUA......CAAC.UUUAAUUUGUCAGGACUGGAAAGUAGCAUCAAUAUA.AGUUAC-.....GUGU.G.U.AGAUAUCU.UUUUCGGGU
#=GR NC_035265.1/7412-7504     PP ***************....**...**.*****......****.************************************.999887......8899.9.9.********.*********
NC_035275.1/9544-9638             GCUCGAAAU..UAUA....CA...UA.UAUAAua.aguUAGA.UAAAGUUUGUCAGAACUGAAAAGUAGCAGCAAUUAA.CUUGUAU.....UAUU.U.A.AG-UAUU-.UUUUCGGGU
#=GR NC_035275.1/9544-9638     PP *********..****....99...87.7777766.6999999.************************************.*******.....****.*.*.*9.***9..9********
NC_035277.1/9194-9283             GCUCGAAAU..UUUA....UA...UA.AAUAG......UUUG.UGAAUUUUGUCAGGACUGAAAAGUAGCAGCAAUAAA.CUUAUUC.....UAUU.U.A.GG-UAUC-.ACUUCGGGU
#=GR NC_035277.1/9194-9283     PP *********..****....**...**.*****......****.************************************.*******.....****.9.9.99.*998..*********
NC_031663.1/23729-23638           GCUCGGAGA..UUAA....CG...CU.UACAG......UAGC.UUGAAAUUGUCAGAACUGGAAGGUAGCAGCAAACAA.GACAUAU.....UGUA.A.A.AGGUCAGU.CUUCCGAGU
#=GR NC_031663.1/23729-23638   PP *********..****....**...**.*****......****.************************************.*******.....****.*.*.********.*********
NC_035297.1/9332-9420             GCUCGAAAU..UUUA....CA...UA.GAUCA......UAAG.UGAAUUUUGUCAGAACUGAAAGGUAGCAGCAAUUAA.CUUAUCU.....GAUU.U.A.GG-UAU--.UUUUCGGGU
#=GR NC_035297.1/9332-9420     PP *********..****....**...**.*****......****.************************************.*******.....****.9.9.99.885...8********
NC_035259.1/9388-9478             GCUCGGAAU..AAUA....CG...UA.AAUAU.....aCUUG.UAAACUUUGUCAGGACUGAAAAGUAGCAGCAAUACA.UUUGUAA.....UAUU.U.A.GG-UAUU-.UUUUCGGGU
#=GR NC_035259.1/9388-9478     PP *********..****....**...**.****9.....67778.999999******************************.*******.....****.9.9.99.****..9********
NC_031661.1/23565-23475           ACUCGGAAG..CUAG....CA...UU.UAUAC......AACC.UUGAAAUUGUCAGGACUGGAAAGUAGCAGCAAACAA.GGAAU-G.....UAAA.A.A.AGGCUAGU.AUUUCGAGU
#=GR NC_031661.1/23565-23475   PP *********..****....**...**.*****......****.********************************9999.87655.5.....****.*.*.********.*********
NC_035294.1/12744-12833           GCUCGGAAU..AAUA....CG...CA.AAAAC......UAAC.UGAAUUUUGUCAGGACUGAAAAGUAGCAGCAAUUAA.GUUAUA-.....UAGU.U.U.GGGUAUU-.UUUUCGGGU
#=GR NC_035294.1/12744-12833   PP *********..****....98...88.88888......899*.************************************.*99988......5666.6.6.666****..9********
NC_035280.1/9165-9257             GCUCGAAAAacGAUA....UG...AU.AUUGA......UGAU.UUAUAUUUGUCAGGACUGAAGAGUAGCAGCAGAUA-.AAUCAAU.....UAAU.A.A.AGUUAUCU.UUUUCGGGU
#=GR NC_035280.1/9165-9257     PP ***************....**...**.*****......****.******************************97655..5677888.....****.*.*.********.*********
NC_035264.1/9409-9496             GCUCGAAU-..UAUG....CA...UA.GAUAA......UCGC.UAAAUUUUGUCAGGACUGAAAAGUAGCAGCAAUUCA.GUUGUAU.....UAUU.U.A.GG-UAUU-.-UUUCGAGU
#=GR NC_035264.1/9409-9496     PP *****997...7999....9*...**.*****......****.************************************.*******.....****.9.9.99.8986...69******
NC_029857.1/117980-117896         GCUCGGAAA..GUUA....AA...UA.UAUAA......UACC.UAGAAAUUGUCAGGACUGGAAGGUAGCAGCAGUAAG.GUUU---.....UAUG.G.-.---UAUUA.UUUCCGAGU
#=GR NC_029857.1/117980-117896 PP *********..****....**...**.*****......****.************************************.*987........8887.5......99999.*********
NC_035282.1/9404-9498             GCUCGGAAC..AAUC....AU...AU.AAUAGua.aguUUCA.UUAAGUUUGUCAGGACCGAAAGGUAGCAGCAAUUAA.CUUGUAC.....UAUU.U.U.AA-GAUU-.GUUUCGGGU
#=GR NC_035282.1/9404-9498     PP *********..****....**...**.****955.5555557.8899********************************.*******.....****.9.9.99.****..*********
NC_035269.1/7496-7594             GCUCGAAAGacGAUA....CA...AA.AAAUA......UAAC.UUUAAUUUGUCAGGACUGGAAAGUAGCAUCAAUAUAaGUUAAAUauauuAUUU.A.A.G-GUAUCU.UUUUCGGGU
#=GR NC_035269.1/7496-7594     PP ***************....**...99.99999......****.*********************************99855544444443325555.5.5.5.5*****.*********
NC_030338.1/18659-18563           ACUCGGAAA..CUUA....CA...AU.UACAUuaauaaUAGC.UAAAAUAUGUCAGGACUGGAAAGUAGCAGCAAUAAG.CUAAAUU.....UUAA.A.A.AGGUAUU-.UUUCCGGGU
#=GR NC_030338.1/18659-18563   PP *********..8888....77...75.44443333477999*.***********************************9.9999999.....9999.9.9.9999998..*********
NC_035271.1/9459-9547             GCUCGAAAU..GAUA....CA...UA.AAUGA......UCAG.UGAAGUUUGUCAGGACUGAAAAGUAGCAGCAAUUAA.CUUGUAU.....UAUU.U.A.GG-UGC--.UUUUCGGGU
#=GR NC_035271.1/9459-9547     PP *********..****....**...**.*****......****.************************************.*******.....****.9.9.99.873...8********
NC_035262.1/9826-9920             GCUCGGAAU..UUUA....CA...UG.AAUAAaa.uaaUUAA.UAAAUUUUGUCAGGACUGAAGAGUAGCAGCAAUUAA.CUUGUUU.....UAUU.U.U.GG-UAUU-.AUUUCGGGU
#=GR NC_035262.1/9826-9920     PP *********..****....99...98.8888866.6557788.99**********************************.*******.....****.9.9.99.**99..*********
NC_035270.1/9248-9338             GCUCGAAAC..UGUA....AU...UA.UUUGU.....uAGGU.AAGACUUUGUCAGAACUGGAAAGUAGCAGCAAUUUA.UUUAUCA.....AUUA.U.A.UG-UAUA-.GUUUCGGGU
#=GR NC_035270.1/9248-9338     PP *********..9999....55...44.44444.....26667.778999***************************988.6666555.....5555.5.5.55.9999..*********
NC_035350.1/12745-12655           GCCCGGGUA..UAUA....UU...UU.UAUAA......UAAA.UUUAAAUUGUCAGAUUUGAAAAAAAGCAGCAAAUA-.AUUUAAU.....UAUA.A.U.UAGUAUGU.AUACCGGGU
#=GR NC_035350.1/12745-12655   PP ******999..****....**...**.*****......****.*******************************9766..5777888.....****.*.*.*******9.999******
NC_023133.1/54817-54906           GCCUGAUA-..CAUA....UG...AU.UGAAA......UAAU.UCGAACUUGCCAGGACUGGAAAGUAGCAACAAGAGA.AUUAAUU.....UAUA.A.U.UGGUAUGU.-UAUCAGGC
#=GR NC_023133.1/54817-54906   PP ***99887...****....**...**.*****......****.************************************.*******.....****.*.*.*******7..78899***
NC_035299.1/9289-9377             GCUCGAAUU..UACA....UA...A-.AAUAC......UAAG.UAAAUUUUGUCAGGACUGAGAAGUAGCAGCAAUUCA.CUUAUAU.....GAUU.U.A.CA-UGUA-.AUUUCGGGU
#=GR NC_035299.1/9289-9377     PP ********9..9999....98...8..78899......****.************************************.9999998.....8888.7.7.77.9999..9********
NC_035295.1/7795-7888             GCUCGAAGAauGAUA....CA...AU.AUAUU......UGAU.CUUAUUUUGUCAGGACUGAAAAGUAGCAGCAAUAUU.AUCAUA-.....GUAA.U.A.AGGUAUCCuUUUUCGGGU
#=GR NC_035295.1/7795-7888     PP *******98899999....99...99.98888......9999.9********************************999.877776......6777.7.7.77799998799*******
NC_031175.1/42408-42317           AGCCUAAUA..CAUA....UA...AU.UAAAU......UAAU.UCGAUCUUGCCAGGACUGGAAAGUAGCAACAAGAGA.AUUGACU.....AUUA.A.U.UGGUAUGU.UAUUAGGCC
#=GR NC_031175.1/42408-42317   PP *********..****....**...**.*****......****.************************************.*******.....****.*.*.********.*********
NC_031667.1/23562-23471           ACUCGGAAG..CUAG....CA...UU.UAUGC......ACCC.UUGAAAUUGUCAGGACUGGAAAGUAGCAACAAACAA.GGAAUGU.....AAAA.A.A.AGGCUAGU.AUUUCGAGU
#=GR NC_031667.1/23562-23471   PP *********..****....**...**.*****......****.************************************.9999999.....9***.*.*.********.*********
NC_031146.1/108875-108784         AUUUAUAGA..UUUG....UA...AU.UAUCA......UAGC.CAGAAUUUGUCAGAACUGGAAGGUAGCAGCAUCACUaGCUUAAU.....AAUA.U.U.AAGCAAA-.UCUAUAAGU
#=GR NC_031146.1/108875-108784 PP *********..****....**...**.*****......****.**********************************999*******.....****.*.*.*******..*********
NC_031177.1/120055-119964         ACUUGUAGG..UUUG....UA...AC.UAUAA......ACAC.UAUAAUUUGUCAGAACUGGGAAGGAGCAGCUAUACUaGUUAAAA.....UAUA.U.U.GAGCAAA-.UCUGUAAGU
#=GR NC_031177.1/120055-119964 PP *********..****....**...**.*****......****.**********************************9989999***.....****.*.*.*******..*********
NC_011087.1/84177-84268           ACUCGGAUC..UACG....CG...GC.UCUGA......CACC.UAAAUUUGGUCAGGACCGGAAGGGAGCAGCCAUAAA.GGAUGCU.....CAAG.G.C.AGGCGUAG.GCUCCGGGU
#=GR NC_011087.1/84177-84268   PP *********..****....**...**.*****......****.************************************.*******.....****.*.*.********.*********
NC_034776.1/216234-216143         GCUCGGGUAuaUAUA....UU...CU.UGUAG......UAAC.UUUAAUAUGUCAGAUUUGAAAAAAAGCAGCA--UAA.GUCUAAC.....UACA.A.A.UGAAAUAU.ACUCCGGGU
#=GR NC_034776.1/216234-216143 PP ******998445555....66...79.*****......****.******************************8..577.8999999.....****.*.*.***99998.889******
NC_035258.1/9549-9641             GCCCGAUAAcuAAUG....CA...UU.GUCUA......GCAU.UUCAUUUUGUCAGGACCGAAAGGUAGCAGUCAUAAA.AUGUAA-.....AAAC.A.A.AGGCUUCU.UUUUCGGGC
#=GR NC_035258.1/9549-9641     PP *********866666....66...66.66678......9999.************************************.*99999......5566.6.6.66678888.*********
NC_035721.1/61139-61051           UCCUGGUUA..CAUA....UA...AA.UGUAU......UGUU.UAAACUUUGUCAGGACUGGAAAGUAGCAGCAAAAGC.AAUU---.....CACA.U.U.UAUGGUGU.AAUCCAGGU
#=GR NC_035721.1/61139-61051   PP *****9866..6666....66...66.66666......9999.********************************9998.7776........4555.5.5.55555555.5679*****
#=GC SS_cons                      <<<<<<<<<..<<<<....--...--.<<<<-......----.--------------<<<____>>>------------.-------.....>>>>.-.-.--->>>>-.>>>>>>>>>
#=GC RF                           gCcCGaAAa..caua....uA...Au.ugUaa......uAac.UaAAucUUGUCAGAACuGgAAaGUAGCAGCAAUAAa.guUUAau.....uAca.a.a.aGauaugu.uUUuCGgGc
//


#for motif_8813
# STOCKHOLM 1.0

NC_029134.1/77347-77394                    UUUCUAGAUCAAGGAGGUUGA..AUAU......AU.............................UCAACCUCCUUGAUCUAGAAA
#=GR NC_029134.1/77347-77394            PP *********************..****......**.............................*********************
NC_029134.1/77394-77347                    UUUCUAGAUCAAGGAGGUUGA..AUAU......AU.............................UCAACCUCCUUGAUCUAGAAA
#=GR NC_029134.1/77394-77347            PP *********************..****......**.............................*********************
NC_029856.1/77025-77072                    UUUCUAGAUCAAGGAUAUUGA..AUAA......AU.............................UCAAUAUCCUUGAUCUAGAAA
#=GR NC_029856.1/77025-77072            PP *********************..****......**.............................*********************
NC_032288.1/117052-117099                  UUUCUAGAGUAAUGAGACUAA..AUAG......AU.............................UUAGUCUCAUUACUCUAGAAA
#=GR NC_032288.1/117052-117099          PP *********************..****......**.............................*********************
NC_018523.1/77161-77206                    UUUCGAAAUUUCCAAGGUUCU..CUAG......--.............................AGAACCUUGGAAAUUUCGAAA
#=GR NC_018523.1/77161-77206            PP *********************..****.....................................*********************
NC_018523.1/77206-77161                    UUUCGAAAUUUCCAAGGUUCU..CUAG......--.............................AGAACCUUGGAAAUUUCGAAA
#=GR NC_018523.1/77206-77161            PP *********************..****.....................................*********************
NC_028502.1/77024-77069                    UUUCGAAAUUUCCAAGGUUCU..CUAG......--.............................AGAACCUUGGAAAUUUCGAAA
#=GR NC_028502.1/77024-77069            PP *********************..****.....................................*********************
NC_028502.1/77069-77024                    UUUCGAAAUUUCCAAGGUUCU..CUAG......--.............................AGAACCUUGGAAAUUUCGAAA
#=GR NC_028502.1/77069-77024            PP *********************..****.....................................*********************
NC_028503.1/77145-77190                    UUUCGAAAUUUCCAAGGUUCU..CUAG......--.............................AGAACCUUGGAAAUUUCGAAA
#=GR NC_028503.1/77145-77190            PP *********************..****.....................................*********************
NC_028503.1/77190-77145                    UUUCGAAAUUUCCAAGGUUCU..CUAG......--.............................AGAACCUUGGAAAUUUCGAAA
#=GR NC_028503.1/77190-77145            PP *********************..****.....................................*********************
NC_029856.1/77072-77025                    UUUCUAGAUCAAGGAUAUUGA..AUUU......AU.............................UCAAUAUCCUUGAUCUAGAAA
#=GR NC_029856.1/77072-77025            PP *********************..****......**.............................*********************
NC_032288.1/117099-117052                  UUUCUAGAGUAAUGAGACUAA..AUCU......AU.............................UUAGUCUCAUUACUCUAGAAA
#=GR NC_032288.1/117099-117052          PP *********************..****......**.............................*********************
NW_017244425.1/18465-18515                 UUUCCCAAUAUUGAAUAUUCC..AUAUa...auAU.............................GGAAUAUUCAAUAUUGGGAAA
#=GR NW_017244425.1/18465-18515         PP *********************..99999...9***.............................*********************
NW_017244425.1/18515-18465                 UUUCCCAAUAUUGAAUAUUCC..AUAUu...auAU.............................GGAAUAUUCAAUAUUGGGAAA
#=GR NW_017244425.1/18515-18465         PP *********************..*****...****.............................*********************
NW_006921800.1/5775272-5775323             UUUCCAAAGCUGGAAGAUUUU..AUAUau..auAU.............................AUAAUCUUCCAGCUUUGGAAA
#=GR NW_006921800.1/5775272-5775323     PP *********************..999999..999*.............................*********************
NC_036461.1/27413823-27413774              UUUAGAAAUGUUUCAGAUUGU..CUAUc....aAG.............................ACAAUCUGAAACAUUUCUAAA
#=GR NC_036461.1/27413823-27413774      PP *********************..***99....9**.............................*********************
NW_019369745.1/27413823-27413774           UUUAGAAAUGUUUCAGAUUGU..CUAUc....aAG.............................ACAAUCUGAAACAUUUCUAAA
#=GR NW_019369745.1/27413823-27413774   PP *********************..***99....9**.............................*********************
NW_019384576.1/214722-214675               UUUCCAUACCAUAGAGAUUUU..CUAC......AG.............................AAAAUCUCUAUGGUAUGGAAA
#=GR NW_019384576.1/214722-214675       PP *********************..****......**.............................*********************
NW_006921800.1/5775323-5775272             UUUCCAAAGCUGGAAGAUUAU..AUAUau..auAU.............................AAAAUCUUCCAGCUUUGGAAA
#=GR NW_006921800.1/5775323-5775272     PP *********************..999999..9999.............................99*******************
NW_017547282.1/2530331-2530380             UUUUCAGAUGAGCCAGAAUUU..AUAGu....uAU.............................AAAUUCUGGCUCAUCUGAAAA
#=GR NW_017547282.1/2530331-2530380     PP *********************..*****....***.............................*********************
NW_004168040.1/2668-2617                   UUUUGAAAUGUCAAAUGUUGA..UUAAac..uuAA.............................UCAACAUUUGACAUUUCAAAA
#=GR NW_004168040.1/2668-2617           PP *********************..***988..8888.............................*********************
NW_017547282.1/2530380-2530331             UUUUCAGAUGAGCCAGAAUUU..AUAAc....uAU.............................AAAUUCUGGCUCAUCUGAAAA
#=GR NW_017547282.1/2530380-2530331     PP *********************..*****....***.............................*********************
NW_018029262.1/96813-96887                 UUUUCAGAUUUUCAAGGUUCA.gAUAG......--ucuacgccagcccuucaaguaaacaauc.UGAACCUUGAAAAUCUGAAAA
#=GR NW_018029262.1/96813-96887         PP *********************.99985........9999***********************9.*********************
NW_018034367.1/8706-8780                   UUUUCAGAUUUUCAAGGUUCA.gAUAG......--ucuacgccagcccuucaaguaaacaauc.UGAACCUUGAAAAUCUGAAAA
#=GR NW_018034367.1/8706-8780           PP *********************.99985........9999***********************9.*********************
NW_004168040.1/2617-2668                   UUUUGAAAUGUCAAAUGUUGA..UUAAgu..uuAA.............................UCAACAUUUGACAUUUCAAAA
#=GR NW_004168040.1/2617-2668           PP *********************..***999..9999.............................*********************
NC_024218.1/56989390-56989437              UUUCACCAGACAAAAGACUAU..AUAU......AU.............................AUAGUCUUUUGUCUGGUGAAA
#=GR NC_024218.1/56989390-56989437      PP *********************..****......**.............................*********************
NC_024218.1/56989437-56989390              UUUCACCAGACAAAAGACUAU..AUAU......AU.............................AUAGUCUUUUGUCUGGUGAAA
#=GR NC_024218.1/56989437-56989390      PP *********************..****......**.............................*********************
NW_007281290.1/14644686-14644733           UUUCACCAGACAAAAGACUAU..AUAU......AU.............................AUAGUCUUUUGUCUGGUGAAA
#=GR NW_007281290.1/14644686-14644733   PP *********************..****......**.............................*********************
NW_007281290.1/14644733-14644686           UUUCACCAGACAAAAGACUAU..AUAU......AU.............................AUAGUCUUUUGUCUGGUGAAA
#=GR NW_007281290.1/14644733-14644686   PP *********************..****......**.............................*********************
NC_024336.1/15544886-15544934              UUUCAUAAACAACUAGAUUGU..UUAU.....gAA.............................ACAAUCUAGUUGUUUAUGAAA
#=GR NC_024336.1/15544886-15544934      PP *********************..****.....999.............................*********************
NW_007614802.1/2847402-2847450             UUUCAUAAACAACUAGAUUGU..UUAU.....gAA.............................ACAAUCUAGUUGUUUAUGAAA
#=GR NW_007614802.1/2847402-2847450     PP *********************..****.....999.............................*********************
NC_036461.1/27413774-27413823              UUUAGAAAUGUUUCAGAUUGU..CUUGa....uAG.............................ACAAUCUGAAACAUUUCUAAA
#=GR NC_036461.1/27413774-27413823      PP *********************..**998....999.............................*********************
NW_019369745.1/27413774-27413823           UUUAGAAAUGUUUCAGAUUGU..CUUGa....uAG.............................ACAAUCUGAAACAUUUCUAAA
#=GR NW_019369745.1/27413774-27413823   PP *********************..**998....999.............................*********************
NW_005855045.1/13626287-13626338           UUUCGAAAUAAGCUAUUUUGA.aAUAGu...gaAU.............................UCAAAAUAGCUUAUUUCGAAA
#=GR NW_005855045.1/13626287-13626338   PP *********************.******...****.............................*********************
NW_006712453.1/199105-199056               UUCUGCAAGUUCCAAGACUAUacAUAU......AU.............................AUAGUCUUGGAACUUGCAGAA
#=GR NW_006712453.1/199105-199056       PP *********************888999......**.............................*********************
NW_007931123.1/62552485-62552533           UUUCCAAAUAGACUAUACUAA.uAUAA......AA.............................UUAGUAUAGUCUAUUUGGAAA
#=GR NW_007931123.1/62552485-62552533   PP *********************.*****......**.............................*********************
NW_019862807.1/1213391-1213437             UUUCUCGACUUUUUAUAUUUU.cAUAA......--.............................AAAAUAUAAAAAGUCGAGAAA
#=GR NW_019862807.1/1213391-1213437     PP *********************.*****.....................................*********************
NW_018408887.1/709331-709383               UUUUGACAUAUCCAAUGUUAU..AUUGgg.auaAU.............................AUAACAUUGGAUAUGUCAAAA
#=GR NW_018408887.1/709331-709383       PP *********************..****9*.*****.............................*********************
NW_006270091.1/177164-177214               UUUCUAAAAUACAAAGAUUCU.gAUUU.....gAU............................uAGAAUCUUUGUAUUUUAGAAA
#=GR NW_006270091.1/177164-177214       PP *********************.*8887.....777............................7*********************
NC_037351.1/7390267-7390220                UUUUAAUAUCUGGGAUAUUAA..UUUA......AU.............................UUAAUAUCCCAGAUAUUAAAA
#=GR NC_037351.1/7390267-7390220        PP *********************..****......**.............................*********************
NW_020190108.1/7390267-7390220             UUUUAAUAUCUGGGAUAUUAA..UUUA......AU.............................UUAAUAUCCCAGAUAUUAAAA
#=GR NW_020190108.1/7390267-7390220     PP *********************..****......**.............................*********************
NW_018408887.1/709383-709331               UUUUGACAUAUCCAAUGUUAU..AUUAuc.ccaAU.............................AUAACAUUGGAUAUGUCAAAA
#=GR NW_018408887.1/709383-709331       PP *********************..88888*.*****.............................*********************
NW_005857347.1/53777-53822                 UUUUGAAAUGUACAAGAGCCC..AUAG......--.............................GGGCUCUUGUACAUUUCAAAA
#=GR NW_005857347.1/53777-53822         PP *********************..****.....................................*********************
NC_037351.1/7390220-7390267                UUUUAAUAUCUGGGAUAUUAA..AUUA......AA.............................UUAAUAUCCCAGAUAUUAAAA
#=GR NC_037351.1/7390220-7390267        PP *********************..****......**.............................*********************
NW_020190108.1/7390220-7390267             UUUUAAUAUCUGGGAUAUUAA..AUUA......AA.............................UUAAUAUCCCAGAUAUUAAAA
#=GR NW_020190108.1/7390220-7390267     PP *********************..****......**.............................*********************
NW_018734406.1/388247771-388247818         UUUUUCAAUUAGAGAGUAUAU..AUAU......AU.............................AUAUACUCUCUAAUUGAAAAA
#=GR NW_018734406.1/388247771-388247818 PP *********************..****......**.............................*********************
NC_035109.1/388247771-388247818            UUUUUCAAUUAGAGAGUAUAU..AUAU......AU.............................AUAUACUCUCUAAUUGAAAAA
#=GR NC_035109.1/388247771-388247818    PP *********************..****......**.............................*********************
NW_011876287.1/108488-108541               UUUCGGAAUCCAAAAAAUUAU..AUAUacauacAU.............................AUAAUUUUUUGGAUUCCGAAA
#=GR NW_011876287.1/108488-108541       PP *********************..****99999999.............................*********************
NW_017127682.1/20574806-20574747           UCUCGAAAUUUUUAAGAUUCU..AUUA......--aacaaua..............gaauaau.AGAAUCUUAAAAAUUUCGAGA
#=GR NW_017127682.1/20574806-20574747   PP *********************..8876........3555566..............6666666.*********************
NW_019384975.1/406970-407017               UUUCAAAAUCAAAUUUGUUCU..UUUG......AU.............................AGAACAAAUUUGAUUUUGAAA
#=GR NW_019384975.1/406970-407017       PP *********************..****......**.............................*********************
NW_019384975.1/418663-418710               UUUCAAAAUCAAAUUUGUUCU..UUUG......AU.............................AGAACAAAUUUGAUUUUGAAA
#=GR NW_019384975.1/418663-418710       PP *********************..****......**.............................*********************
NW_019385577.1/207230-207280               UUUCUAGAUGAUGAGUAUUAG..AUUGa...caAU.............................CUAAUACUCAUCAUCUAGAAA
#=GR NW_019385577.1/207230-207280       PP *********************..*****...****.............................*********************
NW_019385577.1/207280-207230               UUUCUAGAUGAUGAGUAUUAG..AUUGu...caAU.............................CUAAUACUCAUCAUCUAGAAA
#=GR NW_019385577.1/207280-207230       PP *********************..*****...****.............................*********************
NW_003943645.1/9535070-9535019             UUUUUGAAUUAGCGAUGCUCA.gUUAAc...ugAU.............................UGAGCAUCGCUAAUUCAAAAA
#=GR NW_003943645.1/9535070-9535019     PP *********************.77777*...****.............................*********************
#=GC SS_cons                               :<<<<<<<<<<<<<<<<<<<<..____......__~~~~~~~~~~~~~~~~~~~~~~~~~~~~.>>>>>>>>>>>>>>>>>>>>:
#=GC RF                                    UUUCgAgAUuuacgAGgUUcu..AUAg......AU~~~~~~~~~~~~~~~~~~~~~~~~~~~~.agAAcCUcguaaAUcUcGAAA
//

#for motif_4508
# STOCKHOLM 1.0

NC_007932.1/170940-171045         GAAGUAUCUAUAAGUUAAUCAACAACUUUUAGAAUUUCAGUAUUUCAAACACUAACUAGAAUGGGUAACUUUAUACUAAAAGUUGUCAAUCUAAAUUAAGGAGAUA
#=GR NC_007932.1/170940-171045 PP **********************************************************************************************************
NC_024050.1/172235-172340         GAAUUAUUUAUGAGUUUUCUAACAACUUAUAGAAGUUCAGUAUUUCAAACACUAACUAGAAUUGGUAACUUCACAUUUAAAGUCAUCAAUCUAAAUUAAGGAGACA
#=GR NC_024050.1/172235-172340 PP **********************************************************************************************************
NC_029861.1/64483-64588           GAAGUAUCUAUAAGUUUUUUGAUAACUUUUAGAAUUUCAGUAUUUCAAACACUAACUAGAAUUGAUAACUUUCAAUUAAAAGCUGUCAAUCUAAAUUAAGGAGAUA
#=GR NC_029861.1/64483-64588   PP **********************************************************************************************************
NC_000925.1/170029-170133         GAAAUCUCUAUAAGUUGAUU-UCAGCUUUGAGAAUUUCGGUAUUUCAAACACUAACUAGAAUUGGUAACUUAAUAUAACAAGUUAUAAAUCUAAAUUAAGGAGAUA
#=GR NC_000925.1/170029-170133 PP *****************999.99***********************************************************************************
NC_021189.1/173079-173184         GAAGUAUGUAUAAGUUUUUUGAGAACUUGUAUAAUUUCAGUAUUUCAAACAUUAACUAGAAUUGUUAACUUUAAAUUAAAAGUCAUCAAUCUAAAUUAAGGAGAUA
#=GR NC_021189.1/173079-173184 PP **********************************************************************************************************
NC_035573.1/169168-169272         GAAAUGUAUAUAAGUUGGUU-UCAGCUUUUAGAAUUUCGGUAUUUCAAAUACUAACUAGAAUUGGUAACUUAAUAUAAUAAGUUAUGAAUCUAAAUUAAGGAGAUA
#=GR NC_035573.1/169168-169272 PP *****************999.99***********************************************************************************
NC_029576.1/172208-172313         GAAAUAUUUAUAAGCUGCGGUAUGACUUAUAUUAUUUCGGUAUUUAAAAUACUAACUAAGAUUAAUAACUAUUUAUAUAAAGUUAUUAAUUUAAAUUAAGGAGAUA
#=GR NC_029576.1/172208-172313 PP **********************************************************************************************************
#=GC SS_cons                      <<<<<-<<<<<<<<<<_______>>>>>>>>>>>>>>>------------------<<<<-<<<<<<<<<<________>>>>>>>>>>>>>>:::::::::::::
#=GC RF                           GaaguaucuauaaGuuuuuuaauaaCuuauagaacuuCaGUAUUUcAAAcACUAACuaGaAuugguaaCuuuauAUuaaaaGuuaccaauCuaAAUUAAGGAGAUA
//


#for motif_5095
# STOCKHOLM 1.0

NC_022259.1/14070-14156         AACUUAAGGAAACAACAUUUGCU-CUAUAAUAAAGUAGCAAAUGUUGUUUCCUUAAGUUG.....AACUAU.......AUUUAUAAAgaACAAGUAAAUA
#=GR NC_022259.1/14070-14156 PP ***********************.************************************.....******.......**********************
NC_022260.1/14067-14153         AACUUAAGGAACCAAGAUUUACU-CUAUAAUAAAGUAGCAAAUCUUGGUUCCUUAAGUUG.....AACUAU.......AUUUAUAAAgaACAAGUAAAUA
#=GR NC_022260.1/14067-14153 PP ***********************.************************************.....******.......**********************
NC_022262.1/14095-14181         AACUUAAGGAAUCAAGAUUUGCU-CUAUAAUAAAGUAGCAAAUCUUGAUUUCUUAAGUUG.....AACUAU.......AUUUAUAAAgaACAAGUAAAUA
#=GR NC_022262.1/14095-14181 PP ***********************.************************************.....******.......**********************
NC_022263.1/14068-14154         AACUUAAGAAACCAAAAUUUACU-CUAUAACAAGGGAGUAAAUUUUGGUUUCCUAAGUUG.....AACUAU.......AUUUAUAAAgaACAGGUAAGUA
#=GR NC_022263.1/14068-14154 PP ***********************.************************************.....******.......**********************
NC_022260.1/14124-14031         AACUUAAGGAACCAAGAUUUGCUACUUUAUUAUAG-AGUAAAUCUUGGUUCCUUAAGUU-uuccuAAAUA-uauuauaACUUGUCUU..-CAAUCGAGUA
#=GR NC_022260.1/14124-14031 PP ***********************************.***********************.8888855555.5789999*********...**********
NC_022259.1/14127-14034         AACUUAAGGAAACAACAUUUGCUACUUUAUUAUAG-AGCAAAUGUUGUUUCCUUAAGUU-uuccuAAAUA-uauuauaACUUGUCUU..-CAAUCGAGUA
#=GR NC_022259.1/14127-14034 PP ***********************************.***********************.8888855555.5789999*********...**********
NC_022262.1/14152-14059         AACUUAAGAAAUCAAGAUUUGCUACUUUAUUAUAG-AGCAAAUCUUGAUUCCUUAAGUU-uuccuAAAUA-uauuauaACUUGUCUU..-CAAUCGAGUA
#=GR NC_022262.1/14152-14059 PP ***********************************.***********************.8888855555.5789999*********...**********
NC_022263.1/14125-14032         AACUUAGGAAACCAAAAUUUACUCCCUUGUUAUAG-AGUAAAUUUUGGUUUCUUAAGUU-uuccuAAAUA-uauuauaACUUGUCUU..-CAAUCGAGUA
#=GR NC_022263.1/14125-14032 PP ***********************************.***********************.8888855555.5789999*********...**********
NC_022261.1/14101-14187         AACUUACGAAACCAAGAUUUACU-AAUUUAAUAUUAAGUAGAUCUUAGUUCGAUAAGUUG.....AACUAU.......AUUUAUAAAgaACAAGUAGAUA
#=GR NC_022261.1/14101-14187 PP ***********************.888888888999************************.....******.......**********************
NC_020014.1/16628-16714         AACUUACGAAACCAAGAUUUACU-AAUUUAAUAUUAAGUAGAUAUUGGUUCAAUAAGUUG.....AACUAU.......AUUUAUAAAgaACAAGUAAAUA
#=GR NC_020014.1/16628-16714 PP ***********************.888888888999************************.....******.......**********************
NC_022261.1/14158-14065         AACUUAUCGAACUAAGAUCUACUUAAUAUUAAAUU-AGUAAAUCUUGGUUUCGUAAGUU-uuccuAAAUA-uauuauaACUUGUCUU..-CAAUCAAGUA
#=GR NC_022261.1/14158-14065 PP ***********************887666666666.***********************.8888855555.5789999*********...**********
#=GC SS_cons                    <<<<<<<<<<<<<<<<<<<<<<<-<<_______>>->>>>>>>>>>>>>>>>>>>>>>>-.....------.......<<<<<____.._____>>>>>:
#=GC RF                         AACUuAaggAacCAAgAuuUgCUaccuUaauAaggaAGcAaaUcUUGguUccuUaAGUUg.....AAaUAU.......acUugUaua..ACAAgcaAguA
//


#for motif_11540
# STOCKHOLM 1.0

NC_031175.1/231799-231917         CGUUUUGUUAUCUCCUUAUUAAGUU.....................................AUAAGG....UAGGUAAUAG..UUAAUCGC............AUUACUAC..AUGG.UACAAAAC.GGGUAGGAA.GG.AGAG.UAAC........UUCUCCGAAA..A..CUUUCUUAAUA.A-............U.A.AGAGAG
#=GR NC_031175.1/231799-231917 PP *************************.....................................******....**********..********............********..****.********.*********.**.****.****........**********..*..***********.9.............*.*.******
NC_031175.1/220663-220545         CGUUUUGUUAUCUCCUUAUUGAAUU.....................................AUAAGG....UAGGCUAUAG..UUAAUCGC............AUUGCUAU..AUGA.UACAAAAC.GGAUAGGAA.GG.AGAG.UAAC........UUCUCCAAAA..A..CUCUUUUAGAA.A-............U.A.AAAGAG
#=GR NC_031175.1/220663-220545 PP *************************.....................................******....**********..********............********..****.********.*********.**.****.****........**********..*..**********9.9.............*.*.******
NC_031175.1/64797-64916           CGCUUUGCUAUUUACUUAUUAAAUU.....................................AUGAGG....UAAGCAAUAG..UUAAUCGC............AUUGCUAA..AUGA.UGCAAAGC.GGGCAGGAA.GG.AGAG.UAAC........UUCUCUGAAA..A..CUCUCUUAUUC.AU............U.A.AGAGAG
#=GR NC_031175.1/64797-64916   PP *************************.....................................******....**********..********............********..****.********.*********.**.****.****........**********..*..***********.**............*.*.******
NC_031175.1/149061-148939         CGUUUUGUUAUCUCCUUAUUAAUUU............................u.......uAUAAGG....UAGGUAAUAG..UUAAUCGC............AUUGCUAG..AUGG.UACAAAAC.GGGUAGGAA.GG.AGAG.UAAC........UUCUCCGAAA..A.aCUCUCUUAUUA.AA............U.A.AGAAAG
#=GR NC_031175.1/149061-148939 PP **********************999............................9.......9******....**********..********............********..****.********.*********.**.****.****........**********..9.9***********.**............*.*.******
NC_023133.1/139718-139600         CGAAAUGAUAUCUCCUUAUUAAGUU.....................................AUAAGG....UAGGCAAUAG..UUAAUCGC............ACUGCCAU..AUGA.UACAUUUC.GGGUAGGAA.GG.CGAG.UAAC........UUCGCCGAAA..A..GUUAGAUAGUA.A-............U.A.UCUAGC
#=GR NC_023133.1/139718-139600 PP *************************.....................................******....**********..********............********..****.********.*********.**.****.****........**********..*..***********.*.............*.*.******
NC_031175.1/200304-200422         CGUUUUGCUAUCUCCUUAUUAAGUU.....................................AUAAGG....UAGGUAAUAG..UUAAUCGU............AUUACUAC..AUGG.UACAAAAC.GGGUAGGAA.GG.AGAG.UAAC........UUCUCUGAAA..A..CUGUCUUAGUA.A-............U.G.AGAGAG
#=GR NC_031175.1/200304-200422 PP *************************.....................................******....**********..********............********..****.********.*********.**.****.****........**********..*..***********.*.............*.*.******
NC_031175.1/253088-252970         CGCUUUGUUAUCUCCUUGUUAAAAU.....................................AUAAGG....UGGGUAAUAG..UUAAUCGC............AUUGCUAC..AUGA.UACAAGGC.GGGUAGGAA.AG.AGAG.UAAC........UUCUCUGAAA..A..UUCUCUUGUUG.A-............U.A.AGAGAA
#=GR NC_031175.1/253088-252970 PP *************************.....................................******....**********..********............********..****.********.*********.**.****.****........**********..*..***********.9.............*.*.******
NC_031175.1/224436-224557         CGUUUUGUUAUCUCCUUAUUAAGUU.....................................AUAAGG....UAGGUAAUAG..UUAAUCGC............AUUACUAC..AUGG.UACAAAAC.GGGUAGGAA.GG.AGAG.UAAU........UUCUCUAAAAuaA..UUAUCAGUCUU.CG............A.C.UGAUAA
#=GR NC_031175.1/224436-224557 PP *************************.....................................******....**********..********............********..****.********.*********.**.****.****........********99745..***********.**............*.*.******
NC_023133.1/104566-104682         CGGAAUGCUAUCUCCUUAUCAAGUU.....................................AUAAGG....UAGGUACUAG..UUAAUCGC............AGCACCAU..AUGA.UGCAUUUC.GGGUAGGAA.GG.GGAG.UAAU........UUCCCCGAAA..A..GCUUUAGUUU-.--............A.C.UAAAGC
#=GR NC_023133.1/104566-104682 PP *************************.....................................******....**********..********............********..****.********.*********.**.****.****........**********..*..*******998................9.*.******
NC_031175.1/142329-142210         CGCUUUGGUAUCUACUUAUUGAGUU.....................................AUAAAG....UAAGCAAUAG..UUAAUCGC............AUCGCUAC..AUGA.UGCAAAGC.GGGUAGGAA.AG.CGAG.UAAA........UUCGCUAAAA..A.aCUUUCUUAGUU.A-............U.A.AGAAAG
#=GR NC_031175.1/142329-142210 PP *************************.....................................******....**********..********............********..****.********.*********.**.****.****........**********..9.9***********.*.............*.*.******
NC_031175.1/188058-188177         CGUUUUGUUAUCUCCUUAUUAAUUU............................u.......uAUAAGG....UAGGUAAUAG..UUAAUCGC............AUUGCUAG..AUGG.UACAAAAC.GGGUAGGAA.GG.AGAG.CAAC........UUCUCCGAAA..A..GCUAAAUUUU-.-A............A.A.UUUAGC
#=GR NC_031175.1/188058-188177 PP **********************999............................9.......9******....**********..********............********..****.********.*********.**.****.****........**********..*..*******988...6............9.*.******
NC_023133.1/178990-178872         CGAAUUAAUAUCUCCUUACUAAGUU.....................................AUAAGG....UGGGUCUUAG..UUAAUCGC............AAGGCUAA..AUGA.AUUAAUUC.GGGUAGGAA.GG.CGAG.UAAC........UUCGCCAAAA..A..GCUAGAUAGUA.A-............U.A.UUUAGC
#=GR NC_023133.1/178990-178872 PP *************************.....................................******....**********..********............********..****.********.*********.**.****.****........**********..*..***********.*.............*.*.******
NC_031175.1/176859-176978         CGCUUUGUUAUUUCCUUAUUAAGUU.....................................AUAAGG....UGGGUAAUAA..UUAAUUGG............AUUGCUAC..AUGA.UGCAAAGC.GGGUAGGAA.AG.CGAG.UAAC........UUUGCUAAAA..A.aCUUUUUUAGUC.A-............U.A.AAAGAG
#=GR NC_031175.1/176859-176978 PP *************************.....................................******....**********..********............********..****.********.*********.**.****.****........**********..9.9***********.*.............*.*.******
NC_031175.1/835-953               CGUUUUGUUAUCUCCUUAUUAAAUU.....................................AUGAGG....UAGGCAAUAU..UUAGUCAC............AUUGCUAU..ACGA.UAUAAAGC.GGAUAGGA-.AG.AGAG.UAAC........UUCUCUGAAA..A..CUCUCUUAUUA.AU............U.A.AGAGAG
#=GR NC_031175.1/835-953       PP *************************.....................................******....**********..********............********..****.********.*******9..8*.****.****........**********..*..***********.**............*.*.******
NC_031175.1/157611-157734         CGUUUUGUUAUCUCCUUAUUAAUUU............................u.......uAUAAGG....UAGGUAAUAG..UUAAUCGC............AUUGCUAG..AUGG.UACAAAAC.GGGUAGGAA.GG.AGAG.UAAU........UUCUCCAAAAuaA..UUAUCAGUCUU.CG............A.U.UGAUAA
#=GR NC_031175.1/157611-157734 PP **********************999............................9.......9******....**********..********............********..****.********.*********.**.****.****........********99745..***********.**............*.*.******
NC_031175.1/205177-205060         AGCUUUGCUAUCUUCUUAUUAAGUU.....................................AUAAGG....UAAGCAAUAG..UUAAUCGA............AUUGCUAC..AGGA.UGCAAAGC.GGAUAGGAA.AG.AGAG.UAAC........UUCUCUGAAA..A..GCUAAAUUUU-.-C............A.A.UUUAGC
#=GR NC_031175.1/205177-205060 PP *************************.....................................******....**********..********............********..****.********.*********.**.****.****........**********..*..*******988...6............9.*.******
NC_031175.1/15541-15660           CGCUUUGCUAUUUCCUUAUUAAGUU.....................................AUAAAG....UAAGCUAUGU..UGAAUCAC............AUUGCUAC..AAGA.UUCAAAGC.GGAUAGGAA.AG.AGAG.UAAC........UUCUCUGAAA..A..CUCUCUUAAAU.AA............U.A.AGAGAG
#=GR NC_031175.1/15541-15660   PP *************************.....................................******....**********..********............********..****.********.*********.**.****.****........**********..*..***********.**............*.*.******
NC_031175.1/135818-135698         CGUUUUAUUAUUUCCUUAUGAAACU.....................................AUAAGG....UAAGCAAUAG..UAAAUCGC............AUUGCUCA..AUAA.UAUAAAAC.GGGUAGGAA.GG.AGAA.UAAC........UUCUCCAAAA..A.aUUAUCAGUCAA.AA............A.A.UGAUAA
#=GR NC_031175.1/135818-135698 PP *************************.....................................******....**********..********............********..****.********.*********.**.****.****........**********..9.9***********.**............*.*.******
NC_031175.1/6497-6615             CGUUUUGCUAUCUACUUAUUAAUUU.....................................AUAAGG....UAGGCAAUAG..CUAAUUGG............AUUGCUGC..AAGA.UGCAAAAC.GGGUAGGAA.UG.AGAG.UAAC........UCCUCCGAAA..A..CUUUCUUAGUU.A-............U.A.AGAAAG
#=GR NC_031175.1/6497-6615     PP *************************.....................................******....**********..********............********..****.********.*********.**.****.****........**********..*..***********.*.............*.*.******
NC_031175.1/61820-61940           CGCUUUGUUUUCUCCUUGUUAAAAU.....................................AUAAAA....UAGGCAAUAG..UUAAUCAC............AUUGCUAG..ACAA.UACAAAGC.GGGUAGGAA.GG.AGAA.UAAC........UUCUCCAAAA..A.gUUAUCAGUCGA.AA............A.C.UGAUAA
#=GR NC_031175.1/61820-61940   PP *************************.....................................******....**********..********............********..****.********.*********.**.****.****........**********..*.************.**............*.*.******
NC_031175.1/50205-50087           CGUUUUGCUAUCUCCUCAUUAAAUU.....................................ACAAGC....CAGGCAAUAG..UUAAUCGG............AUUGCCAU..ACGA.UGCAAAGC.GGGUAGGCA.GG.AGAA.UAAC........UUCUCCGAAA..A..CUCUCUUCGUA.A-............U.A.AGAGAG
#=GR NC_031175.1/50205-50087   PP *************************.....................................******....**********..********............********..****.********.*********.**.****.****........**********..*..***********.9.............*.*.******
NC_031175.1/202680-202797         CGCUUUGCUAUCUCCUUACUAAAC-.....................................AUAAGG....UAGGUAAUGG..UUAAUCGC............AUUACUAC..AUGG.UACAAAGC.GGGUAGGAA.GG.AGAG.UAAU........UUCUCCGAAA..A..CUCUCUUAGCU.A-............U.A.AGAUAA
#=GR NC_031175.1/202680-202797 PP ********************9999......................................9*****....**********..********............********..****.********.*********.**.****.****........**********..*..***********.*.............*.*.******
NC_031175.1/137213-137095         CGCUUUGUUUUCUCCUUAUUAAAU-.....................................AUAAGC....UAGGCAAUAG..UUAAUCUG............AUUGCUAC..AUGA.UGGAAAGC.GGGUAGGAA.AG.CGAG.UAAA........UUCGCUAAAA..A.aCUUUCUCAGUU.A-............U.A.AGAAAG
#=GR NC_031175.1/137213-137095 PP ********************9999......................................******....**********..********............********..****.********.*********.**.****.****........**********..9.9***********.*.............*.*.******
NC_031175.1/139820-139703         UGCUUUGUUUUCUCCUUAUUAAAU-.....................................AUAAGC....UAGGCAAUAG..UUAAUCUG............AUUGCUAC..AUGA.UGCAAAGC.GGGUAGGAG.GG.AGAG.UAAU........UUCUCCAAAA..A..CUCUCUUAGUA.A-............U.A.GGAGAA
#=GR NC_031175.1/139820-139703 PP ********************9999......................................******....**********..********............********..****.********.*********.**.****.****........**********..*..***********.*.............*.*.******
NC_031175.1/159362-159483         CGCUUUGUUAUCUCCUUGUUAAAAU.....................................AUAAGG....UGGGUAAUAG..UUAAUCGC............AUUGCUAC..AUUA.UGCAAAGC.GGGUAGGAA.AG.AGAG.UAAC........UUCUUCAAAA..AaaUUAUUAGUUUU.CG............A.C.UGAUAA
#=GR NC_031175.1/159362-159483 PP *************************.....................................******....**********..********............********..****.********.*********.**.****.****........****999998..778***********.**............*.*.******
NC_031175.1/247228-247108         CGCUUUAGUAUCUCCUUGUUAAAAU.....................................AUAAAG....UAAGCAAUAU..UUAGUCGU............AUUGCUAU..ACGA.UACAAAGC.GGGUAGGAA.GG.AGAG.UAAC........UUCUCCGAAG..A.cUUAUCAGUCUC.UG............A.U.UGAUAA
#=GR NC_031175.1/247228-247108 PP *************************.....................................******....**********..********............********..****.********.*********.**.****.****........**********..*.************.**............*.*.******
NC_034787.1/51201-51316           CACGUUAA-UUUUCCUUAA--AAUA.....................................UUAAGG....UAAGUUUCAG..UUAAUCGC............GAAACUAU..AUGG.GUUAACGU.GGGUAGGAA.GG.UGAG.UAAC........UUCACCGAAA..A..GUUUUAGAUAA.A-............A.C.UAAAUC
#=GR NC_034787.1/51201-51316   PP ********.9999******..8999.....................................******....**********..********............********..****.********.*********.**.****.****........**********..*..*******9999.9.............9.*.******
NC_031175.1/164623-164740         CGCUUUGUUUUCCCUUUAUUAAAU-.....................................AUAAGG....CAGUCAACAG..UUAAUCUG............AUUGCUAC..AUGA.UGCAAAGC.GGGUAGGAA.GG.AGAG.UAAU........UUCUCCGAAA..A..UUCUCUUGGUA.A-............U.G.AGAGAU
#=GR NC_031175.1/164623-164740 PP ********************9999......................................******....**********..********............********..****.********.*********.**.****.****........**********..*..***********.9.............*.*.******
NC_031175.1/145652-145533         CGCUUUGUUAUGUCCUCAUUCAGCU.....................................AUAAAC....UAGGCAAUAG..CUAAUCGG............AUUGGUAC..AUGA.UACAAAGC.GAGUAGGAA.GG.AGGG.UAAC........UUCUCUGAAA..A..CUCUCUUAUUC.AU............U.A.AGAGAG
#=GR NC_031175.1/145652-145533 PP *************************.....................................******....**********..********............********..****.********.*********.**.****.****........**********..*..***********.**............*.*.******
NC_031175.1/45697-45824           CGCUUUGAUGUAUUCUUAUUAGGCU............................uauuaagcuAUAAGA....UAAGUGAUAG..UUAAUUCC............AUUGCUAU..GUGA.UACAAAGC.GGAUAGGAA.AG.AGAG.UAAC........UUCUCUGAAA..A..CUCUCCUAGUA.A-............U.G.AGAGAG
#=GR NC_031175.1/45697-45824   PP ********************97765............................666666666******....**********..********............********..****.********.*********.**.****.****........**********..*..***********.9.............*.*.******
NC_031175.1/193001-193118         CGUUUUGUUAUCUCCUUGUUAAAAU.....................................AUAAAA....UAGGCAAUAG..UUAAUCGC............AUUGCUAC..AGGA.UGCAAAGC.UGAUAGGAA.AG.AGAG.UAAC........UUCUCUGAAA..A..GCUAAAUUUU-.-C............A.A.UUUAGC
#=GR NC_031175.1/193001-193118 PP *************************.....................................******....**********..********............********..****.********.*********.**.****.****........**********..*..*******988...6............9.*.******
NC_034787.1/475577-475462         CGUGUUAA-GUUUCCUUAA--AAUA.....................................UUAAGG....UGUGUUUCAG..UUAAUCGC............GAAACUAU..AUGG.GUUAACGC.GGGUAGGAA.GG.UGAG.UAAC........UUCACCGAAA..A..GUUUUAGGUAA.A-............A.C.UAAAAU
#=GR NC_034787.1/475577-475462 PP ********.**********..8999.....................................******....**********..********............********..****.********.*********.**.****.****........**********..*..*******9999.9.............9.*.******
NC_031175.1/249325-249205         CGCUUUGCUAUCUUGUUAUUAAGUU.....................................CUAAAA....UAAGCGAUAU..UAGAUCAC............AUUGCUAC..AGGA.UGCAAAGC.AAGUAGGAA.GG.AGAU.UAAU........UUCUCCGAAA..A.gUCAUCAGUUUU.UA............A.C.UGAUGA
#=GR NC_031175.1/249325-249205 PP *************************.....................................******....**********..********............********..****.********.*********.**.****.****........**********..*.************.**............*.*.******
NC_023133.1/180465-180350         CGAAAUGUUAUCUCCUUAAUGAACU.....................................AUGAGG....UAGGUAUUAG..UUAAUCGC............AAUACCAU..AAGG.UGCAUUUU.GGGUAGGAA.GG.GGAG.UAAC........UUCCCUGAAA..A..ACCUGAC--UU.U-............-.G.UCGGGG
#=GR NC_023133.1/180465-180350 PP *************************.....................................******....**********..********............********..****.********.*********.**.****.****........**********..*..9999998..44.4...............8.99999*
NC_034787.1/223286-223403         CGCGUUAA-GUUUCCUUAA--AAUA.....................................UUAAGG....UGGGUUUCAG..UUAAUCGC............GAAACUAU..AUUG.GUUAACGU.GGGUAGGAA.GG.UGAG.UAAC........UUCACCGAAA..A.gUUUUAAGUGAA.AA............A.U.UUAAAC
#=GR NC_034787.1/223286-223403 PP ********.**********..8999.....................................******....**********..********............********..****.********.*********.**.****.****........**********..*.88899*******.**............*.*.******
NC_034787.1/396483-396366         CGCGUUAA-GUUUCCUUAA--AAUA.....................................UUAAGG....UGGGUUUCAG..UUAAUCGC............GAAACUAU..AUUG.GUUAACGU.GGGUAGGAA.GG.UGAG.UAAC........UUCACCGAAA..A.gUUUUAAGUGAA.AA............A.U.UUAAAC
#=GR NC_034787.1/396483-396366 PP ********.**********..8999.....................................******....**********..********............********..****.********.*********.**.****.****........**********..*.88899*******.**............*.*.******
NC_031175.1/244975-244855         CGCUUAGCUAUAUCCUUAUUAAGCU.....................................AUAAAG....UUAGCAAGAA..UUAAUAGA............AUUACUAG..AUGA.UGCAAAGC.GGGUAGGAA.GG.AGAA.UAAC........UUCUCCAAAA..A.gUUAUCGGUCUU.UG............A.U.UGAUAA
#=GR NC_031175.1/244975-244855 PP *************************.....................................******....**********..********............********..****.********.*********.**.****.****........**********..*.************.**............*.*.******
NC_023133.1/153264-153381         CGAAAUGCCAUCUGCUUAUUAGACU.....................................AUAAAG....UUGGUAUUAG..UUAACAAC............AAUACUAC..AAGA.UGCAUUUU.GGGUAGGAA.GG.GGAG.UAAC........UUCUCCGAAA..A..GCUUCAG-CUA.CA............-.C.UGAAGC
#=GR NC_023133.1/153264-153381 PP *************************.....................................******....**********..********............********..****.********.*********.**.****.****........**********..*..*****99.566.65..............9.9*****
NC_031175.1/156527-156644         CGUUUUGCUAUCUCCUUAUUAAGUU.....................................AUAAGG....UAGGCAAUAG..UAAAUCAC............AUUGCUAU..AUGA.UCUAAAAC.GGGCAGGUA.GG.GGAU.CAAC........UUCCCUA--A..A..ACUCUCUUUUA.CU............A.A.GAGAAU
#=GR NC_031175.1/156527-156644 PP *************************.....................................******....**********..********............********..****.********.*********.**.****.****........******5..5..6..889********.9*............*.*.******
NC_031175.1/162406-162526         UGCUUGGGUAUCUCCUUCUUAAUUU.....................................AUAAGG....UAGGCAACAG..UUAAUUUU............AUUGCUAU..ACGA.UACACAGC.GGGUAGGAA.GG.AGAG.UAAC........UUCUCCAAAA..A.gUUAUCGGUCUC.UG............A.U.UGAUAA
#=GR NC_031175.1/162406-162526 PP *************************.....................................******....**********..********............********..****.********.*********.**.****.****........**********..*.************.**............*.*.******
NC_034787.1/7352-7469             CGCGUUAA-GUUUCCUUAA--AAUA.....................................UUAAGG....UAUGUUUCAG..UUAAUCGC............GAAACUAU..AUGG.GUUAACGU.GGGUAGAAA.GG.UGAG.UAAC........UUCACCGAAA..A..GUUUUAAAUGAaAA............U.U.UAACAC
#=GR NC_034787.1/7352-7469     PP ********.**********..8999.....................................******....**********..********............********..****.********.*********.**.****.****........**********..*..*******9876477............*.*.******
NC_031175.1/74494-74612           CGUUUUGUUAUCAUUUCAUUAAUAU.....................................AUAACU....UAGGCAAUAG..UUCAUUAU............AUCCUGAA..AUGA.UACAAAAC.GAAUAGGAA.GG.AGAG.UAAC........UUCUCUGAAA..A..CUCUCCUAGUA.A-............U.G.AGAGAG
#=GR NC_031175.1/74494-74612   PP *************************.....................................******....**********..********............********..****.********.*********.**.****.****........**********..*..***********.9.............*.*.******
NC_023133.1/193938-194056         CGAAGUGUCAUCUUAUUACUAAAUC....................................aUUAAAG....UAAGUAUUAG..UUAAUCAU............AAUAUCAU..AAGA.UGCAUUUU.GGGUAGGAA.GG.GGAG.UAAC........UUCCCUGAAG..A..GCUUCAGUUA-.-A............A.C.UGAGGC
#=GR NC_023133.1/193938-194056 PP *********************9998....................................6888888....**********..********............********..****.********.*********.**.****.****........**********..*..*******866...5............9.*.******
NC_034787.1/123043-123159         CCUGUUAA-GUCUCCUUAA---AUA.....................................UUAAGG....UGUGUUUCAG..UUAAUCGU............GAAACUAU..AUGG.GUUAACGU.GGUUAGGAA.GG.UGAG.UAAC........UUCACCGAAA..A..GUUUUAAAUGAaAA............U.U.GAAGAC
#=GR NC_034787.1/123043-123159 PP ********.*********9...677.....................................9*****....**********..********............********..****.********.*********.**.****.****........**********..*..******99876467............9.9.9*****
NC_062388.1/213182-213064         CAUAUAAAGUUCUCCUUAUUAAAAA.....................................AUAAGA....UAGGUCUUAG..UUAAUCGU............AAAACUAU..AUAA.AUGCAUAU.GGCUAGGAA.GG.UGAG.GAAC........UUCACCGAAA..A..GCUUUUGAUUU.A-............U.U.AAGUGC
#=GR NC_062388.1/213182-213064 PP *************************.....................................******....**********..********............********..****.********.*********.**.****.****........**********..*..***********.*.............*.*.******
NC_023133.1/164492-164379         CGAAAUGUCAUCUGCUUAUUAGACU.....................................AUAAAG....UUGGUAUUAG..UUAA----............CAUACUAC..AAGA.UGCAUUUC.GGGUAGGAA.GG.GGAG.UAAC........UUCCCUGAAG..A..GCUUCAG-CUA.CA............-.C.UGAGGC
#=GR NC_023133.1/164492-164379 PP *************************.....................................******....**********..****................********..****.********.*********.**.****.****........**********..*..******9.566.65..............9.******
NC_031175.1/152403-152523         CGGAUUAAUUUCCCUUUAUCAAAGA.....................................AUUAAG...aUAUUUUUUAG..UUAAUUGU............GCGAUUAU..AUGG.AUUAAUUC.GGGUAUGAA.UG.CGAG.UAAC........UUCGCUAAAA..A.aCUUUCUUAAUU.A-............U.A.AGAAAG
#=GR NC_031175.1/152403-152523 PP *************************.....................................999999...89999999***..********............********..****.********.*********.**.****.****........********99..9.9***********.*.............*.*.******
NC_031175.1/91482-91603           CGCUUUGUUGUCUACUAAUUAAAUU.....................................AUUAUA....GAAAUAAUAA..UUGAUAGA............AUUACUAU..AUAA.UACGAAGC.GGGUAGGAA.GG.AGAG.UAAA........UUCUCCAAAA..AagUUAUUAGUCUU.UG............A.U.UGAUAA
#=GR NC_031175.1/91482-91603   PP *************************.....................................******....**********..********............********..****.********.*********.**.****.****........*********9..99************.**............*.*.******
NC_031175.1/63427-63544           CGAGUUAA-GUCUUAUAAAUAAAAA.....................................AUAAAG....UAAGCUUAUA..UUAAUUUU............AGAAAUGU..UUAG.CUUAACUC.GGGUAGGAA.AG.AGAG.UAAC........UUCUCCGAAA..A..CUCUCUUGAUA.A-............U.A.AGAGAG
#=GR NC_031175.1/63427-63544   PP ********.****************.....................................******....**********..********............********..****.********.*********.**.****.****........**********..*..***********.9.............*.*.******
NC_062394.1/449447-449326         CAAAUUGGCUUCUCCUUAAU-----.....................................AGAAGG....UAGGUCAGAA..GUAAUUGU............CUGGCUAU..AUGA.ACUAAUUU.GGGUAGGAA.GG.GGAG.UAAC........UUCCCCGAAA..A..ACCAGCAA---.--gaauauuaauuuU.C.ACUGGU
#=GR NC_062394.1/449447-449326 PP ****************9442..........................................569***....**********..********............********..****.********.*********.**.****.****........**********..*..***99765......5678999999995.6.799***
#=GC SS_cons                      ((((((((-(((,<<<<<<______~~~~~~~~~~~~~~~~~~~~~~~~~~~~.........>>>>>>....,,<<<<<<__..________~~~~~~~~~~~~>>>>>>,,..,,)).)))))))).),,,,,,,,.<<.<<<<.____~~~~~~~~>>>>>>,,,,..,..<<<<<<<<___.__~~~~~~~~~~~~>.>.>>>>>>
#=GC RF                           CGcuuUgcUaucUccUuAuUAAauU~~~~~~~~~~~~~~~~~~~~~~~~~~~~.........aUaAgg....UAggcaauAG..UUAAUcgc~~~~~~~~~~~~auugccAu..AUga.ugcAaagC.GGGUAGGAA.gG.cGAg.UAAC~~~~~~~~uUCgCcgAAA..A..cucucuuuuUa.aa~~~~~~~~~~~~a.a.agagag
//


#for Motif_15322
# STOCKHOLM 1.0

NC_039971.1/12788-12877           CAGUAUUAUGAGGGAGAGCGU.AGU.GAA.UC.GAGAGGUUCUCGCUACGUUCGG.A..GGA.AGCUU..GACUAGAU.UUAG.............................UUCUA.GGCA.UUC.GAUCCU
#=GR NC_039971.1/12788-12877   PP *********************.***.***.**.**********************.*..***.*****..********.****.............................*****.****.***.******
NC_035156.1/49737-49826           CAGGUAAUUGGGUGAGAGCGU.UGU.GAA.UC.GAAAGAUUCUUGCAACGUUCGG.A..UGG.AGCUA..AAUCAAUA.AUAU.............................UAUUG.GUUU.UUC.GACCAU
#=GR NC_035156.1/49737-49826   PP *********************.***.***.**.**********************.*..***.*****..********.****.............................*****.****.***.******
NC_027269.1/46534-46624           AAGCUCUUUGGGGGAGAGCGUuGGU.GAA.UC.GAAAGGUUCUCGCCACGUUCGG.A..CGG.AGCUU..GAUUUAUU.UUAG.............................AGUAG.AUCA.UUC.GACCGU
#=GR NC_027269.1/46534-46624   PP *************************.***.**.**********************.*..***.*****..********.****.............................*****.****.***.******
NC_071861.1/65524-65436           GAGUUUUAUGGGGGAGAGCGU.AGU.GAA.UC.GAGAGGUUCUCGCUACGUUCGG.A..GGA.AGCUU..GCAUAGGA.UA-G.............................ACUUA.UGCA.UUC.GAUCCU
#=GR NC_071861.1/65524-65436   PP *********************.***.***.**.**********************.*..***.*****..*******9.65.8.............................9****.****.***.******
NC_035156.1/5657-5747             CAGUUUUAUGGGGGAGAGCGU.AGU.AAA.UU.GAAAGGUUUUCGCUACGUUCGG.A..GGA.AGCUU..GUAUAGAUuAGUU.............................AGCUA.UACU.UUC.GAUCCU
#=GR NC_035156.1/5657-5747     PP *********************.***.***.**.**********************.*..***.*****..********99999.............................*****.****.***.******
NC_038187.1/59417-59327           AAACUUCUUGGGGGAGAGCGUuGGU.GAA.UC.GAGAGGUUCUCGCCACGUUCGG.A..CGG.AGCUA..AGUUCUUC.GUAA.............................GAUGG.AUUU.UUC.GACCGU
#=GR NC_038187.1/59417-59327   PP *************************.***.**.**********************.*..***.*****..********.****.............................*****.****.***.******
NC_038187.1/5165-5254             ACAAUUUUAGUAUUAAAGCGU.AAU.GAA.UU.UAUAGGUUCUCAUUACGUUUGC.A..GGA.AGCUU..GUUUAAAG.GUAU.............................UUUUU.AACA.UUC.AAUCCU
#=GR NC_038187.1/5165-5254     PP *********************.***.***.**.**********************.*..***.*****..********.****.............................*****.****.***.******
NC_027286.1/7528-7619             UAGUUUUAAGAGAAAAAGCGU.AAU.GAA.UU.AAGAAGUUCUCGUUACGUUUGA.A..GGA.AGCUU..GUAUAAAUaAUCU............................uGUGUU.UACA.UUC.UAUCCU
#=GR NC_027286.1/7528-7619     PP *********************.***.***.**.**********************.*..***.*****..********97766............................6*****.****.***.******
NC_071863.1/14597-14687           AAUUGUAGUGGGGGAGAGCGU.AGU.GAA.UC.GUAAGGUUCUCGCUACGUUCGG.A..GGA.AGCUU..GUAUUCGUuAACG.............................ACGGG.UGCA.UUC.GAUCCU
#=GR NC_071863.1/14597-14687   PP *********************.***.***.**.**********************.*..***.*****..********99999.............................*****.****.***.******
NC_027286.1/87765-87675           AUAUUUAAGAAUGGAAAGCGU.AAU.GAA.AU.AAAAAUUUCUAGUUACGUUUGG.G..GGA.AGCUU..UUGUAAGAaAAUG.............................UUCUU.CGAA.UUC.UAUCCU
#=GR NC_027286.1/87765-87675   PP *********************.***.***.**.**********************.*..***.*****..***8877769999.............................*****.****.***.******
NC_071865.1/163612-163523         GUUUCUUGAAGGUGAGAGCGU.AGU.GAA.UC.GAGAGGUUCUCGCUACGUUCGG.A..GGA.CGCUU..GGACGGUU.UUAG.............................UAUCG.UGCU.UUC.GAUCCU
#=GR NC_071865.1/163612-163523 PP *********************.***.***.**.**********************.*..***.*****..********.****.............................*****.****.***.******
NC_039969.1/43048-42959           ACAGAUGAAGGGAGAAAGCGUaGGU.GAA.UU.GAAAGAUUCUCGCCACGUUUAG.U..UGG.AGCUU..AGCUGAAA.CA-A.............................AUUUG.GCUA.UUC.GACCAA
#=GR NC_039969.1/43048-42959   PP *************************.***.**.**********************.*..***.*****..*******9.76.5.............................8999*.****.***.******
NC_027286.1/3471-3562             AGAGUUUUUUGGUGAGAGCGUuAAU.AAA.UU.GAAAAGUUUUAGUUACGUUCGG.A..GGA.GGAUU..AGUCAGGAaUUUA.............................AUUUU.GCUA.UUC.UAUCCU
#=GR NC_027286.1/3471-3562     PP *************************.***.**.**********************.*..***.*****..**99999989999.............................*****.****.***.******
NC_071865.1/52661-52751           AAGUUUAAUAGUAGAGAGCGUaAAU.GAA.UU.GAAAAAUUCUUGUUACGUUUGG.A..GGA.AGUUC..UUUUGAAG.GUUA.............................CUUCA.UUAU.UUC.AAUCCU
#=GR NC_071865.1/52661-52751   PP *************************.***.**.**********************.*..***.*****..********.****.............................*****.****.***.******
NC_071865.1/71901-71812           ACAAAUCAUUUGAUAGAGCGUaGGU.GAA.UU.GAAAGGUUCUAGCCACGUUCGG.G..UGG.AGCUU..AUACUCAA.AAUU.............................UUGAG.-AUA.UUC.AACCAC
#=GR NC_071865.1/71901-71812   PP *************************.***.**.**********************.*..***.*****..**999999.9999.............................99998..***.***.******
NC_071859.1/131909-131819         UAUUUUAAUUUGGAAGAGCGU.AGU.GAA.UU.GAAAGGUUCUUGCUAUGUUCGA.A..GGG.AGAUU..AAACAGAUuAGAA.............................AGCUA.UUCA.UUC.UACCCU
#=GR NC_071859.1/131909-131819 PP *********************.***.***.**.**********************.*..***.*****..********99999.............................*****.****.***.******
NC_071859.1/87007-86918           AAGAUAGUUGAGUGAGAGCGU.AAU.GAA.UU.UGAAGAUUCUUUUUACGUUCGG.U..UGG.AGCUA..AAUUGAUAuUUAU.............................UAUUG.G-UU.UUC.UACCAA
#=GR NC_071859.1/87007-86918   PP *********************.***.***.**.**********************.*..***.*****..********56666.............................66666.5.56.79*.******
NC_071863.1/8760-8851             AAGAUUAUAGAGAAAGAGCGUaGGU.GAA.UCaGAAGGAUUCUCGCCACGUUCGG.A..CGG.AGUUA..AGCUUUUA.CUUG.............................AUAAA.GGAU.UUC.GACCGU
#=GR NC_071863.1/8760-8851     PP *************************.***.**4455589****************.*..***.*****..********.6666.............................66666.6667.89*.******
NC_071865.1/93952-93863           AAAGUUGAUAUAAAAGAGCGUgAGU.GAA.AU.GAAAGUUUCUUACCACGUUCGU.A..GGA.GG-UU..UUGUAAAG.AUGA.............................CUUUA.UAAU.UUC.AAUCCU
#=GR NC_071865.1/93952-93863   PP *********************77**.***.**.**********************.*..***.77.67..99999999.9999.............................99999.9999.***.******
NC_071865.1/30154-30244           AUAUUUUGGAGUUAAGAGCUU.GAU.AAA.UU.GAAAAGUUUUUAUUAAGUUCGU.A..GGA.GGCUU..AGGCGAAUaUUUG.............................AGACG.CUUA.UUC.UAUUCU
#=GR NC_071865.1/30154-30244   PP *********************.***.***.**.**********************.*..***.*****..********99999.............................*****.****.***.******
NC_039968.1/27948-28036           AUUUUGAUAUAUCAAGAGCGU.AGU.GAA.UU.UUAUUAUUCUUGCUUCGUUCGG.A..GGA.AGAUA..AAGUAAAA.GU-U.............................UUUUA.CUUU.UUC.AAUCCU
#=GR NC_039968.1/27948-28036   PP *********************.***.***.*9.99999*****************.*..***.*****..********.87.6.............................9****.****.***.******
NC_071859.1/112839-112749         AAAUUUUAAAAUAACAAGCAU.CGU.GAA.UA.UAAAUAUUCUUUCUAUGUUUUG.A..GGA.AGCUU..UUAUGGAAuAUUG.............................UUUCA.UAAU.UUC.UAUCCU
#=GR NC_071859.1/112839-112749 PP *********************.***.***.**.**********************.*..***.*****..********99999.............................*****.****.***.******
NC_039156.1/44745-44834           AAAGUUUUAAAAUAAAAGCUA.AGU.GAA.UU.UAAAGAUUCUUGCUUUGUUUGA.A..GGA.UGUUU..AUUGCGAU.UAUU.............................UUCGU.CUUU.UUC.UAUCCU
#=GR NC_039156.1/44745-44834   PP *********************.***.***.**.**********************.*..***.*****..********.****.............................*****.****.***.******
NC_071859.1/45445-45534           UUUGCUGUUUAGAGAAAACGU.AGU.GAA.UU.AAAAGGUUCUUGCUAUGUUUGA.A..GGA.AGGUU..ACUCAAAA.GAAU.............................UCUUU.UUUA.ACC.AAUCCU
#=GR NC_071859.1/45445-45534   PP *********************.***.***.**.**********************.*..***.*****..**999999.8888.............................99999.99**.***.******
#=GC SS_cons                      :::::::::::::::<<<<<<.<<<.<<<.<<.____>>>>>-->>>>>>>>>--.-..<<<.----<..<<<<<<<<.____~~~~~~~~~~~~~~~~~~~~~~~~~~~~.>>>>>.>>>>.---.-->>>:
#=GC RF                           aaauUUuuugggugAgAGCGU.gGu.GAA.uc.gAaAggUUCUcgCcACGUUcGg.a..gGg.AGCUu..ggccagac.auau~~~~~~~~~~~~~~~~~~~~~~~~~~~~.gucug.gcca.UUC.gAcCcU
//
```

**Part III Original gel pictures**

The samples were run on the same gel. The gel image was subsequently inverted using Adobe Photoshop, and corresponding lanes were cropped for figure preparation.

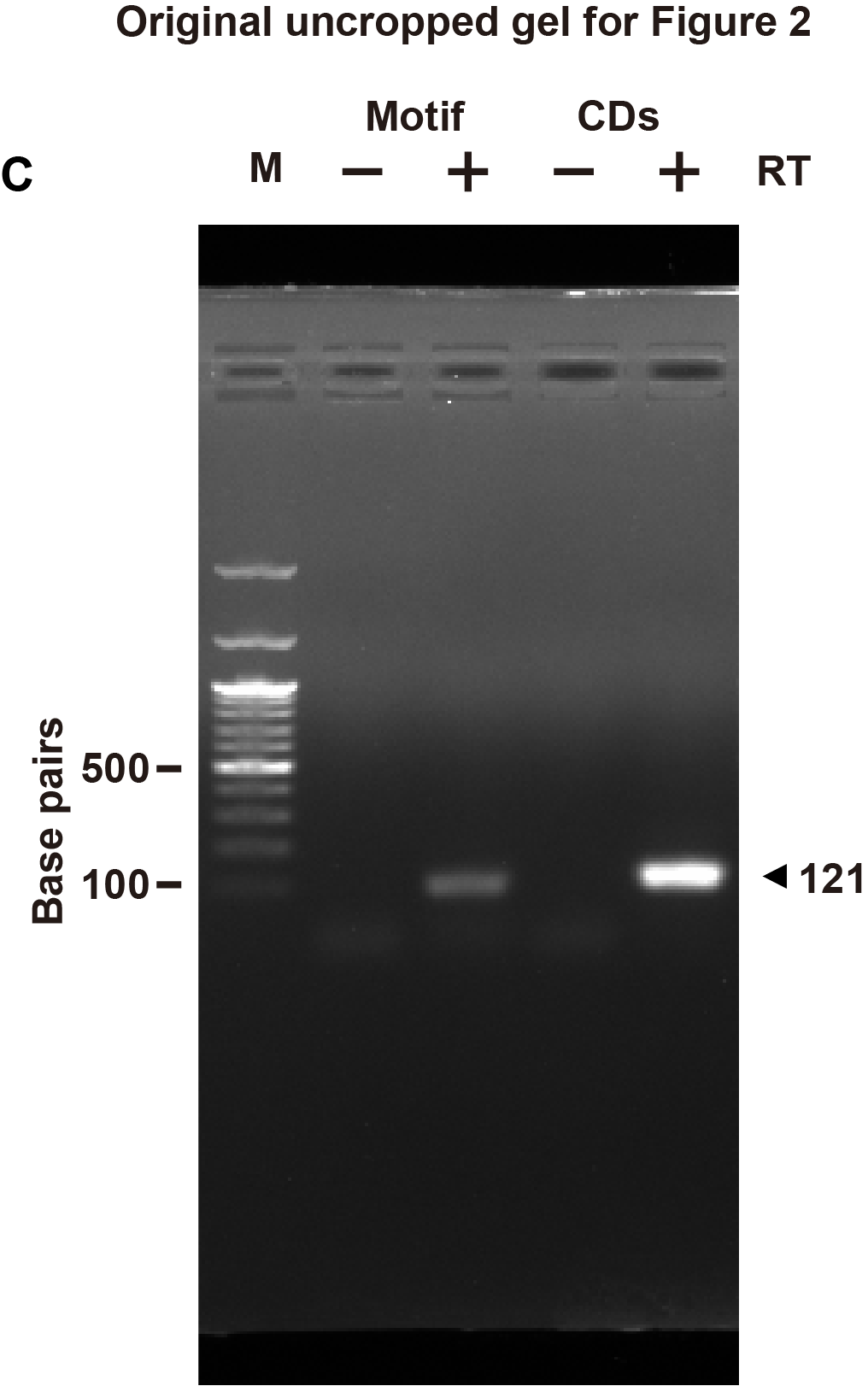

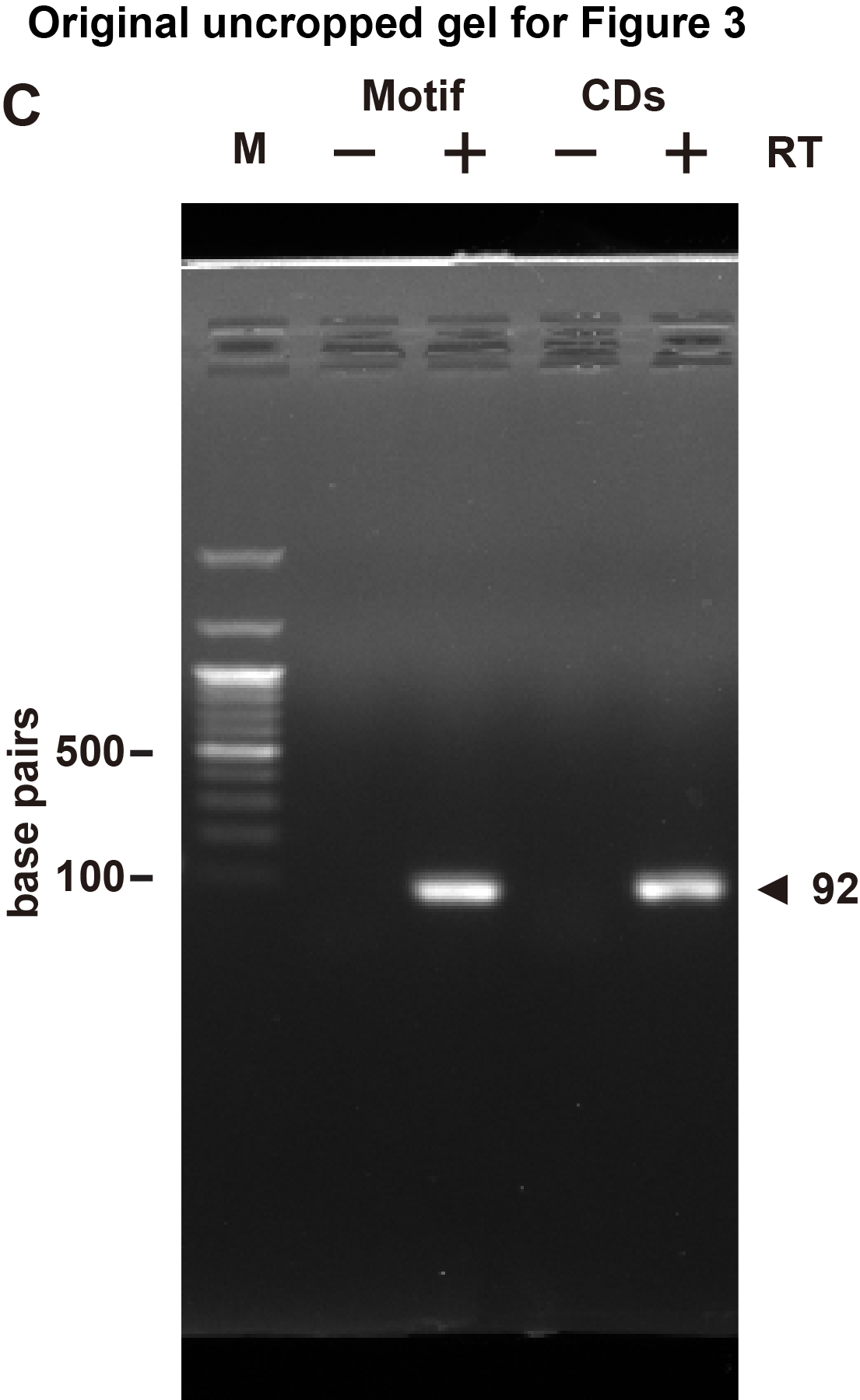

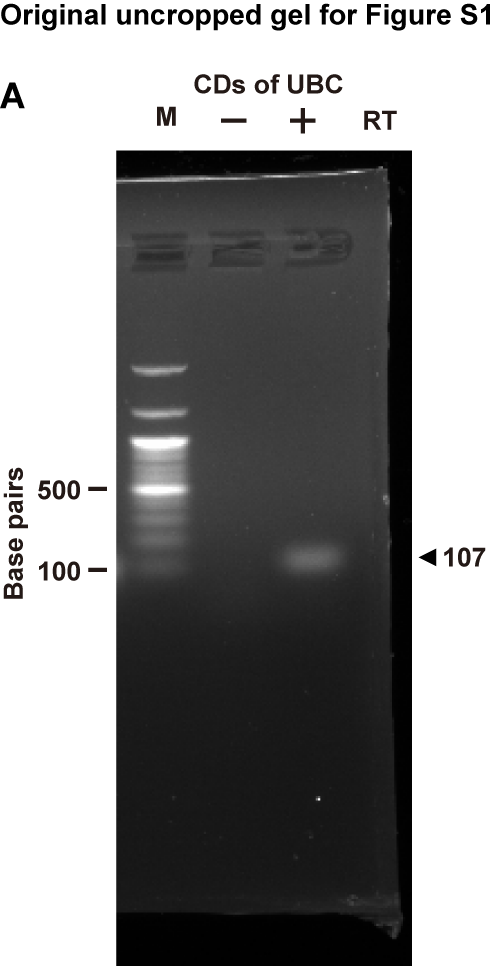
